# Combinatorial engineering for photoautotrophic production of recombinant products from the green microalga *Chlamydomonas reinhardtii*

**DOI:** 10.1101/2022.02.28.482248

**Authors:** Malak N. Abdallah, Gordon B. Wellman, Sebastian Overmans, Kyle J. Lauersen

## Abstract

*Chlamydomonas reinhardtii* has emerged as a powerful green cell factory for metabolic engineering of sustainable products created from the photosynthetic lifestyle of this microalga. Advances in nuclear genome and transgene expression engineering are allowing robust engineering strategies to be demonstrated in this host. However, commonly used lab strains are not equipped with features to enable their broader implementation in non-sterile conditions and high-cell density concepts. Here, we use combinatorial chloroplast and nuclear genome engineering to complement the *C. reinhardtii* strain UVM4 with publicly available genetic tools to enable the use of inorganic phosphite and nitrate as a sole the source phosphorous and nitrogen, respectively. We present recipes to create phosphite-buffered media solutions that enable high cell density algal cultivation. We then combine previously reported engineering strategies to produce the heterologous sesquiterpenoid patchoulol to high titers from our engineered green cell factories and show these products are possible to produce in under non-sterile conditions. Our work presents a straightforward means to generate *C. reinhardtii* strains for broader application in bio-processes for the sustainable generation of products.

## 1. Introduction

The model green microalga *Chlamydomonas reinhardtii* has emerged in recent years as a newcomer in the metabolic engineering space due to enabling advances in transgene design (Baier et al., 2018b, 2020) and the use of nuclear mutants with enhanced transgene expression rates (Neupert et al., 2009). *C. reinhardtii* has been extensively used as a host for chloroplast genome engineering for the expression of recombinant proteins for several years (Wannathong et al., 2016; Dyo and Purton, 2018). However, this alga has historically demonstrated recalcitrance to nuclear transgene expression owing to genetic architectures with high guanine-cytosine (GC) nucleotide content and intron density, as well as a recently characterized epigenetic silencing mechanism (Neupert et al., 2020). Through a series of mutational events, strains UVM4 and UVM11 were generated which exhibited improvements in transgene expression over others (Neupert et al., 2009; Barahimipour et al., 2016). UVM4 has become a workhorse strain for demonstrations of efficient transgene expression, with examples of heterologous production of sesquiterpenes (Lauersen et al., 2016; Wichmann et al., 2018), diterpenes (Lauersen et al., 2018; Einhaus et al., 2021), and polyamines (Freudenberg et al., 2021), modified fatty acid and alkene contents (Yunus et al., 2018), secreted recombinant proteins (Lauersen et al., 2013b, 2013a, 2015a; Baier et al., 2018a), and altered pigment composition (Perozeni et al., 2020). The reduced epigenetic silencing of this strain coupled with improvements of synthetic intron-addition transgene design strategies (Baier et al., 2018b, 2020) and optimized regulatory element combinations (Scranton et al., 2016; Einhaus et al., 2021), have resulted in an increased momentum for algal synthetic biology and green biotechnology applications with these hosts (Lauersen, 2019). *C. reinhardtii* represents a model green microalga that has very well developed molecular toolkits, including optimized (Lauersen et al., 2015b; Wichmann et al., 2018) and modular cloning (MoClo) plasmids (Crozet et al., 2018). Although a great deal of advancement has been made in gene expression and genetic engineering design, major limitations to scalable cultivation of *C. reinhardtii* remain which limits the broader development of engineered algal bio-processes.

Cultivation of *C. reinhardtii* is conducted at neutral pH, which means that the protein-rich algal cells are subject to rapid contamination/predation in non-sterile conditions. Sterility is difficult to maintain in large-scale cultivation concepts or in complicated bio-processes. Other industrially cultivated algae have features such as extreme pH or salinity tolerance which allow cultivation in selective conditions, or are dominant fast-growing, aggressive species. Although fast-growing, the UVM4 strain is also nitrate incapable. The use of nitrate is common in larger-scale algal cultivation media, as this nitrogen source does not cause significant pH shifts during its consumption. Complementation of nitrate metabolism is also important for increasing cell densities in cultivations, as has been recently demonstrated for the production of polyamines from this host (Freudenberg et al., 2021). The risk of contamination in addition to lack of nitrate metabolism capabilities make UVM4 deficient in features which would enable its broader use as an engineered algal green-cell factory outside of proof-of-principle laboratory experiments. One strategy for reducing contamination of algal cultures is the introduction of metabolic capacity for metabolism of inorganic phosphite as a phosphorous source. This has been shown in numerous organisms, including *C. reinhardtii*, to reduce contamination and act as a selection agent (López-Arredondo and Herrera-Estrella, 2012; Loera-Quezada et al., 2016; Changko et al., 2020; Cutolo et al., 2020; Dahlin and Guarnieri, 2022). Engineered expression of *Pseudomonas stutzeri* WM88 phosphite NAD+ oxidoreductase *ptxD* from either the chloroplast or nuclear genomes of algae has been shown to confer the ability to metabolize phosphite (López-Arredondo and Herrera-Estrella, 2012; Changko et al., 2020; Cutolo et al., 2020; Dahlin and Guarnieri, 2022).

To date, demonstrated advances in nuclear transgene expression for metabolic engineering described above have not incorporated combinatorial engineering with chloroplast expression constructs in the same strain. Here, we combined published advances in chloroplast engineering (Changko et al., 2020) with multiple nuclear engineering steps in UVM4 to demonstrate growth of this strain in phosphite- and nitrate-containing media while producing a proof-of-concept heterologous sesquiterpenoid. We present recipes to enable high-cell density cultivation in phosphite-buffered media and show that heterologous metabolites can be generated in the presence of microbial contamination. Our work shows that advances in nuclear and chloroplast engineering can be combined to yield more powerful green cell chassis that are amenable to bio-process goals.

## 2. Materials and Methods

### 2.1. Algal cultivation and growth measurements

*Chlamydomonas reinhardtii* strain UVM4 was used as the parental strain for transformations. This strain was derived from several rounds of mutation in the parental CC-4350 by Dr. Juliane Neupert in the lab of Prof. Dr. Ralph Bock (Neupert et al., 2009) and contains a mutation in Sir2-type histone deacetylase (SRTA) which enables improved transgene expression rates from the algal nuclear genome (Neupert et al., 2020). UVM4 is not able to use nitrate as a nitrogen source due to *nit1/nit2* locus mutations (Freudenberg et al., 2021). Microalgal cultures were routinely maintained in Tris acetate phosphate (TAP) medium (Gorman and Levine, 1965) with updated trace element solution (Kropat et al., 2011) and maintained under 150 μmol m^-2^ s^-1^ mixed cold and warm LED lights with 120-190 rpm agitation in shake flasks or microtiter plates. Light intensities and spectra were measured with a handheld spectrometer (Spectromaster C-7000, Sekonic). Ammonium in TAP salts solution was replaced with equimolar NaNO_3_ to make TAP-NO_3_. Replacements of phosphate with phosphite to make TAPhi and TAPhi-NO_3_ media are described in Supplemental File 1.

High-density 6xP medium was prepared as described in (Freudenberg et al., 2021) and buffered phosphite solutions to match molar concentrations of phosphorous to make 6xPhi medium are as described in Supplemental File 1. All phosphite solutions were filter sterilized and added to media after autoclaving. Cultivation in CellDeg HD100 cultivators (CellDeg GmbH, Germany) was performed with the indicated light and CO_2_ regimes by the growth control unit using either 6xP or 6xPhi media. Illumination was delivered by a Valoya broad spectrum LED board supplied by CellDeg GmbH (Germany, spectrum presented in Supplemental File 2).

Growth of algae and contaminants was analyzed by flow cytometry using an Invitrogen Attune NxT flow cytometer (Thermo Fisher Scientific, UK) equipped with a 488 nm blue laser for forward scatter and side scatter measurements, and a 695/40 nm filter to detect chlorophyll and non-fluorescent particles, respectively. All culture samples were diluted 1/100 with 0.9% NaCl solution and measured in technical triplicates using previously described settings (Overmans and Lauersen, 2022).

### 2.2. Plasmids, algal transformation, and screening for phosphite and nitrate metabolism

Plasmids used in this study are listed in Supplemental Table 1. All cloning and plasmid linearization was performed with Thermofisher FastDigest restriction enzymes, New England Biolabs Quick Ligase, and Q5 polymerase following manufacturer’s protocols. Plasmids were maintained in chemically competent *Escherichia coli* DH5a transformed by heat shock. Glass bead transformation of *C. reinhardtii* was performed as previously described for both chloroplast and nuclear targeted genetic constructs (Kindle, 1990; Kindle et al., 1991). Chloroplast transformation of the pPO_3_ plasmid ((Changko et al., 2020) graciously provided by Prof. Saul Purton) was performed with 10 μg circular DNA and 0.1 mm diameter glass beads rather than 0.424-0.600 mm as commonly used for nuclear transformation. Recovery was performed in 45 mL TAPhi liquid for 5 d with 150 μE PAR prior to plating. Selection was achieved by plating on TAPhi agar plates incubated at 200 μE for 2-3 weeks. Colonies were then grown in TAPhi liquid in microtiter plates until green.

One transformant with clear growth in liquid TAPhi, hereafter named UVM4-phi, was transformed for complementation of nitrate metabolic capacity by co-transformation of linearized pMN24 (Fernández et al., 1989b) and pMN68 (Schnell and Lefebvre, 1993) (Chlamydomonas Resource Center, https://www.chlamycollection.org) by glass beads as previously described (Freudenberg et al., 2021) with overnight recovery and subsequent selection on TAPhi-NO_3_ agar plates. Transformant colonies recovered on TAPhi-NO_3_ plates were then compared in liquid media in 24-well microtiter plates with standard lighting conditions at 180 rpm. Homoplasmy of the pPO3 integration into the chloroplast genome was determined by PCR using primers Fw: AATTGTATGGGCTCACAACAAACTTAAAGT and Rv: TAAAATTGTGAGACCATGAGTAATGTTCCTCC. The resulting transformants were also screened by iodine vapour assay as previously described (Wichmann et al., 2018) to determine if random integration had caused starch synthesis modifications. Modified UVM4 transformants which grew with phosphite and nitrate are referred to as UVM4-Phosphite-Nitrate (UPN) strains.

Efficiency of nuclear transgene expression of intermediate strains was investigated by glass bead transformation of the pOpt2_mVenus_Paro plasmid (Wichmann et al., 2018) followed by selection on each respective modified medium with 10 mg L^-1^ paromomycin and fluorescent reporter expression analysis. Fluorescent mVenus expression intensities were analyzed by picking primary transformant colonies using a PIXL robot (Singer Instruments, UK) to 384 colonies/plate layout on manufacturer supplied rectangular Petri dishes. After one week, colonies were replicated using the Singer Instruments ROTOR to generate imaging-ready colonies. White-light pictures of algae colony plates were taken in the built-in PIXL camera. Chlorophyll and mVenus fluorescence signals were captured in an Analytik Jena Chemstudio Plus gel doc with eLite halogen light source and excitation filters. Chlorophyll fluorescence was captured by 475/20 nm excitation with orange DNA gel emission filter with 1 sec exposure, while mVenus signal was captured with 510/10 nm excitation and 530/10 nm emission filter with 30 sec exposure.

### 2.3. Generation of patchoulol producing UPN transformants

Plasmids for algal nuclear genome-based expression of the *Pogostemon cablin* patchoulol synthase (*Pc*PS, UniProt Q49SP3) were adapted from (Lauersen et al., 2016). The gene expression cassette for *Pc*PS expression was modified from the pOpt2 vector concept of (Wichmann et al., 2018) to contain transgene designs presented in (Einhaus et al., 2021; Freudenberg et al., 2021). Briefly, *Pc*PS expression here was driven by the hybrid heat shock 70A-beta-bubulin promoter described by (Einhaus et al., 2021) and the mVenus cassette was modified to contain 2 copies of the *C. reinhardtii* ribulose-1,5-bisphosphate carboxylase/oxygenase small subunit (RBCS2) intron 1. The RBCS2 intron 2 was moved into the C-terminal strep-II tag of the gene-of-interest expression cassette in the pOpt2_mVenus_Paro plasmid which confers paromomycin resistance in *C. reinhardtii* (Wichmann et al., 2018). *Pc*PS was subcloned into *Bam*HI-*Bgl*II, and 2X, 3X, and 4X *Pc*PS expression cassettes were built by *Sca*I-*Bgl*II inserts from the previous plasmid subcloned into *Sca*I-*Bam*HI of the progenitor plasmid described in Supplemental Figure 1. All constructs contain the C-terminal mVenus (YFP) fusion which enabled plate-level fluorescence detection in UPN colonies picked by the PIXL robot. *C. reinhardtii* squalene synthase (UniProt A8IE29) knockdown was achieved by secondary transformation using the previously described pOpt2_*c*CA-*g*Luc_i3-SQS_Spect plasmid (Wichmann et al., 2018). UPN *Pc*PS-YFP + SQS k.d. double transformants were selected on TAPhi-NO_3_ agar media containing 10 mg L^-1^ paromomycin and 200 mg L^-1^ spectinomycin as previously described (Wichmann et al., 2018). YFP and luciferase signals of UPN colonies were captured in the Chemstudio PLUS with previously described buffers and reagents for *Gaussia princeps* luciferase bioluminescence analysis (Lauersen et al., 2013a). Full-length target recombinant protein was determined by SDS PAGE and in-gel fluorescence of whole cell pellets in the Chemstudio PLUS. All plasmid sequence files used in this work are given in Supplemental File 3.

#### 2.3.1 Gas Chromatography analysis of patchoulol productivity

UPN transformants expressing *Pc*PS variants were screened for heterologous patchoulol productivity by cultivation in 4.5 mL TAPhi-NO_3_ media with 500 μl dodecane overlay in triplicates for 6 d as previously described (Lauersen et al., 2018). Six individual transformants were investigated for each plasmid construct or combination after fluorescence, or fluorescence and luciferase, screening at the agar-plate level. Dodecane samples were collected from cultures, 90 μl of each collected dodecane sample was transferred into triplicate GC vials. A patchoulol standard (18450, Cayman Chemical Company, USA) calibration curve in the range 10-200 μM patchoulol in dodecane was used for linear-range quantification. 250 μM of alpha-humulene (CRM40921, Sigma-Aldrich, USA) was added as an internal standard to each dodecane sample and patchoulol standard. Quantification methods and calculations are shown in Supplemental File 4. The dodecane samples were analyzed using an Agilent 7890A gas chromatograph (GC) equipped with a DB-5MS column (Agilent J&W, USA) attached to a 5975C mass spectrometer (MS) with triple-axis detector (Agilent Technologies, USA). A previously described GC oven temperature protocol was used (Overmans and Lauersen, 2022). All GC-MS measurements were performed in triplicate (n=3), and chromatograms were manually reviewed for quality control. Gas chromatograms were evaluated with MassHunter Workstation software version B.08.00 (Agilent Technologies, USA).

## 2.4. Test of intentional contamination in cultures

To test the ability of engineered *C. reinhardtii* UPN lines to withstand contamination in non-sterile conditions using media modifications presented in this work, cultivation was performed with intentional yeast contamination. TAP-NO_3_, TAPhi-NO_3_, 6xP, and 6xPhi media were used to cultivate an engineered SQS k.d. + 2X*Pc*Ps expressing UPN transformant. *Saccharomyces cerevisiae* (yeast) cells were cultured in Yeast Extract-Peptone-Dextrose (YPD) medium (Cold Spring Harbor Protocols) overnight at 28°C. The following day, pelleted cells were resuspended in 300 mL 6xP or 6xPhi media. Triplicate wells in 6-well microtiter plates containing 4 mL dilute UPN patchoulol culture in 6xP or 6xPhi media were inoculated with either 500 μl of yeast-solutions (above) or clean medium as controls. 500 μl of n-dodecane overlay was also added to each well. Approximately 3mL of concentrated potassium bicarbonate buffer was added between the wells to provide a dilute CO_2_ atmosphere as previously described (Dienst et al., 2020). Cultures in TAP-media were grown for 6 d and 6xP/Phi for 9 d on laboratory shakers at 120 rpm with 12h:12h light:dark cycle (150 μE). Cell densities and patchoulol productivities were analyzed as described above.

## 3. Results

### 3.1. Phosphite and nitrate metabolism can be combined in the nuclear mutant UVM4

The *C. reinhardtii* mutants UVM4 and UVM11(Neupert et al., 2009) exhibit reduced transgene silencing due to a mutation in the in Sir2-type histone deacetylase (SRTA) (Neupert et al., 2020). UVM4 has served as a powerful parent strain for many recent examples of metabolic engineering in this green microalga (Lauersen et al., 2016, 2018; Wichmann et al., 2018; Einhaus et al., 2021; Freudenberg et al., 2021). Despite its value for past experiments, the alga is grown at neutral pH and contains mutations in its nitrate metabolism which prevent use of this nitrogen source. These two features manifest in high-risk of microbial contamination and the inability to use industrially relevant culture media (Changko et al., 2020; Freudenberg et al., 2021). To prepare UVM4 strains for broader applications, we set to complement it with the capacity to use phosphite as a P source and nitrate as an N source.

We complemented UVM4 with plasmid pPO3 (Changko et al., 2020), to express the *P. stutzeri* WM88 phosphite NAD+ oxidoreductase *ptxD* (López-Arredondo and Herrera-Estrella, 2012) that converts inorganic phosphite into organic phosphate from the algal chloroplast genome (Figure 1A). We found it was possible to transform UVM4 with this chloroplast genome-integrating plasmid by glass bead transformation and select colonies on TAPhi medium with no additional selection pressure. A resulting UVM4-phi transformant was then subsequently transformed with pMN24 and pMN68 plasmids which contain genomic copies of the *nit1* and *nit2* loci respectively to complement nitrate metabolism capacity (Fernández et al., 1989a; Schnell and Lefebvre, 1993) (Figure 1A).

**Figure 1.**
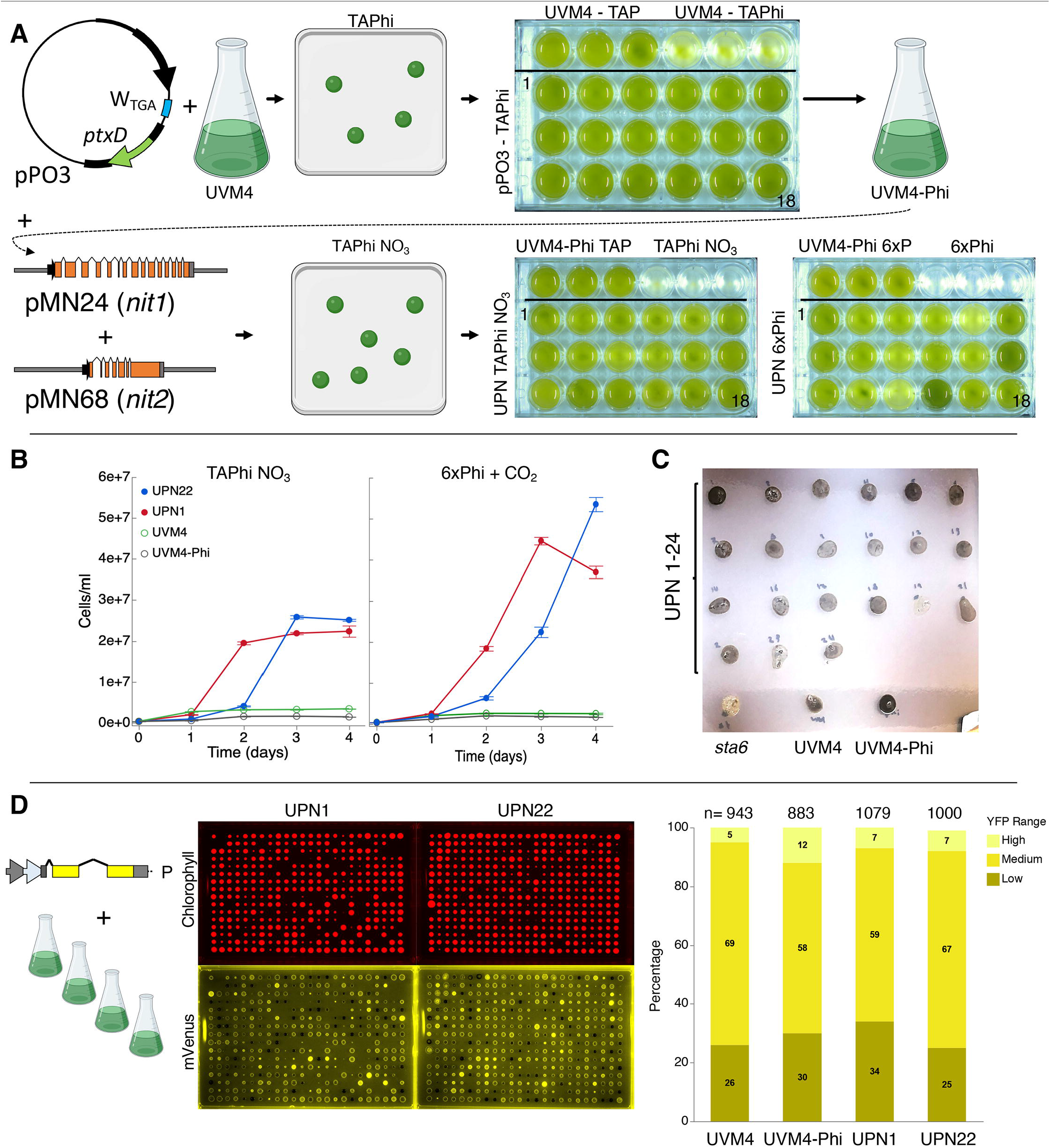
Complementation of C. reinhardtii strain UVM4 for growth on phosphite and nitrate. **A** Plasmid pPO3 (Changko et al., 2020) was transformed into UVM4 and colonies were recovered on TAPhi agar medium. Colonies were then cultivated in TAPhi liquid medium and one strain (UVM4-Phi) was selected for further complementation with pMN24(Fernández et al., 1989b) and pMN68 (Schnell and Lefebvre, 1993) (*nit1*/*nit2*) plasmids. Selection was performed on TAPhi-NO_3_ plates and resultant colonies capable of growth on phosphite and nitrate (UPN) were cultivated in liquid mixotrophic (TAPhi-NO_3_) and autotrophic (6xPhi) media. Parental strains were grown as reference in each previous stage media as shown. **B** Growth curves of liquid cultures of selected colonies which performed well in TAPhi-NO_3_ media in TAPhi-NO_3_ or 6xPhi media. Parental strains UVM4 and UVM4-phi, were not able to proliferate in the nitrate phosphite containing media. **C** UPN strains were investigated by iodine vapour staining at the agar plate level to determine if the transformation of three plasmids above had caused background mutations in starch synthesis. Dark colour of colonies indicates presence of starch, yellow or light colour indicates perturbed starch metabolism as shown for the starchless *sta6* (Zabawinski et al., 2001) mutant. Lighter starch staining is observed in UPN strains 19 and 23. D Strains UVM4, UVM4-Phi, UPN1 and UPN22 were transformed with the pOpt2_mVenus_Paro plasmid (cartoon) conferring paromomycin resistance and expressing the mVenus (YFP reporter). High-throughput robotic colony picking and fluorescence imaging was used to benchmark YFP expression across the transfomant population. Chlorophyll fluorescence (red) was used to identify true colonies, and YFP fluorescence (yellow) was graded for intensity of signal and plotted comparing numbers of high, medium, and low or no expression (right). Individual colonies analyzed for each transformation event summed from several plates are indicated for each strain. *C. reinhardtii* genetic elements: A – HSP70A promoter, R – RBCS2 promoter, i1 – RBCS2 intron 1, i2, RBCS2 intron 2, ß – beta tubulin promoter and its 5’ untranslated region (UTR), 3’UTR – RBCS2 3’ UTR

Colonies were recovered by selection on TAPhi-NO_3_ plates with no antibiotic (Figure 1A). Complementation with *nit1*/*2* can sometimes lead to colonies that survive on agar plate, but do not perform well in liquid medium. Therefore, we also benchmarked performance of 24 UVM4-phosphite-nitrate (UPN) colonies derived from these transformations in TAPhi-NO_3_ and photoautotrophic cultivation with CO_2_ as a carbon source (Figure 1A, lower right). Parental strains were not able to grow in the selective media for each plasmid. UPN strains grown in liquid medium exhibited variable performance, especially in photoautotrophic conditions (Figure 1A,B). Most colonies maintained normal starch accumulation, which was qualitatively assessed by iodine vapour, however, colonies 19 and 23 showed reduced iodine staining (Figure 1C). Homoplasmy of pPO3 integration was determined in UVM4-phi and nitrate complemented strains (Supplemental Figure 2).

To confirm that the three plasmid integrations did not modify the performance of the parent UVM4 capacities for nuclear transgene expression, several UPN strains with acceptable growth in liquid phosphite-nitrate media were transformed with a YFP reporter plasmid(Wichmann et al., 2018). High-throughput robotics assisted colony picking allowed analysis of between 943-1079 colonies for each strain and plate-level reporter imaging was used to quantify reporter expression across the populations. YFP reporter expression efficiencies in the final UPN strains were comparable to parent UVM4 (Figure 1D).

### 3.2. Phosphite can replace phosphate in buffered media for high cell-density cultivation of algal cells

Using mono- and di-basic forms of phosphite, we generated a buffered phosphite solution to emulate the phosphate buffer solution of recently published 6xP medium (Freudenberg et al., 2021) (Supplemental File 1). In order to compare whether our high-density phosphite medium (6xPhi) could be used in a comparable fashion to 6xP, we benchmarked growth of a UPN strain in a high-density cultivation concept using high-light and membrane delivered CO_2_ in CellDeg HD100 cultivators (Figure 2A). We did not observe differences in performance for the UPN strain cultivated in 6xP or 6xPhi which reached comparable cell densities throughout cultivation (Figure 2B).

**Figure 2.**
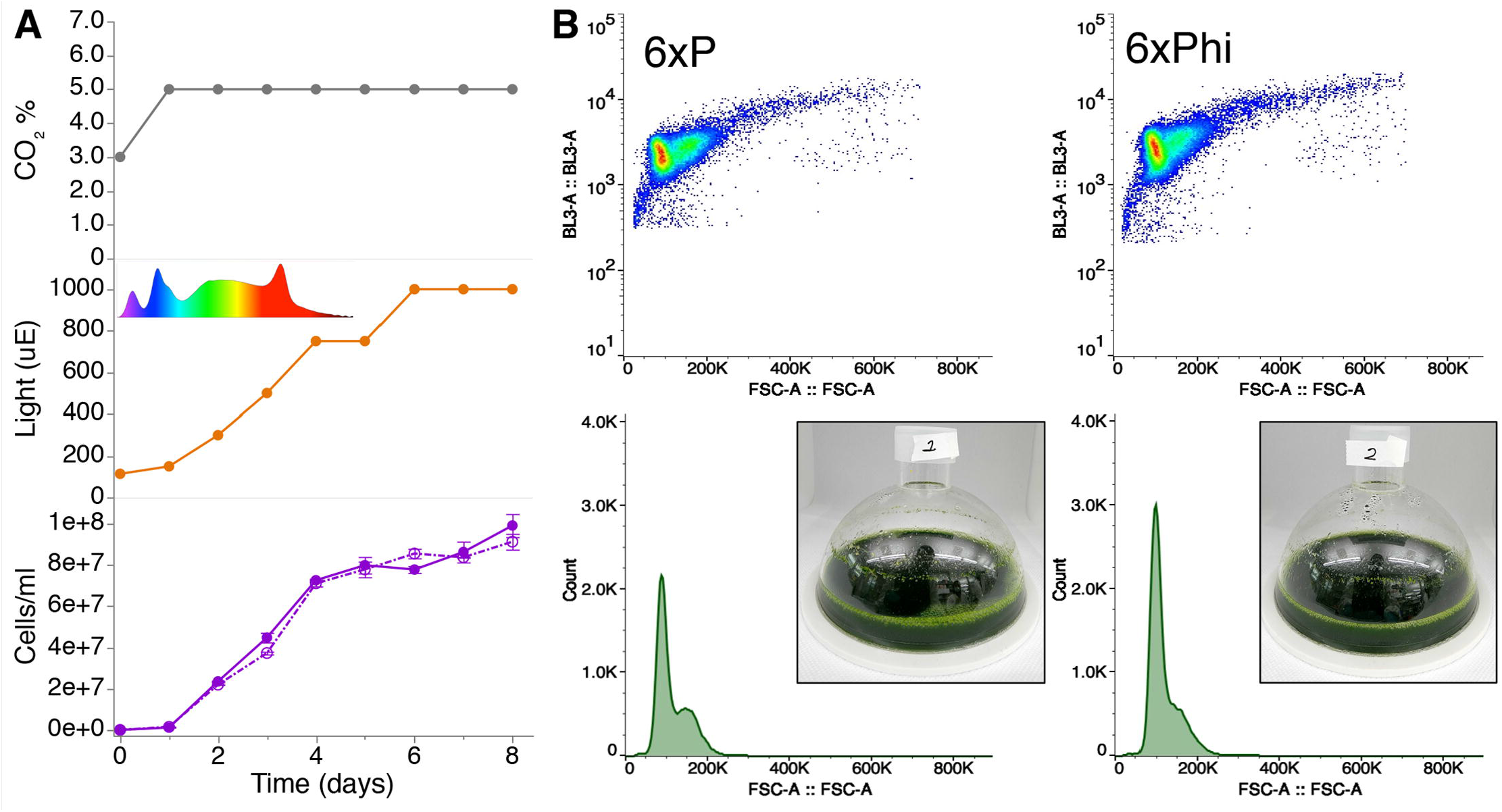
Buffered phosphite solutions can be used in algal high cell density medium concepts to replace phosphate. **A** Growth of strain UPN22 was tracked in cultivations in 100 mL of 6XP (solid line) or 6XPhi (dashed line) media in CellDeg HD100 cultivators following the CO_2_ and light regime indicated. Spectrum of the Valoya daylight lamp is shown. Cell densities were recorded daily. **B** Forward and backscatter plots from flow cytometry of samples from day 6 of each culture with photographs of the dense green culture in either medium.

### 3.3. Heterologous products can be efficiently made in UPN strains

We then chose to combine two proven engineering strategies for sesquiterpenoid production from a UPN strain grown only in TAPhi-NO_3_ medium. The *C. reinhardtii* codon optimized *P. cablin* patchoulol synthase (*Pc*PS) was expressed in 1, 2, 3, and 4X copy fusion protein constructs with C-terminal YFP from the nuclear genome of this alga (Figure 3). Robotics assisted colony picking and YFP screening allowed selection of six transformants per plasmid with confirmed *Pc*PS expression which were benchmarked for patchoulol productivity as previously described (Lauersen et al., 2018). The best performing transformants were subsequently transformed with a secreted luciferase-artificial-micro-RNA expression construct targeting the *C. reinhardtii* squalene synthase (SQS)(Wichmann et al., 2018). After colony recovery and robotics picking, plate-level imaging was used to isolate colonies with YFP fluorescence signal (*Pc*PS) and luciferase activity (SQS k.d.) (Figure 3). Patchoulol productivity analysis indicated striking improvements in patchoulol productivity for SQS k.d. strains compared to parentals with the best performing strains generating ~145 fg patchoulol cell^-1^ (Figure 3). Full-length fusion protein expression could be confirmed only for 1-3X *Pc*PS-YFP constructs by in gel fluorescence (Supplemental Figure 3).

**Figure 3:**
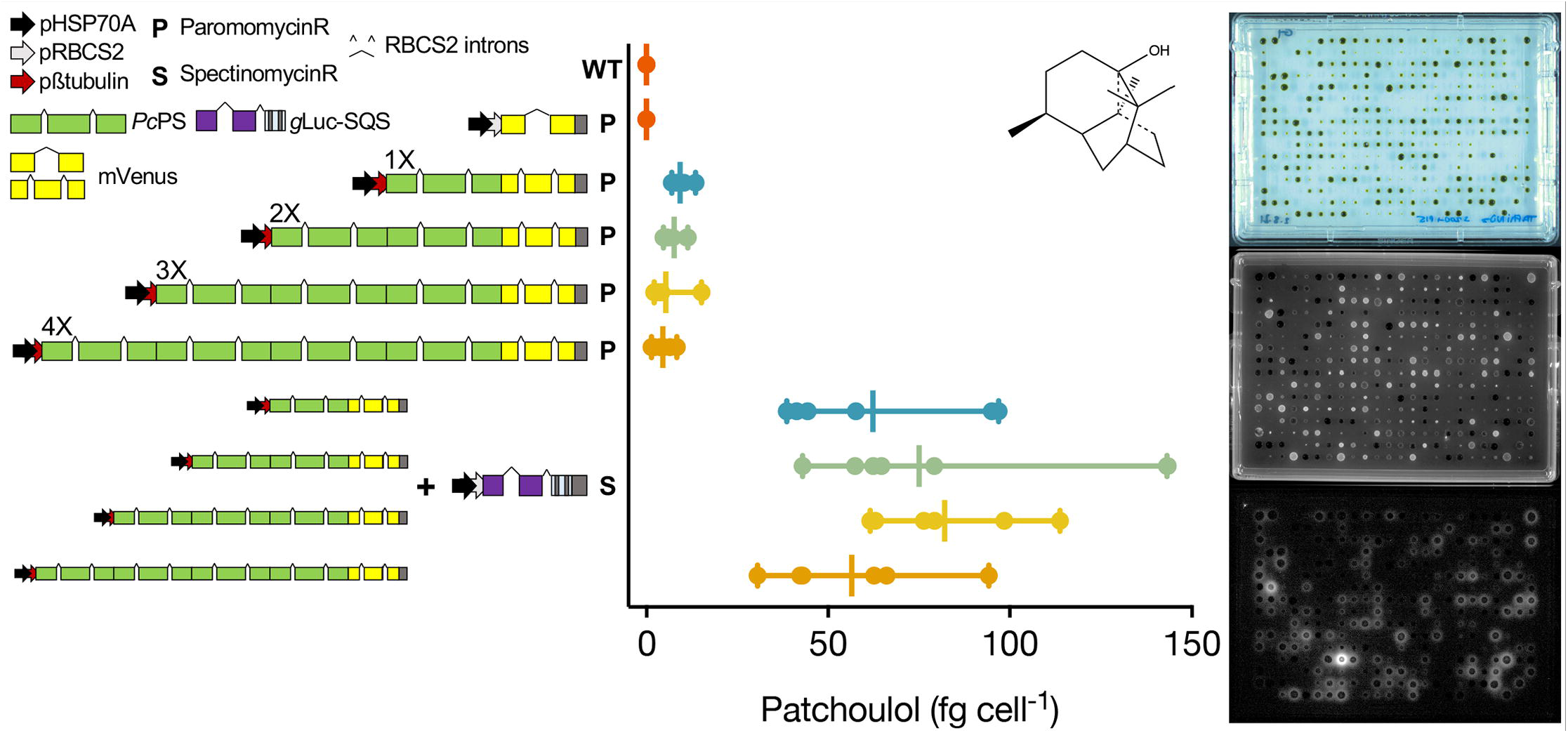
Genetic constructs used to generate heterologous patchoulol production from a UPN strain. Single, double, triple, and quadruple copies of the *C. reinhardtii* codon optimized, intron containing *P. cabiln* patchoulol synthase were fused to generate different expression plasmids with C-terminal mVenus (YFP) reporter fusions as previously described(Lauersen et al., 2016). Each plasmid was transformed into UPN and mVenus (YFP) expressing colonies were isolated for each construct and benchmarked for patchoulol production (n=6). The chemical structure of patchoulol is shown. The best performing individual from each plasmid was then subsequently transformed with a plasmid expressing a luciferase-amiRNA construct which downregulates the *C. reinhardtii* squalene synthase. Combined high-throughput fluorescence and luciferase screening of colonies led to isolation of strains with both constructs expressed (n=6) which were then subsequently benchmarked for patchoulol productivity.

### 3.4. Nitrate and phosphite can both assist contaminant control in algal cultures

As contamination of cultures can be an issue with neutral pH cultivation, we wanted to determine if phosphite and nitrate could permit algal growth in the presence of contamination. We intentionally contaminated the best performing UPN *Pc*PS SQS k.d. strain in mixotrophic (acetic acid) and photoautotrophic cultures in media with nitrate as a nitrogen source and either phosphate or phosphite as a phosphorous source. Yeast cells were added to cultures directly in higher cellular abundances than algal cells (Figure 4). In all media conditions, yeast cells did not proliferate, regardless of the presence of organic carbon, but also did not reduce in number. When acetic acid (TAP-derived) media were used, algal growth was reduced compared to cultivations without yeast, also with phosphite (Figure 4). No difference in performance was noted in photoautotrophic cultures. In all conditions, the presence of high concentrations of yeasts in cultivations did not inhibit heterologous patchoulol production (Figure 4).

**Figure 4.**
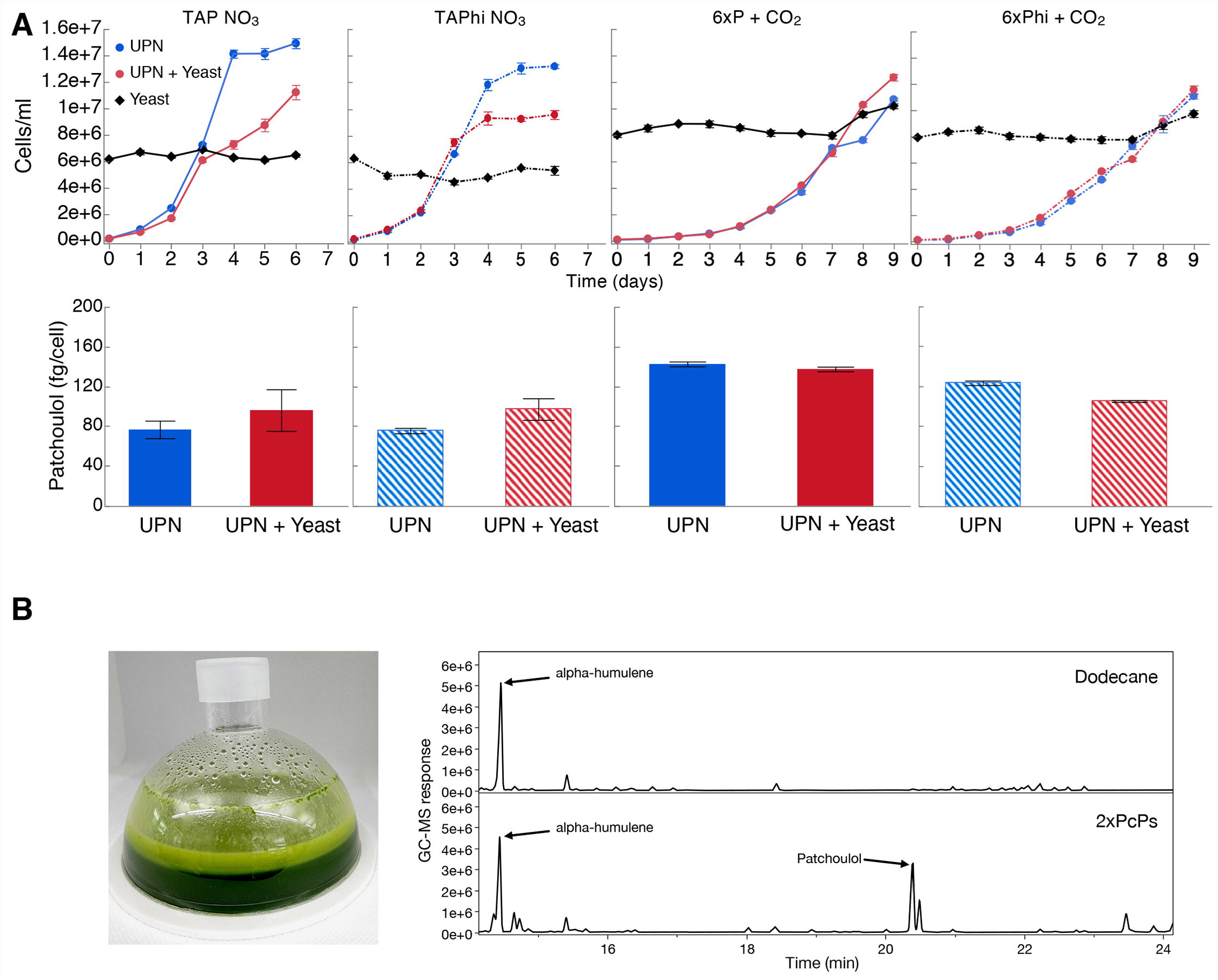
Production of heterologous sesquiterpenoid in the presence of contamination. **A** *C. reinhardtii* UPN 22 expressing 2X*Pc*PS-YFP + SQS-amiRNA was cultivated in different trophic modes with and without phosphite and intentionally contaminated with *S. cerevisiae* cells. TAP-NO_3_ and TAPhi-NO_3_ were used to compare mixotrophic conditions where acetic acid was a sole carbon source, while 6xP and 6xPhi were used to test photoautotrophic conditions. All growth curves with Phi are represented with dashed lines and hashed bars. Dodecane overlay was used to capture heterologous patchoulol produced. Yeast cells were intentionally inoculated at high densities to challenge the algal cells to outcompete them in these conditions. CO_2_ was delivered to autotrophic cultures by placing high-concentration bicarbonate buffer between microtiter plate wells as an inefficient delivery mechanism to further challenge the algal cells. **B** cultivation of this strain in 6xPhi medium in HD100 cultivator (pictured) with dodecane overlay resulted in efficient patchoulol production from CO_2_. Two GC-MS chromatograms are shown, one of dodecane blank with alpha-humulene internal standard and one from algal culture indicating the peak of produced patchoulol.

We then benchmarked patchoulol productivity in a 200 mL culture in a membrane gas delivery bioreactor containing 10% dodecane overlay to capture heterologous sesquiterpenoid product. Culture volume was adjusted to 200 mL to avoid contact of dodecane with the hydrophobic gas delivery membrane during shaking and the culture was operated in non-sterile conditions. The culture accumulated up to 6.5×10^7^ ± 1.9×10^6^ cells mL^-1^ and generated 6.2 mg L^-1^ patchoulol in 6 days using this system.

## 4. Discussion

### 4.1. Designing engineerable strains to be ready for bio-processes

We chose phosphite metabolism complementation with plasmid pPO3 as this also contains extra future chloroplast genome engineering potential through the addition of the W_TGA_ tRNA for tryptophan as previously described (Changko et al., 2020). When filter sterilization was used, transformation and selection on phosphite solutions was greatly improved and appearance of background algal growth at the plate level was reduced (data not shown). After confirmed growth of pPO3 transformants in liquid phosphite, nitrate metabolism was complemented by transformation of both pMN24 (*NIT1*) and pMN68 (*NIT2*) plasmids in UVM4. This double transformation is relatively inefficient, nevertheless, we could generate several dozen colonies per transformation which recovered on nitrate plates. Colonies which recovered on nitrate plates did, however, not all perform well in liquid culture growth in nitrate-containing liquid media. We chose to move forward with only those colonies which appeared to grow to dark green stationary phase (Figure 1A). Colonies were then checked for homoplasmy integration of the pPO3 phosphite conferring plasmid (Supplemental Figure 2) and two were benchmarked for their growth in phosphite- and nitrate-containing media compared to their parental strains (Figure 1B).

To determine if our strategy for UVM4 augmentation would allow future engineering to benefit from these metabolic enhancements, two questions remained: 1.) was nuclear transformation expression efficiency disturbed during these events in UVM4 derivatives? 2.) Can inorganic phosphite be used in a similar way to organic phosphate, for buffered media solutions? We benchmarked two fully complemented UPN transformants, their UVM4-phi parent, and the UVM4 starting strain for YFP efficiency expression from the nuclear genome. Using high-throughput colony picking, we were able to analyze ~1000 colonies per transformation event and compare YFP expression efficiencies across the populations (Figure 1C). Although some variance, there was little difference could be seen in the total ratio of high and mid-range YFP expressing colonies, suggesting our three plasmid integrations had not modified this capacity.

We then set out to make a buffered phosphite solution which could replace buffered phosphate solutions in culture media (Supplemental File 1). In direct comparisons of growth tests, the UPN strain did not show performance differences in phosphite compared to phosphate in photoautotrophic high-density cultivations (Figure 2). Our results indicate this is an effective strategy to use inorganic phosphite as a media component. As phosphate is a globally dwindling resource important to agriculture, bio-conversion of phosphite into phosphate may also enable the use of this waste mineral to yield bio-fertilizers through algal cultivation.

### 4.2. Patchoulol production in metabolically complemented strains

UPN strains were maintained exclusively on TAPhi-NO_3_ medium for all routine lab work. A further aim was to determine if it was possible to conduct additional metabolic engineering in these strains for heterologous isoprenoid production using Phi-NO_3_ media. Plasmids were constructed based on previous designs to express the patchoulol synthase (*Pc*PS) and localize it in the cytoplasm of the alga where this enzyme is known to convert freely available farnesyl pyrophosphate (FPP) into patchouli alcohol (patchoulol)(Lauersen et al., 2016). We chose to combine recently published modifications in promoter(Einhaus et al., 2021) and intron use(Freudenberg et al., 2021) (Supplemental Figure 1), in order to assist recombinant protein accumulation in an effort to enhance product yields. Previous work on *Pc*PS, indicated cellular patchoulol productivities could be enhanced when the protein was fused to itself in a repetitive fashion to yield more active sites per translated protein(Lauersen et al., 2016). Here, we copied this gene design strategy and combined it with an artificial micro-RNA (amiRNA) knockdown of the squalene synthase (SQS) (Figure 3). It was previously found that SQS k.d. improved (*E*)-α-biabolene titers from the cytoplasm of *C. reinhardtii* as this is the direct competitor for FPP precursor(Wichmann et al., 2018). Additive *Pc*PS units were found here to increase cellular patchoulol yields as previously observed (Lauersen et al., 2016). However, increasing repetitions past 3 *Pc*PS copies was found to be unstable and did not generate reliable patchoulol expression strains (Figure 3, Supplemental Figure 3). As expected based on past work with bisabolene(Wichmann et al., 2018), addition of the SQS k.d. to the best performing *Pc*PS variant of each plasmid lead to transformants with drastic improvements in patchoulol productivity. Previous engineering of patchoulol production for *C. reinhardtii* lead to a maximal volumetric productivity of ~350 μg patchoulol L^-1^ in mixotrophic 400 mL bioreactor conditions from a 3X*Pc*PS-YFP transformant(Lauersen et al., 2016). Here, a 2X*Pc*PS transformant subsequently transformed with the SQS k.d. generated lines producing 700-1400 μg patchoulol L^-1^ culture (Figure 3, Supplemental Figures 4 and 5). Apparent improvements in 1-4X *Pc*PS-YFP lines were observed by SQS k.d. and from 1-3X*Pc*PS-YFP, mean production increased with increasing *Pc*PS units from 1X-3X*Pc*PS. Maximal cellular productivity was observed in a single 2X*Pc*PS SQS k.d. line, with up to 143 fg patchoulol cell^-1^ (Figure 3).

A risk to scaled cultivation of engineered *C. reinhardtii* in bio-production concepts is contamination and reduced productivities, which is especially true for cell-wall deficient strains that may be more readily outcompeted by contaminants. We intentionally contaminated mixotrophic and photoautotrophic media containing either NO_3_ or Phi and NO_3_ with yeast, and cultivated a UPN-patchoulol producing strain in these sub-optimal conditions (Figure 4A). We inoculated the 2X*Pc*PS-SQS k.d. UPN strain into media containing 6.0×10^6^ cells mL^-1^ yeast, the same cell density as reached in mid-log phase for the algal cells. We chose to provide CO_2_ using potassium bicarbonate buffers(Dienst et al., 2020) rather than direct gas delivery to further challenge the phototrophic cultures. In all conditions, yeast cells did not proliferate, with either acetic acid as a carbon source, or in the photoautotrophic media (Figure 4A). Mixotrophic cultures exhibited lower algal cell densities in later stages of cultivation than in the absence of yeast, likely due to the yeast sequestering some of the acetic acid. However, in the establishment phase (days 0-3), growth with yeast was not markedly different than that of cultures without yeast. In photoautotrophic cultures, yeast was inoculated at higher starting cell densities (8×10^6^ cells mL^-1^) as there is no organic carbon source. This density was chosen to determine if the yeast cells would competitively inhibit the inoculated algal cells. Here, the UPN strain was able to overtake the yeast cells, demonstrating linear growth in both conditions relative to carbon diffusion rates in the medium. Under all conditions, the presence of yeast contaminants did not hinder the accumulation of heterologous patchoulol in dodecane overlays which exhibited similar productivities per algal cell (Figure 4A). Our results indicate that both nitrate and phosphite are a powerful combination to limit contaminating microbial competitors in engineered algal cultivation concepts.

To determine if we could produce patchoulol in non-sterile conditions, we cultivated this strain in a CellDEG HD100 bioreactor with dodecane overlay using 6xPhi medium (Figure 4B). Dodecane impairs the hydrophobic gas delivery membrane of the reactors, so we used 200 mL culture volume to prevent the solvent interacting with the membrane. Previous photoautotrophic yields of this product were only 350 μg L^-1^ in 8 days (Lauersen et al., 2016). Here, without any process optimization, ~6.2 mg patchoulol L^-1^ was produced from CO_2_ in 7 days (Figure 4). A previous study with *Synechocystis* sp. PCC 6803 used a similar membrane gas delivery system for 10 mL cultures to generate up to 17.3 mg patchoulol L^-1^ in 8 days using a two-stage semi batch mode where half of the culture medium was replaced after 96 hours. We did not further optimize our production experiments in the HD100, as the risk of dodecane-membrane wetting means cultivation must be performed with volumes not intended for the system. A further issue of the dodecane overlay in turbid algal cultures is the formation of emulsions with hydrophobic cellular components and the dodecane solvent (Lauersen, 2019). The surface interaction of culture and dodecane causes significant emulsion formation in this volume ratio (Figure 4B), which is less pronounced in smaller volume cultivation units. Nevertheless, our results indicated the combination of nitrate and phosphite metabolic capacities enable high-density cultivations of engineered strains to be performed with reduced risks of contamination without affecting process yields. To our knowledge, this is the first demonstration of combinatorial plastid and nuclear genome engineering in a green alga to generate a strain producing a heterologous product.

### 4.3. Conclusions

Here, we have demonstrated the complementation of the common UVM4 nuclear mutant with the genetic capacity for phosphite and nitrate metabolism. These modifications were possible without affecting the nuclear transgene expression abilities of UVM4 and enabled the engineering of heterologous isoprenoid production in strains capable of growth in contamination reducing media. We present a recipe for buffered phosphite solutions to replace those of phosphate in common *C. reinhardtii* media and show improved titers of patchoulol though combination of strategies known to improve flux to sesquiterpenoid products. Our work can be used as a guide for others to adapt phosphite-nitrate metabolism into their strains and may enhance the transition of lab-scale engineering to less-sterile production concepts.

## Supporting information

Supplemental File 1 - Media Recipes

Supplemental File 2 - Light Spectrum

Supplemental File 3 - Plasmid Sequence

Supplemental File 4 - GC Quantification data

Supplemental Table 1 - Plasmids used in this work

## Acknowledgements

Subcloning of *Pc*PS plasmid 1X was performed in the lab of Prof. Dr. Olaf Kruse by Dr. Julian Wichmann and Dr. Thomas Baier as part of an Institute for Innovation Transfer (IIT), Universität Bielefeld, project funded by Lauersen (KAUST). The authors are grateful to Saul Purton for providing plasmid pPO3 and Prof. Dr. Ralph Bock for graciously providing strain UVM4 through MTA between the Max Planck Institute of Molecular Physiology and KAUST. We would like to express thanks to SSB group members for cooperation and collaboration during this project. The research reported in this publication was supported by baseline funding from KAUST to KL.

## Supplemental figure legends

**Supplemental Figure 1:** Modifications to the pOpt2 plasmids used in this work. **Above**: Overview of plasmid architecture of the pOpt 2.0 vectors from Wichmann et al. (2018) 10.1016/j.ymben.2017.12.010. The gene of interest (GOI) reporter cassette is highlighted. **Middle**: in this work, we modified the GOI cassette to contain the optimized HSP70A, beta-tubulin (and 5’UTR) from Einhaus et al. (2021) 10.1021/acssynbio.0c00632. The mVenus reporter (NCBI: AAZ65844) was also modified to contain two copies of the RBCS2 intron 1, and the RBCS2 intron 2 was moved into the C-terminal StrepII tag, so that future C-terminal fusions could benefit from this orientation in a similar fashion to that presented in Freudenberg et al. (2021) 10.1016/j.biortech.2020.124542. All elements are from *C. reinhardtii:* A – HSP70A promoter, R – RBCS2 promoter, i1 – RBCS2 intron 1, i2, RBCS2 intron 2, ß – beta tubulin promoter and its 5’ untranslated region (UTR), 3’UTR - RBCS2 3’ UTR. AmpR – ampicillin resistance cassette of the pBluescript SK(+) backbone. **Below**: Plasmids for the expression of the *C. reinhardtii* codon optimized and synthetic intron-containing *Pogostemon cablin* patchoulol synthase (UniProt: Q49SP3, *Pc*PS). Modified pOpt2.0 expression cassettes were used to subclone the *Pc*PS which had been previously codon optimized (including intron spreading) for expression from the nuclear genome of *C. reinhardtii* (NCBI: KX097887, Lauersen et al. (2016) 10.1016/j.ymben.2016.07.013). All plasmids have the pOpt2.0 – paromomycin resistance cassette (P) 3’ of the GOI expression cassette pictured (as in Wichmann et al. (2018)). Subcloning of *Pc*PS plasmid 1X was performed in the lab of Prof. Dr. Olaf Kruse by Dr. Julian Wichmann and Dr. Thomas Baier as part of an Institute for Innovation Transfer (IIT) project funded by Lauersen (KAUST). Further subcloning of 2X, 3X, 4X were performed by Dr. Gordon Wellman in Lauersen’s lab at KAUST. Abbreviations: Y – mVenus expression cassette, 1X, 2X, 3X, 4X plasmids contain the respective numbers of *Pc*Ps copies. Cloning was achieved by using the previous plasmid, with *Sca*I-*Bam*HI as the receiving vector and *Sca*I-*Bgl*II as an insert to amplify the *Pc*PS coding sequence.

**Supplemental Figure 2:** Confirmation of pPO3 integration into the chloroplast genome of UVM4 in derivative strains UVM4-phi and UPN22. Primers Fw: AATTGTATGGGCTCACAACAAACTTAAAGT and Rv: TAAAATTGTGAGACCATGAGTAATGTTCCTCC were used to perform PCR on DNA extracts from each strain. The target region without amplification should yield 1050 bp band, while integration should yield 3075 bp products.

**Supplemental Figure 3:** In gel fluorescence of SDS PAGE samples from one representative mutant of each of the genetic constructs indicated. Fluorescence image was captured with 510/10 nm excitation and 530/10 nm emission filter in the Analytik Jena Chemstudio Plus with eLite. White contrast black and white image was taken without emission filter using 510/10 nm excitation to visualize the marker.

**Supplemental Figure 4:** Patchoulol productivities observed in dodecane overlays for 6 transformants selected for bright YFP fluorescence from each of the above plasmids and compared to parental UPN strain and empty vector (Y) generated control strains. Numbers at the bottom of the graph correspond to plasmid name and mutant number (1.1 = 1X *Pc*PS, transformant #1). Productivity-grouped averages are shown on the left and labelled with each plasmid name.

**Supplemental Figure 5:** Patchoulol productivities observed in dodecane overlays for six transformants isolated from secondary transformation of the best 1-4X *Pc*PS strains with *c*CA_*g*Luc_i3_SQSk.d. plasmid (Wichmann et al. 2018). **Upper right**: representitive GC-MS chromatograms showing the drastic increase of patchoulol production in SQS k.d. secondary transformants relative to alpha-humulene internal standard. **Lower graphs**: Numbers at the bottom of the graph correspond to plasmid name and mutant number (1×1 = 1X *Pc*PS SQS k.d. mutant 1). The averages of all transformants per group are shown on the left and labelled with each plasmid name. Corresponding parent performances are also shown from the data in previous figure.

## Tables

**Supplemental Table 1.** Genetic constructs used in this study. Plasmids for transformation of *C. reinhardtii* are shown as well as some of their respective properties. References to plasmid sequences are given and those generated in this work are provided in the supplement.

## 5. Contribution to the field statement

In this work, we combine previously published optimizations in algal metabolism in a mutant workhorse for nuclear transgene expression. We expand this strain with the metabolic capacity for use of inorganic phosphite as a sole phosphorous source, and nitrate as a nitrogen source. These combined engineering steps enable the cultivation of this strain in both high-density culture media, with loading of nitrates to increase cell densities, and the use of phosphite to mitigate contamination. We present recipes for replacing phosphate buffers with phosphite buffers and demonstrate heterologous production of the sesquiterpenoid patchoulol by combining synthase overexpression with competitive pathway knockdown. Our work is the first example of combinatorial chloroplast and nuclear genome engineering for heterologous metabolite engineering in a green algal host and sets an example for future engineering in these hosts to enable scalable cultivation concepts.

## Notes

### Competing Interest Statement

The authors have declared no competing interest.

