## Supplementary figures and images for "Combinatorial engineering for photoautotrophic production of recombinant products from the green microalga *Chlamydomonas reinhardtii*"

### Supplemental Figure 1

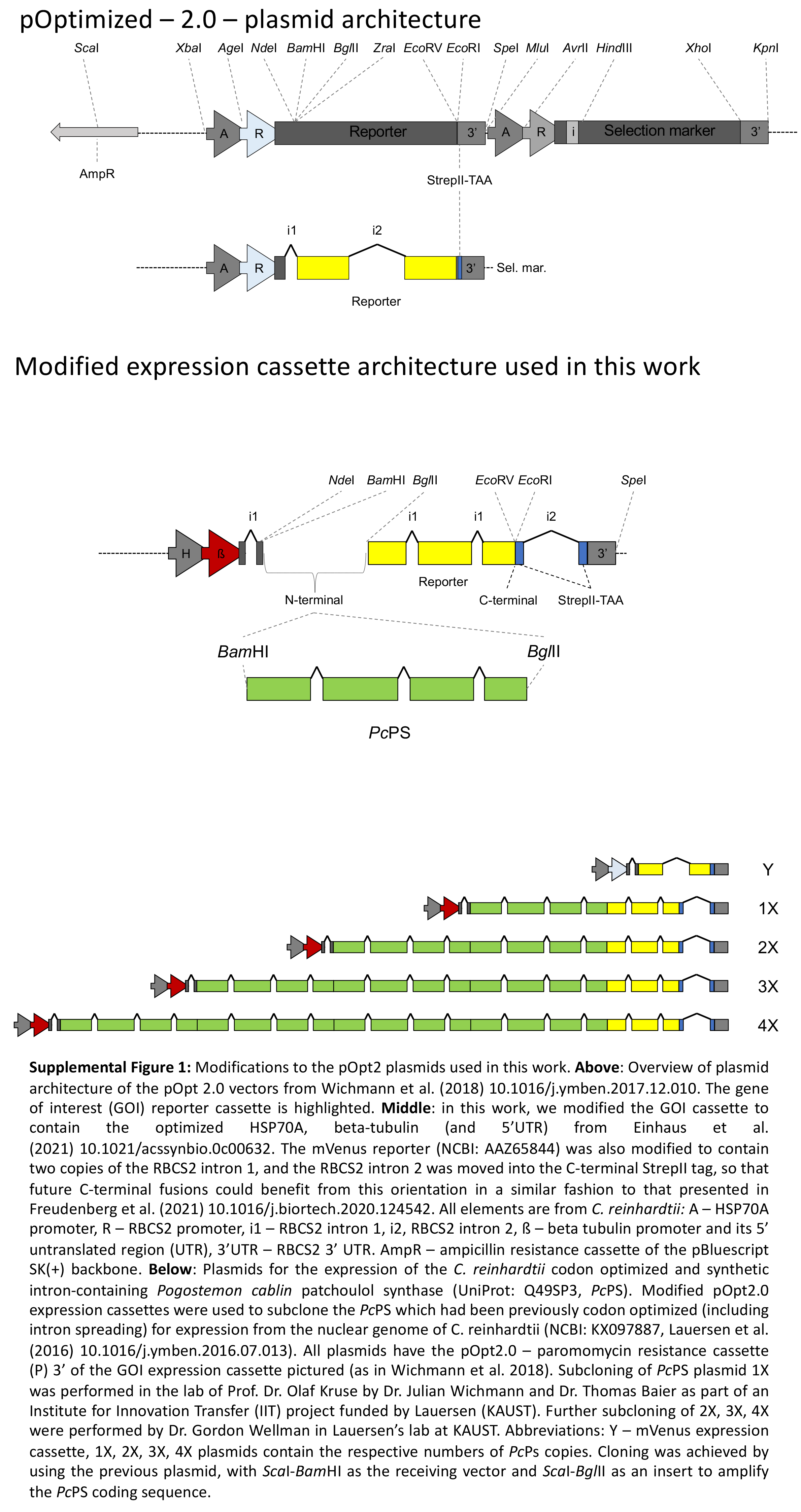

### Supplemental Figure 2

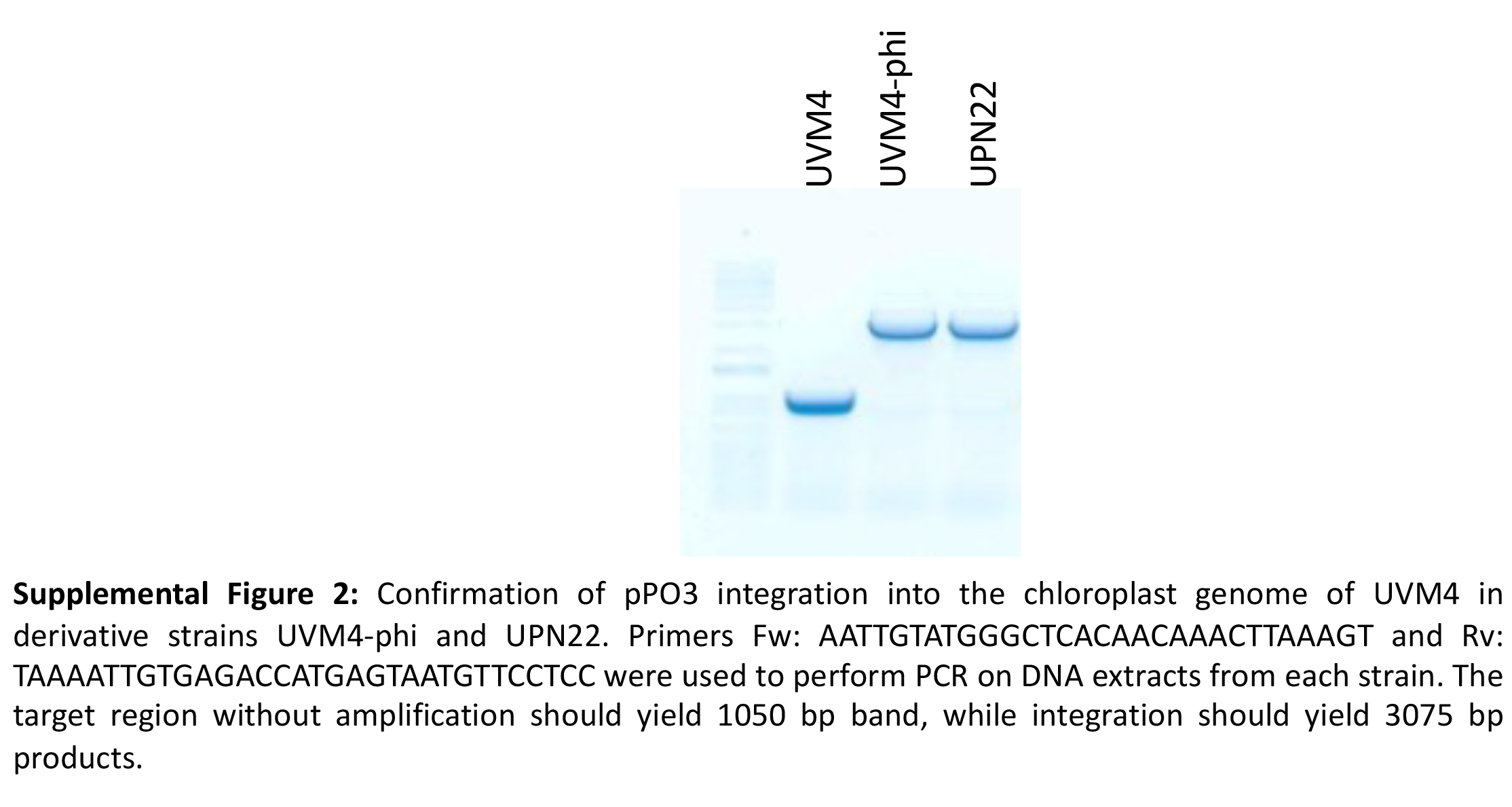

### Supplemental Figure 3

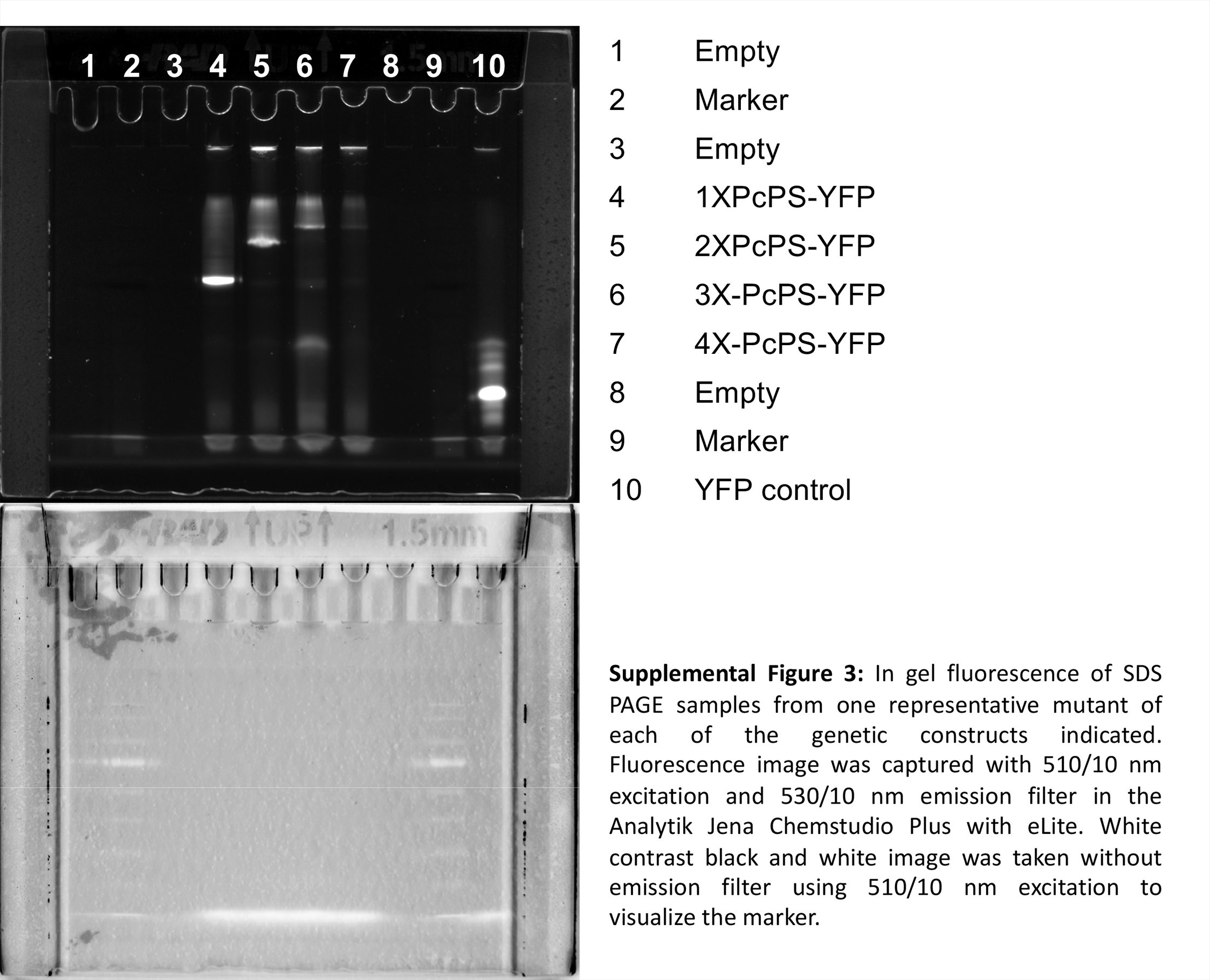

### Supplemental Figure 4

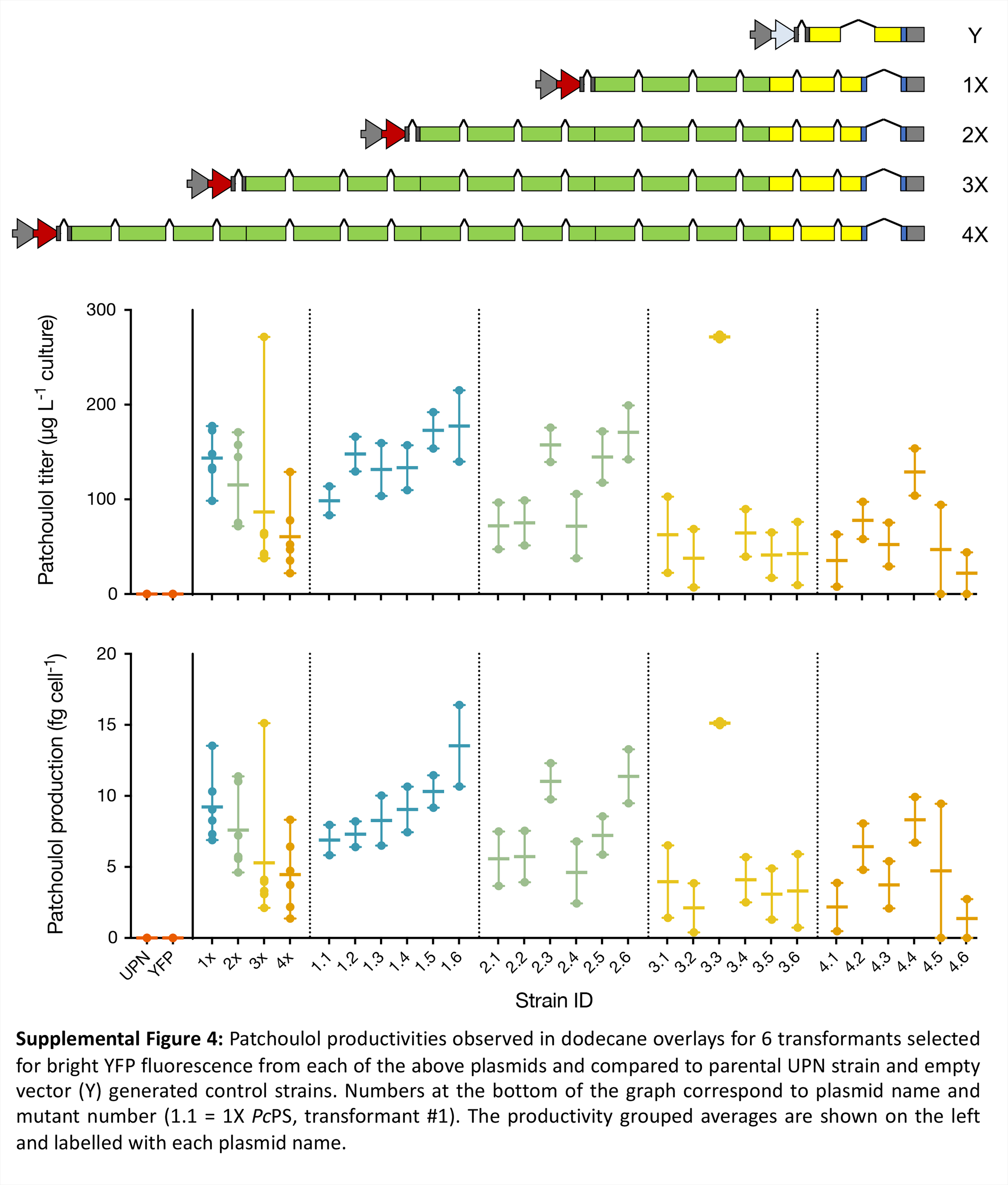

### Supplemental Figure 5

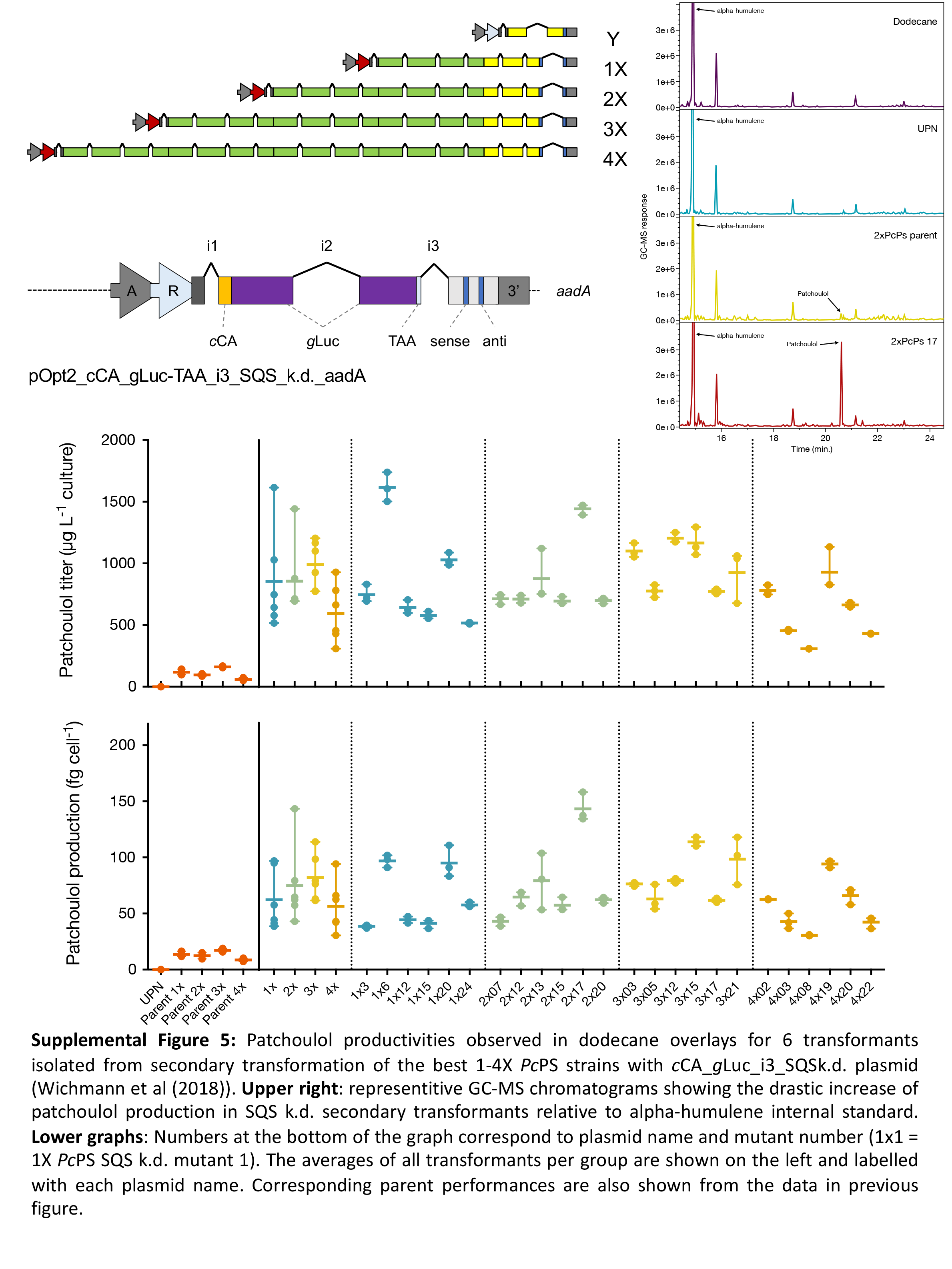
