## Supplemental File 1 - Media Recipes for "Combinatorial engineering for photoautotrophic production of recombinant products from the green microalga *Chlamydomonas reinhardtii*"

**TAP medium**

| | Mw (g/mol) | M | 1 | 2 | 3 | 4 | 5 | L |
|---|---|---|---|---|---|---|---|---|
| TRIS | 121.14 | 0.02 | 2.42 | 4.84 | 7.26 | 9.68 | 12.1 | g |
| TAP-Salts* | - | - | 25 | 50 | 75 | 100 | 125 | mL |
| P-Solution** | - | - | 0.375 | 0.75 | 1.125 | 1.5 | 1.875 | mL |
| Hutner's trace elements | - | - | 1 | 2 | 3 | 4 | 5 | mL |
| Glacial acetic acid | - | - | 1 | 2 | 3 | 4 | 5 | mL |

Adjust final pH to 7.0 with 1M HCL. Generally, 2.2 mL per L
* For Tris-minimal medium, omit the acetic acid and titrate the final solution to pH 7.0 with HCl
** For Phi containing medium, omit P-Solution and add filter sterilised Phi-Solution after autoclaving

| P-Solution (buffering phosph**ate** solution) for TAP | | | | | | | Notes: |
|---|---|---|---|---|---|---|---|
| | Mw (g/mol) | M | 0.1 | L (100 mL) | P M | Final medium P M | |
| $K_2HPO_4$ | 174.176 | 1.653 | 28.80 | g | 2.741 | 1.03 mM | Use 0.375 mL per 1L TAP. Can be added to TAP and autoclaved. |
| $KH_2PO_4$ | 136.086 | 1.088 | 14.80 | g | | | |

| Phi-Solution (buffering phosph**ite**-solution) for TAPhi | | | | | | | Notes: |
|---|---|---|---|---|---|---|---|
| | Mw (g/mol) | M | 0.1 | L (100 mL) | P M | Final medium P M | |
| 50% m/m $K_2HPO_3$:$H_2O$ | 158.18 | 1.653 | 35.80 | mL | 2.741 | 1.03 mM | Use 0.375 mL per 1L TAP. Add filter sterilised after autoclaved. Keep <60°C. See notes on $K_2HPO_3$ supplier. |
| $KH_2PO_3$ | 120.085 | 1.088 | 13.06 | g | | | |

| Phosph**ite** solution - unbuffering for TAPhi | | | | | | | Notes: |
|---|---|---|---|---|---|---|---|
| | Mw (g/mol) | M | 0.1 | L (100 mL) | P M | Final medium P M | |
| $KH_2PO_3$ | 120.085 | 2.741 | 32.92 | g | 2.741 | 1.03 mM | Adapted from Sandoval-Vargas et al. 2018. Must be filter sterilised added after autoclaving to cooled medium. Keep phosphite solution and medium below 60°C |

**6P medium is made with the recipe found below, but replacing 6xP with 6xPhi and filter sterilizing after media cooling**

Following: www.chlamycollection.org/content/uploads/2021/01/optimized-chlamy-medium-by-RAF-II.docx - Freudenberg et al. 2021

| 500x Stock 6xP buffered phosph**ate**-solution (6P-Buffer) | | | | | | | Notes: |
|---|---|---|---|---|---|---|---|
| | Mw (g/mol) | M | 0.1 | L (100 mL) | P M | Final medium P M | |
| $K_2HPO_4$ | 174.176 | 2.2 | 38.32 | g | 3.100 | 6.2 mM | Use 2 mL per 1L 6xP. Can be added to 6xP and autoclaved. |
| $KH_2PO_4$ | 136.086 | 0.9 | 12.25 | g | | | |

| 500x Stock 6xPhi buffered phosph**ite**-solution (6Phi-Buffer) | | | | | | | Notes: |
|---|---|---|---|---|---|---|---|
| | Mw (g/mol) | M | 0.1 | L (100 mL) | P M | Final medium P M | |
| 50% m/m $K_2HPO_3$:$H_2O$ | 158.18 | 2.2 | 47.64 | mL | 3.100 | 6.2 mM | Use 2 mL per 1L 6xPhi. Add filter sterilised after autoclaving to cooled medium. Keep phosphite below 60°C. See notes on $K_2HPO_3$ supplier. |
| $KH_2PO_3$ | 120.085 | 0.9 | 10.81 | g | | | |

**Notes**

| | | |
|---|---|---|
| **$K_2HPO_4$** | Potassium monohygrogen phosph**ate** - Potassium phosph**ate** monobasic - Monopotassium phosph**ate** | |
| | CAS: | 16788-57-1 |
| **$KH_2PO_4$** | Potassium dihygrogen phosph**ate** - Potassium phosph**ate** dibasic - Dipotassium phosph**ate** | |
| | CAS: | 7778-77-0 |
| **$K_2HPO_3$** | Dipotassium hydrogenphosph**ite** | |
| | CAS: | 13492-26-7 |
| | Hygroscopic, so anydrous forms may be difficult to source. We have successfully used a 50% mass percentage solution (Density 1.461 g/mL) from BOC Sciences: https://www.bocsci.com/dipotassium-hydrogenphosphite-cas-13492-26-7-item-48789.html | |
| **$KH_2PO_3$** | Monopotassium phosph**ite** - Potassium dihydrogen phosph**ite** | |
| | CAS: | 13997-65-6 |
| | Anhydrous, sourced from BOC Sciences - https://www.bocsci.com/potassium-dihydrogen-phosphite-cas-13977-65-6-item-9855.html | |
