## Supplemental File 3 - Plasmid Sequence for "Combinatorial engineering for photoautotrophic production of recombinant products from the green microalga *Chlamydomonas reinhardtii*"

```
1XPcPS Plasmid

LOCUS       Exported                9168 bp DNA     circular SYN 07-JAN-2021
DEFINITION  synthetic circular DNA
ACCESSION   .
VERSION     .
KEYWORDS    .
SOURCE      synthetic DNA construct
  ORGANISM  synthetic DNA construct
REFERENCE   1  (bases 1 to 9168)
  AUTHORS   jw
  TITLE     Direct Submission
  JOURNAL   Exported Feb 13, 2022 from SnapGene 5.3.3
            https://www.snapgene.com
FEATURES             Location/Qualifiers
     source          1..9168
                     /organism="synthetic DNA construct"
                     /mol_type="other DNA"
     promoter        6..272
                     /label=HSP70Ap promoter
                     /label=HSP70Ap
     promoter        273..464
                     /label=P-bTUB2
     misc_feature    359..364
                     /label=inserted ATANTT motif
     5'UTR           join(471..546,692..746)
                     /label=5UTR bTUB2
     misc_feature    493..498
                     /label=rebuild ATANTT from ATATT
     misc_feature    536
                     /label=XhoI site deleted T->A
     intron          547..691
                     /label=rbcS2 intron 1
     CDS             750..752
                     /codon_start=1
                     /label=Start
                     /translation="M"
     CDS             join(759..1196,1342..1842,1988..2461,2607..2855)
                     /codon_start=1
                     /label=Patchoulol synthase
                     /translation="ELYAQSVGVGAASRPLANFHPCVWGDKFIVYNPQSCQAGEREEAE
                     ELKVELKRELKEASDNYMRQLKMVDAIQRLGIDYLFVEDVDEALKNLFEMFDAFCKNNH
                     DMHATALSFRLLRQHGYRVSCEVFEKFKDGKDGFKVPNEDGAVAVLEFFEATHLRVHGE
                     DVLDNAFDFTRNYLESVYATLNDPTAKQVHNALNEFSFRRGLPRVEARKYISIYEQYAS
                     HHKGLLKLAKLDFNLVQALHRRELSEDSRWWKTLQVPTKLSFVRDRLVESYFWASGSYF
                     EPNYSVARMILAKGLAVLSLMDDVYDAYGTFEELQMFTDAIERWDASCLDKLPDYMKIV
                     YKALLDVFEEVDEELIKLGAPYRAYYGKEAMKYAARAYMEEAQWREQKHKPTTKEYMKL
                     ATKTCGYITLIILSCLGVEEGIVTKEAFDWVFSRPPFIEATLIIARLVNDITGHEFEKK
                     REHVRTAVECYMEEHKVGKQEVVSEFYNQMESAWKDINEGFLRPVEFPIPLLYLILNSV
                     RTLEVIYKEGDSYTHVGPAMQNIIKQLYLHPVPYGSG"
     intron          1197..1341
                     /label=rbcS2 intron 1
     intron          1843..1987
                     /label=rbcS2 intron 1
     intron          2462..2606
                     /label=rbcS2 intron 1
     CDS             2868..2885
                     /codon_start=1
                     /label=GSGS-Linker
                     /translation="GSGSGS"
     CDS             join(2886..3084,3230..3546,3692..3889)
                     /codon_start=1
                     /label=mVenus
                     /translation="VSKGEELFTGVVPILVELDGDVNGHKFSVSGEGEGDATYGKLTLK
                     LICTTGKLPVPWPTLVTTLGYGLQCFARYPDHMKQHDFFKSAMPEGYVQERTIFFKDDG
                     NYKTRAEVKFEGDTLVNRIELKGIDFKEDGNILGHKLEYNYNSHNVYITADKQKNGIKA
                     NFKIRHNIEDGGVQLADHYQQNTPIGDGPVLLPDNHYLSYQSKLSKDPNEKRDHMVLLE
                     FVTAAGITLGMDELYK"
     intron          3085..3229
                     /label=rbcS2 intron 1
     intron          3547..3691
                     /label=rbcS2 intron 1
     CDS             3890..3907
                     /codon_start=1
                     /label=GSGS-Linker
                     /translation="GSGSGS"
     CDS             join(3920..3938,4268..4269)
                     /codon_start=1
                     /label=GSGS-Linker
                     /translation="GSGSGSG"
     intron          3939..4267
                     /label=rbcS2 intron 2
     CDS             4270..4293
                     /codon_start=1
                     /product="peptide that binds Strep-Tactin(R), an engineered
                     form of streptavidin"
                     /label=Strep-Tag II
                     /translation="WSHPQFEK"
     misc_feature    4297..4530
                     /label=3'UTR
     misc_feature    4531..4542
                     /label=MCS
     promoter        4543..4809
                     /label=HSP70Ap
                     /label=HSP70Ap(1)
     promoter        join(4816..5044,5190..5191)
                     /label=RBCS2p promoter
                     /label=RBCS2p
     intron          5045..5189
                     /label=rbcS2 intron 1
     CDS             5232..6035
                     /codon_start=1
                     /label=APHVIII
                     /translation="MDDALRALRGRYPGCEWVVVEDGASGAGVYRLRGGGRELFVKVAA
                     LGAGVGLLGEAERLVWLAEVGIPVPRVVEGGGDERVAWLVTEAVPGRPASARWPREQRL
                     DVAVALAGLARSLHALDWERCPFDRSLAVTVPQAARAVAEGSVDLEDLDEERKGWSGER
                     LLAELERTRPADEDLAVCHGDLCPDNVLLDPRTCEVTGLIDVGRVGRADRHSDLALVLR
                     ELAHEEDPWFGPECSAAFLREYGRGWDGAVSEEKLAFYRLLDEFF"
     misc_feature    6045..6283
                     /label=3'UTR
                     /label=3'UTR(1)
     rep_origin      complement(6743..7331)
                     /direction=LEFT
                     /label=ori
                     /note="high-copy-number ColE1/pMB1/pBR322/pUC origin of
                     replication"
     CDS             complement(7502..8362)
                     /codon_start=1
                     /gene="bla"
                     /product="beta-lactamase"
                     /label=AmpR
                     /note="confers resistance to ampicillin, carbenicillin, and
                     related antibiotics"
                     /translation="MSIQHFRVALIPFFAAFCLPVFAHPETLVKVKDAEDQLGARVGYI
                     ELDLNSGKILESFRPEERFPMMSTFKVLLCGAVLSRIDAGQEQLGRRIHYSQNDLVEYS
                     PVTEKHLTDGMTVRELCSAAITMSDNTAANLLLTTIGGPKELTAFLHNMGDHVTRLDRW
                     EPELNEAIPNDERDTTMPVAMATTLRKLLTGELLTLASRQQLIDWMEADKVAGPLLRSA
                     LPAGWFIADKSGAGERGSRGIIAALGPDGKPSRIVVIYTTGSQATMDERNRQIAEIGAS
                     LIKHW"
     promoter        complement(8363..8467)
                     /gene="bla"
                     /label=AmpR promoter
     rep_origin      complement(8494..8949)
                     /direction=LEFT
                     /label=f1 ori
                     /note="f1 bacteriophage origin of replication; arrow
                     indicates direction of (+) strand synthesis"
ORIGIN
        1 ctagagctga ggcttgacat gattggtgcg tatgtttgta tgaagctaca ggactgattt
       61 ggcgggctat gagggcgggg gaagctctgg aagggccgcg atggggcgcg cggcgtccag
      121 aaggcgccat acggcccgct ggcggcaccc atccggtata aaagcccgcg accccgaacg
      181 gtgacctcca ctttcagcga caaacgagca cttatacata cgcgactatt ctgccgctat
      241 acataaccac tcagctagct taagatccca tcctggcact ttcttgcgct atgacacttc
      301 cagcaaaagg tagggcgggc tgcgagacgg cttcccggcg ctgcatgcaa caccgatgat
      361 acttatgctt cgaccccccg aagctccttc ggggctgcat gggcgctccg atgccgctcc
      421 agggcgagcg ctgtttaaat agccaggccc ccgactgcaa agacaccggt attatagcga
      481 gctaccaaag ccatacttca aacacctaga tcactaccac ttctacacag gccacacgag
      541 cttgtggtga gtcgacgagc aagcccggcg gatcaggcag cgtgcttgca gatttgactt
      601 gcaacgcccg cattgtgtcg acgaaggctt ttggctcctc tgtcgctgtc tcaagcagca
      661 tctaaccctg cgtcgccgtt tccatttgca gatcgcactc cgctaagggg gcgcctcttc
      721 ctcttcgttt cagtcacaac ccgcaacata tgggatccga gctgtacgcc cagagcgtgg
      781 gcgtgggcgc cgccagccgc cccctggcca acttccaccc ctgcgtgtgg ggcgacaagt
      841 tcatcgtgta caacccccag agctgccagg ccggcgagcg cgaggaggcc gaggagctga
      901 aggtggagct gaagcgcgag ctgaaggagg ccagcgacaa ctacatgcgc cagctgaaga
      961 tggtggacgc catccagcgc ctgggcatcg actacctgtt cgtggaggac gtggacgagg
     1021 ccctgaagaa cctgttcgag atgttcgacg ccttctgcaa gaacaaccac gacatgcacg
     1081 ccaccgccct gagcttccgc ctgctgcgcc agcacggcta ccgcgtgtcc tgcgaggtgt
     1141 tcgagaagtt caaggacggc aaggacggct tcaaggtgcc caacgaggac ggcgcggtga
     1201 gtcgacgagc aagcccggcg gatcaggcag cgtgcttgca gatttgactt gcaacgcccg
     1261 cattgtgtcg acgaaggctt ttggctcctc tgtcgctgtc tcaagcagca tctaaccctg
     1321 cgtcgccgtt tccatttgca ggtggcggtg ctggagttct tcgaggccac ccacctgcgc
     1381 gtgcacggcg aggacgtgct ggacaacgcc ttcgacttca cccgcaacta cctggagagc
     1441 gtgtacgcca ccctgaacga ccccaccgcc aagcaggtgc acaacgcgct gaacgagttc
     1501 tccttccgcc gcggcctgcc ccgcgtggag gcccgcaagt acatcagcat ctacgagcag
     1561 tacgccagcc accacaaggg cctgctgaag ctggccaagc tggacttcaa cctggtgcag
     1621 gcgctgcacc gccgcgagct gtccgaggac agccgctggt ggaagaccct gcaggtgccc
     1681 accaagctga gcttcgtgcg cgaccgcctg gtggagagct acttctgggc cagcggcagc
     1741 tacttcgagc ccaactacag cgtggcccgc atgatcctgg cgaagggcct ggccgtgctg
     1801 agcctgatgg acgacgtgta cgacgcctac ggcaccttcg aggtgagtcg acgagcaagc
     1861 ccggcggatc aggcagcgtg cttgcagatt tgacttgcaa cgcccgcatt gtgtcgacga
     1921 aggcttttgg ctcctctgtc gctgtctcaa gcagcatcta accctgcgtc gccgtttcca
     1981 tttgcaggag ctgcagatgt tcaccgacgc gatcgagcgc tgggacgcca gctgcctgga
     2041 caagctgccc gactacatga agatcgtgta caaggccctg ctggacgtgt tcgaggaggt
     2101 ggacgaggag ctgatcaagc tgggcgcccc ctaccgcgcc tactacggca aggaggccat
     2161 gaagtacgcc gcccgcgcct acatggagga ggcccagtgg cgcgagcaga agcacaagcc
     2221 caccaccaag gagtacatga agctggcgac caagacctgc ggctacatca ccctgatcat
     2281 cctgtcctgc ctgggcgtgg aggagggcat cgtgaccaag gaggcgttcg actgggtgtt
     2341 cagccgcccg ccgttcatcg aggcgaccct gatcatcgcc cgcctggtca acgacatcac
     2401 cggccacgag ttcgagaaga agcgcgagca cgtgcgcacc gccgtggagt gctacatgga
     2461 ggtgagtcga cgagcaagcc cggcggatca ggcagcgtgc ttgcagattt gacttgcaac
     2521 gcccgcattg tgtcgacgaa ggcttttggc tcctctgtcg ctgtctcaag cagcatctaa
     2581 ccctgcgtcg ccgtttccat ttgcaggagc acaaggtggg caagcaggag gtggtgtccg
     2641 agttctacaa ccagatggag agcgcctgga aggacatcaa cgagggcttc ctgcgccccg
     2701 tggagttccc catccccctg ctgtacctga tcctgaacag cgtgcgcacc ctggaggtga
     2761 tctacaagga gggcgacagc tacacccacg tgggcccggc catgcagaac atcatcaagc
     2821 agctgtacct gcaccccgtg ccctacggca gcggcagatc tgacgtcggc agcggcagcg
     2881 gcagcgtgag caagggcgag gagctgttca ccggcgtggt gcccatcctg gtggagctgg
     2941 acggcgacgt gaacggccac aagttcagcg tgagcggcga gggcgagggc gacgccacct
     3001 acggcaagct gaccctgaag ctgatctgca ccaccggcaa gctgcccgtg ccctggccca
     3061 ccctggtgac caccctgggc tacggtgagt cgacgagcaa gcccggcgga tcaggcagcg
     3121 tgcttgcaga tttgacttgc aacgcccgca ttgtgtcgac gaaggctttt ggctcctctg
     3181 tcgctgtctc aagcagcatc taaccctgcg tcgccgtttc catttgcagg cctgcagtgc
     3241 ttcgcccgct accccgacca catgaagcag cacgacttct tcaagagcgc catgcccgag
     3301 ggctacgtgc aggagcgcac catcttcttc aaggacgacg gtaactacaa gacccgcgcc
     3361 gaggtgaagt tcgagggcga caccctggtg aaccgcatcg agctgaaggg catcgacttc
     3421 aaggaggacg gcaacatcct gggccacaag ctggagtaca actacaacag ccacaacgtg
     3481 tacatcaccg ccgacaagca gaagaacggc atcaaggcca acttcaagat ccgccacaac
     3541 atcgaggtga gtcgacgagc aagcccggcg gatcaggcag cgtgcttgca gatttgactt
     3601 gcaacgcccg cattgtgtcg acgaaggctt ttggctcctc tgtcgctgtc tcaagcagca
     3661 tctaaccctg cgtcgccgtt tccatttgca ggacggcggc gtgcagctgg ccgaccacta
     3721 ccagcagaac acccccatcg gcgacggccc cgtgctgctg cccgacaacc actacctgag
     3781 ctaccagagc aagctgagca aggaccccaa cgagaagcgc gaccacatgg tgctgctgga
     3841 gttcgtgacc gccgccggca tcaccctggg catggacgag ctgtacaagg gcagcggcag
     3901 cggcagcgat atcgaattcg gcagcggcag cggctcaggt gagcttgcgg ggttgcgagc
     3961 aacactccag caacgaacag tgcccaagtc aggaatctgc agtcagcctg ggctttcggc
     4021 ggctttttct tgggcaaaca gcttgcactc atgccagcgc ggcttgtcca gcctcacttg
     4081 agctttccag ctgctaccag ccgggctata cgacagcgac agagccatag cgtggaatca
     4141 cttatttggg ttgccgaagt agcggtcgga gcgtgagttc ttggtcaagc cgccccttat
     4201 ccggttcctg tccgtgtctt tgtccctcgt tcacccttcg cggcaccctt catccccttg
     4261 cttgcaggtt ggagccaccc gcagttcgag aagtaaccgc tccgtgtaaa tggaggcgct
     4321 cgttgatctg agccttgccc cctgacgaac ggcggtggat ggaagatact gctctcaagt
     4381 gctgaagcgg tagcttagct ccccgtttcg tgctgatcag tctttttcaa cacgtaaaaa
     4441 gcggaggagt tttgcaattt tgttggttgt aacgatcctc cgttgatttt ggcctctttc
     4501 tccatgggcg ggctgggcgt atttgaagcg actagtacgc gtgctgaggc ttgacatgat
     4561 tggtgcgtat gtttgtatga agctacagga ctgatttggc gggctatgag ggcgggggaa
     4621 gctctggaag ggccgcgatg gggcgcgcgg cgtccagaag gcgccatacg gcccgctggc
     4681 ggcacccatc cggtataaaa gcccgcgacc ccgaacggtg acctccactt tcagcgacaa
     4741 acgagcactt atacatacgc gactattctg ccgctataca taaccactca gctagcttaa
     4801 gatcccatcc ctagggcatg ccgggcgcgc cagaaggagc gcagccaaac caggatgatg
     4861 tttgatgggg tatttgagca cttgcaaccc ttatccggaa gccccctggc ccacaaaggc
     4921 taggcgccaa tgcaagcagt tcgcatgcag cccctggagc ggtgccctcc tgataaaccg
     4981 gccagggggc ctatgttctt tactttttta caagagaagt cactcaacat cttaaaatgg
     5041 ccaggtgagt cgacgagcaa gcccggcgga tcaggcagcg tgcttgcaga tttgacttgc
     5101 aacgcccgca ttgtgtcgac gaaggctttt ggctcctctg tcgctgtctc aagcagcatc
     5161 taaccctgcg tcgccgtttc catttgcagg aagcttactc cgccctcccc ggtgctgaag
     5221 aatttcgaag catggacgat gcgttgcgtg cactgcgggg tcggtatccc ggttgtgagt
     5281 gggttgttgt ggaggatggg gcctcggggg ctggtgttta tcggcttcgg ggtggtgggc
     5341 gggagttgtt tgtcaaggtg gcagctctgg gggccggggt gggcttgttg ggtgaggctg
     5401 agcggctggt gtggttggcg gaggtgggga ttcccgtacc tcgtgttgtg gagggtggtg
     5461 gggacgagag ggtcgcctgg ttggtcaccg aagcggttcc ggggcgtccg gccagtgcgc
     5521 ggtggccgcg ggagcagcgg ctggacgtgg cggtggcgct cgcggggctc gctcgttcgc
     5581 tgcacgcgct ggactgggag cggtgtccgt tcgatcgcag tctcgcggtg acggtgccgc
     5641 aggcggcccg tgctgtcgct gaagggagcg tcgacttgga ggatctggac gaggagcgga
     5701 aggggtggtc gggggagcgg cttctcgccg agctggagcg gactcggcct gcggacgagg
     5761 atctggcggt ttgccacggt gacctgtgcc cggacaacgt gctgctcgac cctcgtacct
     5821 gcgaggtgac cgggctgatc gacgtggggc gggtcggccg tgcggaccgg cactccgatc
     5881 tcgcgctggt gctgcgcgag ctggcccacg aggaggaccc gtggttcggg ccggagtgtt
     5941 ccgcggcgtt cctgcgggag tacgggcgcg ggtgggatgg ggcggtatcg gaggaaaagc
     6001 tggcgtttta ccggctgttg gacgagttct tctgactcga gtgaccgctc cgtgtaaatg
     6061 gaggcgctcg ttgatctgag ccttgccccc tgacgaacgg cggtggatgg aagatactgc
     6121 tctcaagtgc tgaagcggta gcttagctcc ccgtttcgtg ctgatcagtc tttttcaaca
     6181 cgtaaaaagc ggaggagttt tgcaattttg ttggttgtaa cgatcctccg ttgattttgg
     6241 cctctttctc catgggcggg ctgggcgtat ttgaagcgga cccggtaccc agcttttgtt
     6301 ccctttagtg agggttaatt gcgcgcttgg cgtaatcatg gtcatagctg tttcctgtgt
     6361 gaaattgtta tccgctcaca attccacaca acatacgagc cggaagcata aagtgtaaag
     6421 cctggggtgc ctaatgagtg agctaactca cattaattgc gttgcgctca ctgcccgctt
     6481 tccagtcggg aaacctgtcg tgccagctgc attaatgaat cggccaacgc gcggggagag
     6541 gcggtttgcg tattgggcgc tcttccgctt cctcgctcac tgactcgctg cgctcggtcg
     6601 ttcggctgcg gcgagcggta tcagctcact caaaggcggt aatacggtta tccacagaat
     6661 caggggataa cgcaggaaag aacatgtgag caaaaggcca gcaaaaggcc aggaaccgta
     6721 aaaaggccgc gttgctggcg tttttccata ggctccgccc ccctgacgag catcacaaaa
     6781 atcgacgctc aagtcagagg tggcgaaacc cgacaggact ataaagatac caggcgtttc
     6841 cccctggaag ctccctcgtg cgctctcctg ttccgaccct gccgcttacc ggatacctgt
     6901 ccgcctttct cccttcggga agcgtggcgc tttctcatag ctcacgctgt aggtatctca
     6961 gttcggtgta ggtcgttcgc tccaagctgg gctgtgtgca cgaacccccc gttcagcccg
     7021 accgctgcgc cttatccggt aactatcgtc ttgagtccaa cccggtaaga cacgacttat
     7081 cgccactggc agcagccact ggtaacagga ttagcagagc gaggtatgta ggcggtgcta
     7141 cagagttctt gaagtggtgg cctaactacg gctacactag aaggacagta tttggtatct
     7201 gcgctctgct gaagccagtt accttcggaa aaagagttgg tagctcttga tccggcaaac
     7261 aaaccaccgc tggtagcggt ggtttttttg tttgcaagca gcagattacg cgcagaaaaa
     7321 aaggatctca agaagatcct ttgatctttt ctacggggtc tgacgctcag tggaacgaaa
     7381 actcacgtta agggattttg gtcatgagat tatcaaaaag gatcttcacc tagatccttt
     7441 taaattaaaa atgaagtttt aaatcaatct aaagtatata tgagtaaact tggtctgaca
     7501 gttaccaatg cttaatcagt gaggcaccta tctcagcgat ctgtctattt cgttcatcca
     7561 tagttgcctg actccccgtc gtgtagataa ctacgatacg ggagggctta ccatctggcc
     7621 ccagtgctgc aatgataccg cgagacccac gctcaccggc tccagattta tcagcaataa
     7681 accagccagc cggaagggcc gagcgcagaa gtggtcctgc aactttatcc gcctccatcc
     7741 agtctattaa ttgttgccgg gaagctagag taagtagttc gccagttaat agtttgcgca
     7801 acgttgttgc cattgctaca ggcatcgtgg tgtcacgctc gtcgtttggt atggcttcat
     7861 tcagctccgg ttcccaacga tcaaggcgag ttacatgatc ccccatgttg tgcaaaaaag
     7921 cggttagctc cttcggtcct ccgatcgttg tcagaagtaa gttggccgca gtgttatcac
     7981 tcatggttat ggcagcactg cataattctc ttactgtcat gccatccgta agatgctttt
     8041 ctgtgactgg tgagtactca accaagtcat tctgagaata gtgtatgcgg cgaccgagtt
     8101 gctcttgccc ggcgtcaata cgggataata ccgcgccaca tagcagaact ttaaaagtgc
     8161 tcatcattgg aaaacgttct tcggggcgaa aactctcaag gatcttaccg ctgttgagat
     8221 ccagttcgat gtaacccact cgtgcaccca actgatcttc agcatctttt actttcacca
     8281 gcgtttctgg gtgagcaaaa acaggaaggc aaaatgccgc aaaaaaggga ataagggcga
     8341 cacggaaatg ttgaatactc atactcttcc tttttcaata ttattgaagc atttatcagg
     8401 gttattgtct catgagcgga tacatatttg aatgtattta gaaaaataaa caaatagggg
     8461 ttccgcgcac atttccccga aaagtgccac actaaattgt aagcgttaat attttgttaa
     8521 aattcgcgtt aaatttttgt taaatcagct cattttttaa ccaataggcc gaaatcggca
     8581 aaatccctta taaatcaaaa gaatagaccg agatagggtt gagtgttgtt ccagtttgga
     8641 acaagagtcc actattaaag aacgtggact ccaacgtcaa agggcgaaaa accgtctatc
     8701 agggcgatgg cccactacgt gaaccatcac cctaatcaag ttttttgggg tcgaggtgcc
     8761 gtaaagcact aaatcggaac cctaaaggga gcccccgatt tagagcttga cggggaaagc
     8821 cggcgaacgt ggcgagaaag gaagggaaga aagcgaaagg agcgggcgct agggcgctgg
     8881 caagtgtagc ggtcacgctg cgcgtaacca ccacacccgc cgcgcttaat gcgccgctac
     8941 agggcgcgtc ccattcgcca ttcaggctgc gcaactgttg ggaagggcga tcggtgcggg
     9001 cctcttcgct attacgccag ctggcgaaag ggggatgtgc tgcaaggcga ttaagttggg
     9061 taacgccagg gttttcccag tcacgacgtt gtaaaacgac ggccagtgag cgcgcgtaat
     9121 acgactcact atagggcgaa ttggagctcc accgcggtgg cggccgct
//
```

**2XPcPS Plasmid**

```
LOCUS       Exported               11271 bp DNA     circular SYN 04-SEP-2021
DEFINITION  synthetic circular DNA
ACCESSION   .
VERSION     .
KEYWORDS    .
SOURCE      synthetic DNA construct
  ORGANISM  recombinant plasmid
REFERENCE   1  (bases 1 to 11271)
  AUTHORS   Gordon Wellman
  TITLE     Direct Submission
  JOURNAL   Exported Feb 13, 2022 from SnapGene 5.3.3
            https://www.snapgene.com
FEATURES             Location/Qualifiers
     source          1..11271
                     /organism="recombinant plasmid"
                     /mol_type="other DNA"
     promoter        8..274
                     /label=HSP70Ap promoter
                     /label=HSP70Ap
     promoter        275..466
                     /label=P-bTUB2
     misc_feature    361..366
                     /label=inserted ATANTT motif
     5'UTR           join(473..548,694..748)
                     /label=5UTR bTUB2
     misc_feature    495..500
                     /label=rebuild ATANTT from ATATT
     misc_feature    538
                     /label=XhoI site deleted T->A
     intron          549..693
                     /label=rbcS2 intron 1
     CDS             752..754
                     /codon_start=1
                     /label=Start
                     /translation="M"
     CDS             join(761..1198,1344..1844,1990..2463,2609..2857)
                     /codon_start=1
                     /label=Patchoulol synthase
                     /translation="ELYAQSVGVGAASRPLANFHPCVWGDKFIVYNPQSCQAGEREEAE
                     ELKVELKRELKEASDNYMRQLKMVDAIQRLGIDYLFVEDVDEALKNLFEMFDAFCKNNH
                     DMHATALSFRLLRQHGYRVSCEVFEKFKDGKDGFKVPNEDGAVAVLEFFEATHLRVHGE
                     DVLDNAFDFTRNYLESVYATLNDPTAKQVHNALNEFSFRRGLPRVEARKYISIYEQYAS
                     HHKGLLKLAKLDFNLVQALHRRELSEDSRWWKTLQVPTKLSFVRDRLVESYFWASGSYF
                     EPNYSVARMILAKGLAVLSLMDDVYDAYGTFEELQMFTDAIERWDASCLDKLPDYMKIV
                     YKALLDVFEEVDEELIKLGAPYRAYYGKEAMKYAARAYMEEAQWREQKHKPTTKEYMKL
                     ATKTCGYITLIILSCLGVEEGIVTKEAFDWVFSRPPFIEATLIIARLVNDITGHEFEKK
                     REHVRTAVECYMEEHKVGKQEVVSEFYNQMESAWKDINEGFLRPVEFPIPLLYLILNSV
                     RTLEVIYKEGDSYTHVGPAMQNIIKQLYLHPVPYGSG"
     intron          1199..1343
                     /label=rbcS2 intron 1
     intron          1845..1989
                     /label=rbcS2 intron 1
     intron          2464..2608
                     /label=rbcS2 intron 1
     CDS             join(2864..3301,3447..3947,4093..4566,4712..4960)
                     /codon_start=1
                     /label=Patchoulol synthase
                     /translation="ELYAQSVGVGAASRPLANFHPCVWGDKFIVYNPQSCQAGEREEAE
                     ELKVELKRELKEASDNYMRQLKMVDAIQRLGIDYLFVEDVDEALKNLFEMFDAFCKNNH
                     DMHATALSFRLLRQHGYRVSCEVFEKFKDGKDGFKVPNEDGAVAVLEFFEATHLRVHGE
                     DVLDNAFDFTRNYLESVYATLNDPTAKQVHNALNEFSFRRGLPRVEARKYISIYEQYAS
                     HHKGLLKLAKLDFNLVQALHRRELSEDSRWWKTLQVPTKLSFVRDRLVESYFWASGSYF
                     EPNYSVARMILAKGLAVLSLMDDVYDAYGTFEELQMFTDAIERWDASCLDKLPDYMKIV
                     YKALLDVFEEVDEELIKLGAPYRAYYGKEAMKYAARAYMEEAQWREQKHKPTTKEYMKL
                     ATKTCGYITLIILSCLGVEEGIVTKEAFDWVFSRPPFIEATLIIARLVNDITGHEFEKK
                     REHVRTAVECYMEEHKVGKQEVVSEFYNQMESAWKDINEGFLRPVEFPIPLLYLILNSV
                     RTLEVIYKEGDSYTHVGPAMQNIIKQLYLHPVPYGSG"
     intron          3302..3446
                     /label=rbcS2 intron 1
     intron          3948..4092
                     /label=rbcS2 intron 1
     intron          4567..4711
                     /label=rbcS2 intron 1
     CDS             4973..4990
                     /codon_start=1
                     /label=GSGS-Linker
                     /translation="GSGSGS"
     CDS             join(4991..5189,5335..5651,5797..5994)
                     /codon_start=1
                     /label=mVenus
                     /translation="VSKGEELFTGVVPILVELDGDVNGHKFSVSGEGEGDATYGKLTLK
                     LICTTGKLPVPWPTLVTTLGYGLQCFARYPDHMKQHDFFKSAMPEGYVQERTIFFKDDG
                     NYKTRAEVKFEGDTLVNRIELKGIDFKEDGNILGHKLEYNYNSHNVYITADKQKNGIKA
                     NFKIRHNIEDGGVQLADHYQQNTPIGDGPVLLPDNHYLSYQSKLSKDPNEKRDHMVLLE
                     FVTAAGITLGMDELYK"
     intron          5190..5334
                     /label=rbcS2 intron 1
     intron          5652..5796
                     /label=rbcS2 intron 1
     CDS             5995..6012
                     /codon_start=1
                     /label=GSGS-Linker
                     /translation="GSGSGS"
     CDS             join(6025..6043,6373..6374)
                     /codon_start=1
                     /label=GSGS-Linker
                     /translation="GSGSGSG"
     intron          6044..6372
                     /label=rbcS2 intron 2
     CDS             6375..6398
                     /codon_start=1
                     /product="peptide that binds Strep-Tactin(R), an engineered
                     form of streptavidin"
                     /label=Strep-Tag II
                     /translation="WSHPQFEK"
     misc_feature    6402..6635
                     /label=3'UTR
     misc_feature    6636..6647
                     /label=MCS
     promoter        6648..6914
                     /label=HSP70Ap
                     /label=HSP70Ap(1)
     promoter        join(6921..7149,7295..7296)
                     /label=RBCS2p promoter
                     /label=RBCS2p
     intron          7150..7294
                     /label=rbcS2 intron 1
     CDS             7337..8140
                     /codon_start=1
                     /label=APHVIII
                     /translation="MDDALRALRGRYPGCEWVVVEDGASGAGVYRLRGGGRELFVKVAA
                     LGAGVGLLGEAERLVWLAEVGIPVPRVVEGGGDERVAWLVTEAVPGRPASARWPREQRL
                     DVAVALAGLARSLHALDWERCPFDRSLAVTVPQAARAVAEGSVDLEDLDEERKGWSGER
                     LLAELERTRPADEDLAVCHGDLCPDNVLLDPRTCEVTGLIDVGRVGRADRHSDLALVLR
                     ELAHEEDPWFGPECSAAFLREYGRGWDGAVSEEKLAFYRLLDEFF"
     misc_feature    8150..8388
                     /label=3'UTR
                     /label=3'UTR(1)
     rep_origin      complement(8848..9436)
                     /direction=LEFT
                     /label=ori
                     /note="high-copy-number ColE1/pMB1/pBR322/pUC origin of
                     replication"
     CDS             complement(9607..10467)
                     /codon_start=1
                     /gene="bla"
                     /product="beta-lactamase"
                     /label=AmpR
                     /note="confers resistance to ampicillin, carbenicillin, and
                     related antibiotics"
                     /translation="MSIQHFRVALIPFFAAFCLPVFAHPETLVKVKDAEDQLGARVGYI
                     ELDLNSGKILESFRPEERFPMMSTFKVLLCGAVLSRIDAGQEQLGRRIHYSQNDLVEYS
                     PVTEKHLTDGMTVRELCSAAITMSDNTAANLLLTTIGGPKELTAFLHNMGDHVTRLDRW
                     EPELNEAIPNDERDTTMPVAMATTLRKLLTGELLTLASRQQLIDWMEADKVAGPLLRSA
                     LPAGWFIADKSGAGERGSRGIIAALGPDGKPSRIVVIYTTGSQATMDERNRQIAEIGAS
                     LIKHW"
     promoter        complement(10468..10572)
                     /gene="bla"
                     /label=AmpR promoter
     rep_origin      complement(10599..11054)
                     /direction=LEFT
                     /label=f1 ori
                     /note="f1 bacteriophage origin of replication; arrow
                     indicates direction of (+) strand synthesis"
ORIGIN
        1 ctctagagct gaggcttgac atgattggtg cgtatgtttg tatgaagcta caggactgat
       61 ttggcgggct atgagggcgg gggaagctct ggaagggccg cgatggggcg cgcggcgtcc
      121 agaaggcgcc atacggcccg ctggcggcac ccatccggta taaaagcccg cgaccccgaa
      181 cggtgacctc cactttcagc gacaaacgag cacttataca tacgcgacta ttctgccgct
      241 atacataacc actcagctag cttaagatcc catcctggca ctttcttgcg ctatgacact
      301 tccagcaaaa ggtagggcgg gctgcgagac ggcttcccgg cgctgcatgc aacaccgatg
      361 atacttatgc ttcgaccccc cgaagctcct tcggggctgc atgggcgctc cgatgccgct
      421 ccagggcgag cgctgtttaa atagccaggc ccccgactgc aaagacaccg gtattatagc
      481 gagctaccaa agccatactt caaacaccta gatcactacc acttctacac aggccacacg
      541 agcttgtggt gagtcgacga gcaagcccgg cggatcaggc agcgtgcttg cagatttgac
      601 ttgcaacgcc cgcattgtgt cgacgaaggc ttttggctcc tctgtcgctg tctcaagcag
      661 catctaaccc tgcgtcgccg tttccatttg cagatcgcac tccgctaagg gggcgcctct
      721 tcctcttcgt ttcagtcaca acccgcaaca tatgggatcc gagctgtacg cccagagcgt
      781 gggcgtgggc gccgccagcc gccccctggc caacttccac ccctgcgtgt ggggcgacaa
      841 gttcatcgtg tacaaccccc agagctgcca ggccggcgag cgcgaggagg ccgaggagct
      901 gaaggtggag ctgaagcgcg agctgaagga ggccagcgac aactacatgc gccagctgaa
      961 gatggtggac gccatccagc gcctgggcat cgactacctg ttcgtggagg acgtggacga
     1021 ggccctgaag aacctgttcg agatgttcga cgccttctgc aagaacaacc acgacatgca
     1081 cgccaccgcc ctgagcttcc gcctgctgcg ccagcacggc taccgcgtgt cctgcgaggt
     1141 gttcgagaag ttcaaggacg gcaaggacgg cttcaaggtg cccaacgagg acggcgcggt
     1201 gagtcgacga gcaagcccgg cggatcaggc agcgtgcttg cagatttgac ttgcaacgcc
     1261 cgcattgtgt cgacgaaggc ttttggctcc tctgtcgctg tctcaagcag catctaaccc
     1321 tgcgtcgccg tttccatttg caggtggcgg tgctggagtt cttcgaggcc acccacctgc
     1381 gcgtgcacgg cgaggacgtg ctggacaacg ccttcgactt cacccgcaac tacctggaga
     1441 gcgtgtacgc caccctgaac gaccccaccg ccaagcaggt gcacaacgcg ctgaacgagt
     1501 tctccttccg ccgcggcctg ccccgcgtgg aggcccgcaa gtacatcagc atctacgagc
     1561 agtacgccag ccaccacaag ggcctgctga agctggccaa gctggacttc aacctggtgc
     1621 aggcgctgca ccgccgcgag ctgtccgagg acagccgctg gtggaagacc ctgcaggtgc
     1681 ccaccaagct gagcttcgtg cgcgaccgcc tggtggagag ctacttctgg gccagcggca
     1741 gctacttcga gcccaactac agcgtggccc gcatgatcct ggcgaagggc ctggccgtgc
     1801 tgagcctgat ggacgacgtg tacgacgcct acggcacctt cgaggtgagt cgacgagcaa
     1861 gcccggcgga tcaggcagcg tgcttgcaga tttgacttgc aacgcccgca ttgtgtcgac
     1921 gaaggctttt ggctcctctg tcgctgtctc aagcagcatc taaccctgcg tcgccgtttc
     1981 catttgcagg agctgcagat gttcaccgac gcgatcgagc gctgggacgc cagctgcctg
     2041 gacaagctgc ccgactacat gaagatcgtg tacaaggccc tgctggacgt gttcgaggag
     2101 gtggacgagg agctgatcaa gctgggcgcc ccctaccgcg cctactacgg caaggaggcc
     2161 atgaagtacg ccgcccgcgc ctacatggag gaggcccagt ggcgcgagca gaagcacaag
     2221 cccaccacca aggagtacat gaagctggcg accaagacct gcggctacat caccctgatc
     2281 atcctgtcct gcctgggcgt ggaggagggc atcgtgacca aggaggcgtt cgactgggtg
     2341 ttcagccgcc cgccgttcat cgaggcgacc ctgatcatcg cccgcctggt caacgacatc
     2401 accggccacg agttcgagaa gaagcgcgag cacgtgcgca ccgccgtgga gtgctacatg
     2461 gaggtgagtc gacgagcaag cccggcggat caggcagcgt gcttgcagat ttgacttgca
     2521 acgcccgcat tgtgtcgacg aaggcttttg gctcctctgt cgctgtctca agcagcatct
     2581 aaccctgcgt cgccgtttcc atttgcagga gcacaaggtg ggcaagcagg aggtggtgtc
     2641 cgagttctac aaccagatgg agagcgcctg gaaggacatc aacgagggct tcctgcgccc
     2701 cgtggagttc cccatccccc tgctgtacct gatcctgaac agcgtgcgca ccctggaggt
     2761 gatctacaag gagggcgaca gctacaccca cgtgggcccg gccatgcaga acatcatcaa
     2821 gcagctgtac ctgcaccccg tgccctacgg cagcggcaga tccgagctgt acgcccagag
     2881 cgtgggcgtg ggcgccgcca gccgccccct ggccaacttc cacccctgcg tgtggggcga
     2941 caagttcatc gtgtacaacc cccagagctg ccaggccggc gagcgcgagg aggccgagga
     3001 gctgaaggtg gagctgaagc gcgagctgaa ggaggccagc gacaactaca tgcgccagct
     3061 gaagatggtg gacgccatcc agcgcctggg catcgactac ctgttcgtgg aggacgtgga
     3121 cgaggccctg aagaacctgt tcgagatgtt cgacgccttc tgcaagaaca accacgacat
     3181 gcacgccacc gccctgagct tccgcctgct gcgccagcac ggctaccgcg tgtcctgcga
     3241 ggtgttcgag aagttcaagg acggcaagga cggcttcaag gtgcccaacg aggacggcgc
     3301 ggtgagtcga cgagcaagcc cggcggatca ggcagcgtgc ttgcagattt gacttgcaac
     3361 gcccgcattg tgtcgacgaa ggcttttggc tcctctgtcg ctgtctcaag cagcatctaa
     3421 ccctgcgtcg ccgtttccat ttgcaggtgg cggtgctgga gttcttcgag gccacccacc
     3481 tgcgcgtgca cggcgaggac gtgctggaca acgccttcga cttcacccgc aactacctgg
     3541 agagcgtgta cgccaccctg aacgacccca ccgccaagca ggtgcacaac gcgctgaacg
     3601 agttctcctt ccgccgcggc ctgccccgcg tggaggcccg caagtacatc agcatctacg
     3661 agcagtacgc cagccaccac aagggcctgc tgaagctggc caagctggac ttcaacctgg
     3721 tgcaggcgct gcaccgccgc gagctgtccg aggacagccg ctggtggaag accctgcagg
     3781 tgcccaccaa gctgagcttc gtgcgcgacc gcctggtgga gagctacttc tgggccagcg
     3841 gcagctactt cgagcccaac tacagcgtgg cccgcatgat cctggcgaag ggcctggccg
     3901 tgctgagcct gatggacgac gtgtacgacg cctacggcac cttcgaggtg agtcgacgag
     3961 caagcccggc ggatcaggca gcgtgcttgc agatttgact tgcaacgccc gcattgtgtc
     4021 gacgaaggct tttggctcct ctgtcgctgt ctcaagcagc atctaaccct gcgtcgccgt
     4081 ttccatttgc aggagctgca gatgttcacc gacgcgatcg agcgctggga cgccagctgc
     4141 ctggacaagc tgcccgacta catgaagatc gtgtacaagg ccctgctgga cgtgttcgag
     4201 gaggtggacg aggagctgat caagctgggc gccccctacc gcgcctacta cggcaaggag
     4261 gccatgaagt acgccgcccg cgcctacatg gaggaggccc agtggcgcga gcagaagcac
     4321 aagcccacca ccaaggagta catgaagctg gcgaccaaga cctgcggcta catcaccctg
     4381 atcatcctgt cctgcctggg cgtggaggag ggcatcgtga ccaaggaggc gttcgactgg
     4441 gtgttcagcc gcccgccgtt catcgaggcg accctgatca tcgcccgcct ggtcaacgac
     4501 atcaccggcc acgagttcga gaagaagcgc gagcacgtgc gcaccgccgt ggagtgctac
     4561 atggaggtga gtcgacgagc aagcccggcg gatcaggcag cgtgcttgca gatttgactt
     4621 gcaacgcccg cattgtgtcg acgaaggctt ttggctcctc tgtcgctgtc tcaagcagca
     4681 tctaaccctg cgtcgccgtt tccatttgca ggagcacaag gtgggcaagc aggaggtggt
     4741 gtccgagttc tacaaccaga tggagagcgc ctggaaggac atcaacgagg gcttcctgcg
     4801 ccccgtggag ttccccatcc ccctgctgta cctgatcctg aacagcgtgc gcaccctgga
     4861 ggtgatctac aaggagggcg acagctacac ccacgtgggc ccggccatgc agaacatcat
     4921 caagcagctg tacctgcacc ccgtgcccta cggcagcggc agatctgacg tcggcagcgg
     4981 cagcggcagc gtgagcaagg gcgaggagct gttcaccggc gtggtgccca tcctggtgga
     5041 gctggacggc gacgtgaacg gccacaagtt cagcgtgagc ggcgagggcg agggcgacgc
     5101 cacctacggc aagctgaccc tgaagctgat ctgcaccacc ggcaagctgc ccgtgccctg
     5161 gcccaccctg gtgaccaccc tgggctacgg tgagtcgacg agcaagcccg gcggatcagg
     5221 cagcgtgctt gcagatttga cttgcaacgc ccgcattgtg tcgacgaagg cttttggctc
     5281 ctctgtcgct gtctcaagca gcatctaacc ctgcgtcgcc gtttccattt gcaggcctgc
     5341 agtgcttcgc ccgctacccc gaccacatga agcagcacga cttcttcaag agcgccatgc
     5401 ccgagggcta cgtgcaggag cgcaccatct tcttcaagga cgacggtaac tacaagaccc
     5461 gcgccgaggt gaagttcgag ggcgacaccc tggtgaaccg catcgagctg aagggcatcg
     5521 acttcaagga ggacggcaac atcctgggcc acaagctgga gtacaactac aacagccaca
     5581 acgtgtacat caccgccgac aagcagaaga acggcatcaa ggccaacttc aagatccgcc
     5641 acaacatcga ggtgagtcga cgagcaagcc cggcggatca ggcagcgtgc ttgcagattt
     5701 gacttgcaac gcccgcattg tgtcgacgaa ggcttttggc tcctctgtcg ctgtctcaag
     5761 cagcatctaa ccctgcgtcg ccgtttccat ttgcaggacg gcggcgtgca gctggccgac
     5821 cactaccagc agaacacccc catcggcgac ggccccgtgc tgctgcccga caaccactac
     5881 ctgagctacc agagcaagct gagcaaggac cccaacgaga agcgcgacca catggtgctg
     5941 ctggagttcg tgaccgccgc cggcatcacc ctgggcatgg acgagctgta caagggcagc
     6001 ggcagcggca gcgatatcga attcggcagc ggcagcggct caggtgagct tgcggggttg
     6061 cgagcaacac tccagcaacg aacagtgccc aagtcaggaa tctgcagtca gcctgggctt
     6121 tcggcggctt tttcttgggc aaacagcttg cactcatgcc agcgcggctt gtccagcctc
     6181 acttgagctt tccagctgct accagccggg ctatacgaca gcgacagagc catagcgtgg
     6241 aatcacttat ttgggttgcc gaagtagcgg tcggagcgtg agttcttggt caagccgccc
     6301 cttatccggt tcctgtccgt gtctttgtcc ctcgttcacc cttcgcggca cccttcatcc
     6361 ccttgcttgc aggttggagc cacccgcagt tcgagaagta accgctccgt gtaaatggag
     6421 gcgctcgttg atctgagcct tgccccctga cgaacggcgg tggatggaag atactgctct
     6481 caagtgctga agcggtagct tagctccccg tttcgtgctg atcagtcttt ttcaacacgt
     6541 aaaaagcgga ggagttttgc aattttgttg gttgtaacga tcctccgttg attttggcct
     6601 ctttctccat gggcgggctg ggcgtatttg aagcgactag tacgcgtgct gaggcttgac
     6661 atgattggtg cgtatgtttg tatgaagcta caggactgat ttggcgggct atgagggcgg
     6721 gggaagctct ggaagggccg cgatggggcg cgcggcgtcc agaaggcgcc atacggcccg
     6781 ctggcggcac ccatccggta taaaagcccg cgaccccgaa cggtgacctc cactttcagc
     6841 gacaaacgag cacttataca tacgcgacta ttctgccgct atacataacc actcagctag
     6901 cttaagatcc catccctagg gcatgccggg cgcgccagaa ggagcgcagc caaaccagga
     6961 tgatgtttga tggggtattt gagcacttgc aacccttatc cggaagcccc ctggcccaca
     7021 aaggctaggc gccaatgcaa gcagttcgca tgcagcccct ggagcggtgc cctcctgata
     7081 aaccggccag ggggcctatg ttctttactt ttttacaaga gaagtcactc aacatcttaa
     7141 aatggccagg tgagtcgacg agcaagcccg gcggatcagg cagcgtgctt gcagatttga
     7201 cttgcaacgc ccgcattgtg tcgacgaagg cttttggctc ctctgtcgct gtctcaagca
     7261 gcatctaacc ctgcgtcgcc gtttccattt gcaggaagct tactccgccc tccccggtgc
     7321 tgaagaattt cgaagcatgg acgatgcgtt gcgtgcactg cggggtcggt atcccggttg
     7381 tgagtgggtt gttgtggagg atggggcctc gggggctggt gtttatcggc ttcggggtgg
     7441 tgggcgggag ttgtttgtca aggtggcagc tctgggggcc ggggtgggct tgttgggtga
     7501 ggctgagcgg ctggtgtggt tggcggaggt ggggattccc gtacctcgtg ttgtggaggg
     7561 tggtggggac gagagggtcg cctggttggt caccgaagcg gttccggggc gtccggccag
     7621 tgcgcggtgg ccgcgggagc agcggctgga cgtggcggtg gcgctcgcgg ggctcgctcg
     7681 ttcgctgcac gcgctggact gggagcggtg tccgttcgat cgcagtctcg cggtgacggt
     7741 gccgcaggcg gcccgtgctg tcgctgaagg gagcgtcgac ttggaggatc tggacgagga
     7801 gcggaagggg tggtcggggg agcggcttct cgccgagctg gagcggactc ggcctgcgga
     7861 cgaggatctg gcggtttgcc acggtgacct gtgcccggac aacgtgctgc tcgaccctcg
     7921 tacctgcgag gtgaccgggc tgatcgacgt ggggcgggtc ggccgtgcgg accggcactc
     7981 cgatctcgcg ctggtgctgc gcgagctggc ccacgaggag gacccgtggt tcgggccgga
     8041 gtgttccgcg gcgttcctgc gggagtacgg gcgcgggtgg gatggggcgg tatcggagga
     8101 aaagctggcg ttttaccggc tgttggacga gttcttctga ctcgagtgac cgctccgtgt
     8161 aaatggaggc gctcgttgat ctgagccttg ccccctgacg aacggcggtg gatggaagat
     8221 actgctctca agtgctgaag cggtagctta gctccccgtt tcgtgctgat cagtcttttt
     8281 caacacgtaa aaagcggagg agttttgcaa ttttgttggt tgtaacgatc ctccgttgat
     8341 tttggcctct ttctccatgg gcgggctggg cgtatttgaa gcggacccgg tacccagctt
     8401 ttgttccctt tagtgagggt taattgcgcg cttggcgtaa tcatggtcat agctgtttcc
     8461 tgtgtgaaat tgttatccgc tcacaattcc acacaacata cgagccggaa gcataaagtg
     8521 taaagcctgg ggtgcctaat gagtgagcta actcacatta attgcgttgc gctcactgcc
     8581 cgctttccag tcgggaaacc tgtcgtgcca gctgcattaa tgaatcggcc aacgcgcggg
     8641 gagaggcggt ttgcgtattg ggcgctcttc cgcttcctcg ctcactgact cgctgcgctc
     8701 ggtcgttcgg ctgcggcgag cggtatcagc tcactcaaag gcggtaatac ggttatccac
     8761 agaatcaggg gataacgcag gaaagaacat gtgagcaaaa ggccagcaaa aggccaggaa
     8821 ccgtaaaaag gccgcgttgc tggcgttttt ccataggctc cgcccccctg acgagcatca
     8881 caaaaatcga cgctcaagtc agaggtggcg aaacccgaca ggactataaa gataccaggc
     8941 gtttccccct ggaagctccc tcgtgcgctc tcctgttccg accctgccgc ttaccggata
     9001 cctgtccgcc tttctccctt cgggaagcgt ggcgctttct catagctcac gctgtaggta
     9061 tctcagttcg gtgtaggtcg ttcgctccaa gctgggctgt gtgcacgaac cccccgttca
     9121 gcccgaccgc tgcgccttat ccggtaacta tcgtcttgag tccaacccgg taagacacga
     9181 cttatcgcca ctggcagcag ccactggtaa caggattagc agagcgaggt atgtaggcgg
     9241 tgctacagag ttcttgaagt ggtggcctaa ctacggctac actagaagga cagtatttgg
     9301 tatctgcgct ctgctgaagc cagttacctt cggaaaaaga gttggtagct cttgatccgg
     9361 caaacaaacc accgctggta gcggtggttt ttttgtttgc aagcagcaga ttacgcgcag
     9421 aaaaaaagga tctcaagaag atcctttgat cttttctacg gggtctgacg ctcagtggaa
     9481 cgaaaactca cgttaaggga ttttggtcat gagattatca aaaaggatct tcacctagat
     9541 ccttttaaat taaaaatgaa gttttaaatc aatctaaagt atatatgagt aaacttggtc
     9601 tgacagttac caatgcttaa tcagtgaggc acctatctca gcgatctgtc tatttcgttc
     9661 atccatagtt gcctgactcc ccgtcgtgta gataactacg atacgggagg gcttaccatc
     9721 tggccccagt gctgcaatga taccgcgaga cccacgctca ccggctccag atttatcagc
     9781 aataaaccag ccagccggaa gggccgagcg cagaagtggt cctgcaactt tatccgcctc
     9841 catccagtct attaattgtt gccgggaagc tagagtaagt agttcgccag ttaatagttt
     9901 gcgcaacgtt gttgccattg ctacaggcat cgtggtgtca cgctcgtcgt ttggtatggc
     9961 ttcattcagc tccggttccc aacgatcaag gcgagttaca tgatccccca tgttgtgcaa
    10021 aaaagcggtt agctccttcg gtcctccgat cgttgtcaga agtaagttgg ccgcagtgtt
    10081 atcactcatg gttatggcag cactgcataa ttctcttact gtcatgccat ccgtaagatg
    10141 cttttctgtg actggtgagt actcaaccaa gtcattctga gaatagtgta tgcggcgacc
    10201 gagttgctct tgcccggcgt caatacggga taataccgcg ccacatagca gaactttaaa
    10261 agtgctcatc attggaaaac gttcttcggg gcgaaaactc tcaaggatct taccgctgtt
    10321 gagatccagt tcgatgtaac ccactcgtgc acccaactga tcttcagcat cttttacttt
    10381 caccagcgtt tctgggtgag caaaaacagg aaggcaaaat gccgcaaaaa agggaataag
    10441 ggcgacacgg aaatgttgaa tactcatact cttccttttt caatattatt gaagcattta
    10501 tcagggttat tgtctcatga gcggatacat atttgaatgt atttagaaaa ataaacaaat
    10561 aggggttccg cgcacatttc cccgaaaagt gccacactaa attgtaagcg ttaatatttt
    10621 gttaaaattc gcgttaaatt tttgttaaat cagctcattt tttaaccaat aggccgaaat
    10681 cggcaaaatc ccttataaat caaaagaata gaccgagata gggttgagtg ttgttccagt
    10741 ttggaacaag agtccactat taaagaacgt ggactccaac gtcaaagggc gaaaaaccgt
    10801 ctatcagggc gatggcccac tacgtgaacc atcaccctaa tcaagttttt tggggtcgag
    10861 gtgccgtaaa gcactaaatc ggaaccctaa agggagcccc cgatttagag cttgacgggg
    10921 aaagccggcg aacgtggcga gaaaggaagg gaagaaagcg aaaggagcgg gcgctagggc
    10981 gctggcaagt gtagcggtca cgctgcgcgt aaccaccaca cccgccgcgc ttaatgcgcc
    11041 gctacagggc gcgtcccatt cgccattcag gctgcgcaac tgttgggaag ggcgatcggt
    11101 gcgggcctct tcgctattac gccagctggc gaaaggggga tgtgctgcaa ggcgattaag
    11161 ttgggtaacg ccagggtttt cccagtcacg acgttgtaaa acgacggcca gtgagcgcgc
    11221 gtaatacgac tcactatagg gcgaattgga gctccaccgc ggtggcggcc g
//
```

**3XPcPS Plasmid**

```
LOCUS       Exported               13374 bp DNA     circular SYN 04-SEP-2021
DEFINITION  synthetic circular DNA
ACCESSION   .
VERSION     .
KEYWORDS    .
SOURCE      synthetic DNA construct
  ORGANISM  recombinant plasmid
REFERENCE   1  (bases 1 to 13374)
  AUTHORS   Gordon Wellman
  TITLE     Direct Submission
  JOURNAL   Exported Feb 13, 2022 from SnapGene 5.3.3
            https://www.snapgene.com
FEATURES             Location/Qualifiers
     source          1..13374
                     /organism="recombinant plasmid"
                     /mol_type="other DNA"
     promoter        8..274
                     /label=HSP70Ap promoter
                     /label=HSP70Ap
     promoter        275..466
                     /label=P-bTUB2
     misc_feature    361..366
                     /label=inserted ATANTT motif
     5'UTR           join(473..548,694..748)
                     /label=5UTR bTUB2
     misc_feature    495..500
                     /label=rebuild ATANTT from ATATT
     misc_feature    538
                     /label=XhoI site deleted T->A
     intron          549..693
                     /label=rbcS2 intron 1
     CDS             752..754
                     /codon_start=1
                     /label=Start
                     /translation="M"
     CDS             join(761..1198,1344..1844,1990..2463,2609..2857)
                     /codon_start=1
                     /label=Patchoulol synthase
                     /translation="ELYAQSVGVGAASRPLANFHPCVWGDKFIVYNPQSCQAGEREEAE
                     ELKVELKRELKEASDNYMRQLKMVDAIQRLGIDYLFVEDVDEALKNLFEMFDAFCKNNH
                     DMHATALSFRLLRQHGYRVSCEVFEKFKDGKDGFKVPNEDGAVAVLEFFEATHLRVHGE
                     DVLDNAFDFTRNYLESVYATLNDPTAKQVHNALNEFSFRRGLPRVEARKYISIYEQYAS
                     HHKGLLKLAKLDFNLVQALHRRELSEDSRWWKTLQVPTKLSFVRDRLVESYFWASGSYF
                     EPNYSVARMILAKGLAVLSLMDDVYDAYGTFEELQMFTDAIERWDASCLDKLPDYMKIV
                     YKALLDVFEEVDEELIKLGAPYRAYYGKEAMKYAARAYMEEAQWREQKHKPTTKEYMKL
                     ATKTCGYITLIILSCLGVEEGIVTKEAFDWVFSRPPFIEATLIIARLVNDITGHEFEKK
                     REHVRTAVECYMEEHKVGKQEVVSEFYNQMESAWKDINEGFLRPVEFPIPLLYLILNSV
                     RTLEVIYKEGDSYTHVGPAMQNIIKQLYLHPVPYGSG"
     intron          1199..1343
                     /label=rbcS2 intron 1
     intron          1845..1989
                     /label=rbcS2 intron 1
     intron          2464..2608
                     /label=rbcS2 intron 1
     CDS             join(2864..3301,3447..3947,4093..4566,4712..4960)
                     /codon_start=1
                     /label=Patchoulol synthase
                     /translation="ELYAQSVGVGAASRPLANFHPCVWGDKFIVYNPQSCQAGEREEAE
                     ELKVELKRELKEASDNYMRQLKMVDAIQRLGIDYLFVEDVDEALKNLFEMFDAFCKNNH
                     DMHATALSFRLLRQHGYRVSCEVFEKFKDGKDGFKVPNEDGAVAVLEFFEATHLRVHGE
                     DVLDNAFDFTRNYLESVYATLNDPTAKQVHNALNEFSFRRGLPRVEARKYISIYEQYAS
                     HHKGLLKLAKLDFNLVQALHRRELSEDSRWWKTLQVPTKLSFVRDRLVESYFWASGSYF
                     EPNYSVARMILAKGLAVLSLMDDVYDAYGTFEELQMFTDAIERWDASCLDKLPDYMKIV
                     YKALLDVFEEVDEELIKLGAPYRAYYGKEAMKYAARAYMEEAQWREQKHKPTTKEYMKL
                     ATKTCGYITLIILSCLGVEEGIVTKEAFDWVFSRPPFIEATLIIARLVNDITGHEFEKK
                     REHVRTAVECYMEEHKVGKQEVVSEFYNQMESAWKDINEGFLRPVEFPIPLLYLILNSV
                     RTLEVIYKEGDSYTHVGPAMQNIIKQLYLHPVPYGSG"
     intron          3302..3446
                     /label=rbcS2 intron 1
     intron          3948..4092
                     /label=rbcS2 intron 1
     intron          4567..4711
                     /label=rbcS2 intron 1
     CDS             join(4967..5404,5550..6050,6196..6669,6815..7063)
                     /codon_start=1
                     /label=Patchoulol synthase
                     /translation="ELYAQSVGVGAASRPLANFHPCVWGDKFIVYNPQSCQAGEREEAE
                     ELKVELKRELKEASDNYMRQLKMVDAIQRLGIDYLFVEDVDEALKNLFEMFDAFCKNNH
                     DMHATALSFRLLRQHGYRVSCEVFEKFKDGKDGFKVPNEDGAVAVLEFFEATHLRVHGE
                     DVLDNAFDFTRNYLESVYATLNDPTAKQVHNALNEFSFRRGLPRVEARKYISIYEQYAS
                     HHKGLLKLAKLDFNLVQALHRRELSEDSRWWKTLQVPTKLSFVRDRLVESYFWASGSYF
                     EPNYSVARMILAKGLAVLSLMDDVYDAYGTFEELQMFTDAIERWDASCLDKLPDYMKIV
                     YKALLDVFEEVDEELIKLGAPYRAYYGKEAMKYAARAYMEEAQWREQKHKPTTKEYMKL
                     ATKTCGYITLIILSCLGVEEGIVTKEAFDWVFSRPPFIEATLIIARLVNDITGHEFEKK
                     REHVRTAVECYMEEHKVGKQEVVSEFYNQMESAWKDINEGFLRPVEFPIPLLYLILNSV
                     RTLEVIYKEGDSYTHVGPAMQNIIKQLYLHPVPYGSG"
     intron          5405..5549
                     /label=rbcS2 intron 1
     intron          6051..6195
                     /label=rbcS2 intron 1
     intron          6670..6814
                     /label=rbcS2 intron 1
     CDS             7076..7093
                     /codon_start=1
                     /label=GSGS-Linker
                     /translation="GSGSGS"
     CDS             join(7094..7292,7438..7754,7900..8097)
                     /codon_start=1
                     /label=mVenus
                     /translation="VSKGEELFTGVVPILVELDGDVNGHKFSVSGEGEGDATYGKLTLK
                     LICTTGKLPVPWPTLVTTLGYGLQCFARYPDHMKQHDFFKSAMPEGYVQERTIFFKDDG
                     NYKTRAEVKFEGDTLVNRIELKGIDFKEDGNILGHKLEYNYNSHNVYITADKQKNGIKA
                     NFKIRHNIEDGGVQLADHYQQNTPIGDGPVLLPDNHYLSYQSKLSKDPNEKRDHMVLLE
                     FVTAAGITLGMDELYK"
     intron          7293..7437
                     /label=rbcS2 intron 1
     intron          7755..7899
                     /label=rbcS2 intron 1
     CDS             8098..8115
                     /codon_start=1
                     /label=GSGS-Linker
                     /translation="GSGSGS"
     CDS             join(8128..8146,8476..8477)
                     /codon_start=1
                     /label=GSGS-Linker
                     /translation="GSGSGSG"
     intron          8147..8475
                     /label=rbcS2 intron 2
     CDS             8478..8501
                     /codon_start=1
                     /product="peptide that binds Strep-Tactin(R), an engineered
                     form of streptavidin"
                     /label=Strep-Tag II
                     /translation="WSHPQFEK"
     misc_feature    8505..8738
                     /label=3'UTR
     misc_feature    8739..8750
                     /label=MCS
     promoter        8751..9017
                     /label=HSP70Ap
                     /label=HSP70Ap(1)
     promoter        join(9024..9252,9398..9399)
                     /label=RBCS2p promoter
                     /label=RBCS2p
     intron          9253..9397
                     /label=rbcS2 intron 1
     CDS             9440..10243
                     /codon_start=1
                     /label=APHVIII
                     /translation="MDDALRALRGRYPGCEWVVVEDGASGAGVYRLRGGGRELFVKVAA
                     LGAGVGLLGEAERLVWLAEVGIPVPRVVEGGGDERVAWLVTEAVPGRPASARWPREQRL
                     DVAVALAGLARSLHALDWERCPFDRSLAVTVPQAARAVAEGSVDLEDLDEERKGWSGER
                     LLAELERTRPADEDLAVCHGDLCPDNVLLDPRTCEVTGLIDVGRVGRADRHSDLALVLR
                     ELAHEEDPWFGPECSAAFLREYGRGWDGAVSEEKLAFYRLLDEFF"
     misc_feature    10253..10491
                     /label=3'UTR
                     /label=3'UTR(1)
     rep_origin      complement(10951..11539)
                     /direction=LEFT
                     /label=ori
                     /note="high-copy-number ColE1/pMB1/pBR322/pUC origin of
                     replication"
     CDS             complement(11710..12570)
                     /codon_start=1
                     /gene="bla"
                     /product="beta-lactamase"
                     /label=AmpR
                     /note="confers resistance to ampicillin, carbenicillin, and
                     related antibiotics"
                     /translation="MSIQHFRVALIPFFAAFCLPVFAHPETLVKVKDAEDQLGARVGYI
                     ELDLNSGKILESFRPEERFPMMSTFKVLLCGAVLSRIDAGQEQLGRRIHYSQNDLVEYS
                     PVTEKHLTDGMTVRELCSAAITMSDNTAANLLLTTIGGPKELTAFLHNMGDHVTRLDRW
                     EPELNEAIPNDERDTTMPVAMATTLRKLLTGELLTLASRQQLIDWMEADKVAGPLLRSA
                     LPAGWFIADKSGAGERGSRGIIAALGPDGKPSRIVVIYTTGSQATMDERNRQIAEIGAS
                     LIKHW"
     promoter        complement(12571..12675)
                     /gene="bla"
                     /label=AmpR promoter
     rep_origin      complement(12702..13157)
                     /direction=LEFT
                     /label=f1 ori
                     /note="f1 bacteriophage origin of replication; arrow
                     indicates direction of (+) strand synthesis"
ORIGIN
        1 ctctagagct gaggcttgac atgattggtg cgtatgtttg tatgaagcta caggactgat
       61 ttggcgggct atgagggcgg gggaagctct ggaagggccg cgatggggcg cgcggcgtcc
      121 agaaggcgcc atacggcccg ctggcggcac ccatccggta taaaagcccg cgaccccgaa
      181 cggtgacctc cactttcagc gacaaacgag cacttataca tacgcgacta ttctgccgct
      241 atacataacc actcagctag cttaagatcc catcctggca ctttcttgcg ctatgacact
      301 tccagcaaaa ggtagggcgg gctgcgagac ggcttcccgg cgctgcatgc aacaccgatg
      361 atacttatgc ttcgaccccc cgaagctcct tcggggctgc atgggcgctc cgatgccgct
      421 ccagggcgag cgctgtttaa atagccaggc ccccgactgc aaagacaccg gtattatagc
      481 gagctaccaa agccatactt caaacaccta gatcactacc acttctacac aggccacacg
      541 agcttgtggt gagtcgacga gcaagcccgg cggatcaggc agcgtgcttg cagatttgac
      601 ttgcaacgcc cgcattgtgt cgacgaaggc ttttggctcc tctgtcgctg tctcaagcag
      661 catctaaccc tgcgtcgccg tttccatttg cagatcgcac tccgctaagg gggcgcctct
      721 tcctcttcgt ttcagtcaca acccgcaaca tatgggatcc gagctgtacg cccagagcgt
      781 gggcgtgggc gccgccagcc gccccctggc caacttccac ccctgcgtgt ggggcgacaa
      841 gttcatcgtg tacaaccccc agagctgcca ggccggcgag cgcgaggagg ccgaggagct
      901 gaaggtggag ctgaagcgcg agctgaagga ggccagcgac aactacatgc gccagctgaa
      961 gatggtggac gccatccagc gcctgggcat cgactacctg ttcgtggagg acgtggacga
     1021 ggccctgaag aacctgttcg agatgttcga cgccttctgc aagaacaacc acgacatgca
     1081 cgccaccgcc ctgagcttcc gcctgctgcg ccagcacggc taccgcgtgt cctgcgaggt
     1141 gttcgagaag ttcaaggacg gcaaggacgg cttcaaggtg cccaacgagg acggcgcggt
     1201 gagtcgacga gcaagcccgg cggatcaggc agcgtgcttg cagatttgac ttgcaacgcc
     1261 cgcattgtgt cgacgaaggc ttttggctcc tctgtcgctg tctcaagcag catctaaccc
     1321 tgcgtcgccg tttccatttg caggtggcgg tgctggagtt cttcgaggcc acccacctgc
     1381 gcgtgcacgg cgaggacgtg ctggacaacg ccttcgactt cacccgcaac tacctggaga
     1441 gcgtgtacgc caccctgaac gaccccaccg ccaagcaggt gcacaacgcg ctgaacgagt
     1501 tctccttccg ccgcggcctg ccccgcgtgg aggcccgcaa gtacatcagc atctacgagc
     1561 agtacgccag ccaccacaag ggcctgctga agctggccaa gctggacttc aacctggtgc
     1621 aggcgctgca ccgccgcgag ctgtccgagg acagccgctg gtggaagacc ctgcaggtgc
     1681 ccaccaagct gagcttcgtg cgcgaccgcc tggtggagag ctacttctgg gccagcggca
     1741 gctacttcga gcccaactac agcgtggccc gcatgatcct ggcgaagggc ctggccgtgc
     1801 tgagcctgat ggacgacgtg tacgacgcct acggcacctt cgaggtgagt cgacgagcaa
     1861 gcccggcgga tcaggcagcg tgcttgcaga tttgacttgc aacgcccgca ttgtgtcgac
     1921 gaaggctttt ggctcctctg tcgctgtctc aagcagcatc taaccctgcg tcgccgtttc
     1981 catttgcagg agctgcagat gttcaccgac gcgatcgagc gctgggacgc cagctgcctg
     2041 gacaagctgc ccgactacat gaagatcgtg tacaaggccc tgctggacgt gttcgaggag
     2101 gtggacgagg agctgatcaa gctgggcgcc ccctaccgcg cctactacgg caaggaggcc
     2161 atgaagtacg ccgcccgcgc ctacatggag gaggcccagt ggcgcgagca gaagcacaag
     2221 cccaccacca aggagtacat gaagctggcg accaagacct gcggctacat caccctgatc
     2281 atcctgtcct gcctgggcgt ggaggagggc atcgtgacca aggaggcgtt cgactgggtg
     2341 ttcagccgcc cgccgttcat cgaggcgacc ctgatcatcg cccgcctggt caacgacatc
     2401 accggccacg agttcgagaa gaagcgcgag cacgtgcgca ccgccgtgga gtgctacatg
     2461 gaggtgagtc gacgagcaag cccggcggat caggcagcgt gcttgcagat ttgacttgca
     2521 acgcccgcat tgtgtcgacg aaggcttttg gctcctctgt cgctgtctca agcagcatct
     2581 aaccctgcgt cgccgtttcc atttgcagga gcacaaggtg ggcaagcagg aggtggtgtc
     2641 cgagttctac aaccagatgg agagcgcctg gaaggacatc aacgagggct tcctgcgccc
     2701 cgtggagttc cccatccccc tgctgtacct gatcctgaac agcgtgcgca ccctggaggt
     2761 gatctacaag gagggcgaca gctacaccca cgtgggcccg gccatgcaga acatcatcaa
     2821 gcagctgtac ctgcaccccg tgccctacgg cagcggcaga tccgagctgt acgcccagag
     2881 cgtgggcgtg ggcgccgcca gccgccccct ggccaacttc cacccctgcg tgtggggcga
     2941 caagttcatc gtgtacaacc cccagagctg ccaggccggc gagcgcgagg aggccgagga
     3001 gctgaaggtg gagctgaagc gcgagctgaa ggaggccagc gacaactaca tgcgccagct
     3061 gaagatggtg gacgccatcc agcgcctggg catcgactac ctgttcgtgg aggacgtgga
     3121 cgaggccctg aagaacctgt tcgagatgtt cgacgccttc tgcaagaaca accacgacat
     3181 gcacgccacc gccctgagct tccgcctgct gcgccagcac ggctaccgcg tgtcctgcga
     3241 ggtgttcgag aagttcaagg acggcaagga cggcttcaag gtgcccaacg aggacggcgc
     3301 ggtgagtcga cgagcaagcc cggcggatca ggcagcgtgc ttgcagattt gacttgcaac
     3361 gcccgcattg tgtcgacgaa ggcttttggc tcctctgtcg ctgtctcaag cagcatctaa
     3421 ccctgcgtcg ccgtttccat ttgcaggtgg cggtgctgga gttcttcgag gccacccacc
     3481 tgcgcgtgca cggcgaggac gtgctggaca acgccttcga cttcacccgc aactacctgg
     3541 agagcgtgta cgccaccctg aacgacccca ccgccaagca ggtgcacaac gcgctgaacg
     3601 agttctcctt ccgccgcggc ctgccccgcg tggaggcccg caagtacatc agcatctacg
     3661 agcagtacgc cagccaccac aagggcctgc tgaagctggc caagctggac ttcaacctgg
     3721 tgcaggcgct gcaccgccgc gagctgtccg aggacagccg ctggtggaag accctgcagg
     3781 tgcccaccaa gctgagcttc gtgcgcgacc gcctggtgga gagctacttc tgggccagcg
     3841 gcagctactt cgagcccaac tacagcgtgg cccgcatgat cctggcgaag ggcctggccg
     3901 tgctgagcct gatggacgac gtgtacgacg cctacggcac cttcgaggtg agtcgacgag
     3961 caagcccggc ggatcaggca gcgtgcttgc agatttgact tgcaacgccc gcattgtgtc
     4021 gacgaaggct tttggctcct ctgtcgctgt ctcaagcagc atctaaccct gcgtcgccgt
     4081 ttccatttgc aggagctgca gatgttcacc gacgcgatcg agcgctggga cgccagctgc
     4141 ctggacaagc tgcccgacta catgaagatc gtgtacaagg ccctgctgga cgtgttcgag
     4201 gaggtggacg aggagctgat caagctgggc gccccctacc gcgcctacta cggcaaggag
     4261 gccatgaagt acgccgcccg cgcctacatg gaggaggccc agtggcgcga gcagaagcac
     4321 aagcccacca ccaaggagta catgaagctg gcgaccaaga cctgcggcta catcaccctg
     4381 atcatcctgt cctgcctggg cgtggaggag ggcatcgtga ccaaggaggc gttcgactgg
     4441 gtgttcagcc gcccgccgtt catcgaggcg accctgatca tcgcccgcct ggtcaacgac
     4501 atcaccggcc acgagttcga gaagaagcgc gagcacgtgc gcaccgccgt ggagtgctac
     4561 atggaggtga gtcgacgagc aagcccggcg gatcaggcag cgtgcttgca gatttgactt
     4621 gcaacgcccg cattgtgtcg acgaaggctt ttggctcctc tgtcgctgtc tcaagcagca
     4681 tctaaccctg cgtcgccgtt tccatttgca ggagcacaag gtgggcaagc aggaggtggt
     4741 gtccgagttc tacaaccaga tggagagcgc ctggaaggac atcaacgagg gcttcctgcg
     4801 ccccgtggag ttccccatcc ccctgctgta cctgatcctg aacagcgtgc gcaccctgga
     4861 ggtgatctac aaggagggcg acagctacac ccacgtgggc ccggccatgc agaacatcat
     4921 caagcagctg tacctgcacc ccgtgcccta cggcagcggc agatccgagc tgtacgccca
     4981 gagcgtgggc gtgggcgccg ccagccgccc cctggccaac ttccacccct gcgtgtgggg
     5041 cgacaagttc atcgtgtaca acccccagag ctgccaggcc ggcgagcgcg aggaggccga
     5101 ggagctgaag gtggagctga agcgcgagct gaaggaggcc agcgacaact acatgcgcca
     5161 gctgaagatg gtggacgcca tccagcgcct gggcatcgac tacctgttcg tggaggacgt
     5221 ggacgaggcc ctgaagaacc tgttcgagat gttcgacgcc ttctgcaaga acaaccacga
     5281 catgcacgcc accgccctga gcttccgcct gctgcgccag cacggctacc gcgtgtcctg
     5341 cgaggtgttc gagaagttca aggacggcaa ggacggcttc aaggtgccca acgaggacgg
     5401 cgcggtgagt cgacgagcaa gcccggcgga tcaggcagcg tgcttgcaga tttgacttgc
     5461 aacgcccgca ttgtgtcgac gaaggctttt ggctcctctg tcgctgtctc aagcagcatc
     5521 taaccctgcg tcgccgtttc catttgcagg tggcggtgct ggagttcttc gaggccaccc
     5581 acctgcgcgt gcacggcgag gacgtgctgg acaacgcctt cgacttcacc cgcaactacc
     5641 tggagagcgt gtacgccacc ctgaacgacc ccaccgccaa gcaggtgcac aacgcgctga
     5701 acgagttctc cttccgccgc ggcctgcccc gcgtggaggc ccgcaagtac atcagcatct
     5761 acgagcagta cgccagccac cacaagggcc tgctgaagct ggccaagctg gacttcaacc
     5821 tggtgcaggc gctgcaccgc cgcgagctgt ccgaggacag ccgctggtgg aagaccctgc
     5881 aggtgcccac caagctgagc ttcgtgcgcg accgcctggt ggagagctac ttctgggcca
     5941 gcggcagcta cttcgagccc aactacagcg tggcccgcat gatcctggcg aagggcctgg
     6001 ccgtgctgag cctgatggac gacgtgtacg acgcctacgg caccttcgag gtgagtcgac
     6061 gagcaagccc ggcggatcag gcagcgtgct tgcagatttg acttgcaacg cccgcattgt
     6121 gtcgacgaag gcttttggct cctctgtcgc tgtctcaagc agcatctaac cctgcgtcgc
     6181 cgtttccatt tgcaggagct gcagatgttc accgacgcga tcgagcgctg ggacgccagc
     6241 tgcctggaca agctgcccga ctacatgaag atcgtgtaca aggccctgct ggacgtgttc
     6301 gaggaggtgg acgaggagct gatcaagctg ggcgccccct accgcgccta ctacggcaag
     6361 gaggccatga agtacgccgc ccgcgcctac atggaggagg cccagtggcg cgagcagaag
     6421 cacaagccca ccaccaagga gtacatgaag ctggcgacca agacctgcgg ctacatcacc
     6481 ctgatcatcc tgtcctgcct gggcgtggag gagggcatcg tgaccaagga ggcgttcgac
     6541 tgggtgttca gccgcccgcc gttcatcgag gcgaccctga tcatcgcccg cctggtcaac
     6601 gacatcaccg gccacgagtt cgagaagaag cgcgagcacg tgcgcaccgc cgtggagtgc
     6661 tacatggagg tgagtcgacg agcaagcccg gcggatcagg cagcgtgctt gcagatttga
     6721 cttgcaacgc ccgcattgtg tcgacgaagg cttttggctc ctctgtcgct gtctcaagca
     6781 gcatctaacc ctgcgtcgcc gtttccattt gcaggagcac aaggtgggca agcaggaggt
     6841 ggtgtccgag ttctacaacc agatggagag cgcctggaag gacatcaacg agggcttcct
     6901 gcgccccgtg gagttcccca tccccctgct gtacctgatc ctgaacagcg tgcgcaccct
     6961 ggaggtgatc tacaaggagg gcgacagcta cacccacgtg ggcccggcca tgcagaacat
     7021 catcaagcag ctgtacctgc accccgtgcc ctacggcagc ggcagatctg acgtcggcag
     7081 cggcagcggc agcgtgagca agggcgagga gctgttcacc ggcgtggtgc ccatcctggt
     7141 ggagctggac ggcgacgtga acggccacaa gttcagcgtg agcggcgagg gcgagggcga
     7201 cgccacctac ggcaagctga ccctgaagct gatctgcacc accggcaagc tgcccgtgcc
     7261 ctggcccacc ctggtgacca ccctgggcta cggtgagtcg acgagcaagc ccggcggatc
     7321 aggcagcgtg cttgcagatt tgacttgcaa cgcccgcatt gtgtcgacga aggcttttgg
     7381 ctcctctgtc gctgtctcaa gcagcatcta accctgcgtc gccgtttcca tttgcaggcc
     7441 tgcagtgctt cgcccgctac cccgaccaca tgaagcagca cgacttcttc aagagcgcca
     7501 tgcccgaggg ctacgtgcag gagcgcacca tcttcttcaa ggacgacggt aactacaaga
     7561 cccgcgccga ggtgaagttc gagggcgaca ccctggtgaa ccgcatcgag ctgaagggca
     7621 tcgacttcaa ggaggacggc aacatcctgg gccacaagct ggagtacaac tacaacagcc
     7681 acaacgtgta catcaccgcc gacaagcaga agaacggcat caaggccaac ttcaagatcc
     7741 gccacaacat cgaggtgagt cgacgagcaa gcccggcgga tcaggcagcg tgcttgcaga
     7801 tttgacttgc aacgcccgca ttgtgtcgac gaaggctttt ggctcctctg tcgctgtctc
     7861 aagcagcatc taaccctgcg tcgccgtttc catttgcagg acggcggcgt gcagctggcc
     7921 gaccactacc agcagaacac ccccatcggc gacggccccg tgctgctgcc cgacaaccac
     7981 tacctgagct accagagcaa gctgagcaag gaccccaacg agaagcgcga ccacatggtg
     8041 ctgctggagt tcgtgaccgc cgccggcatc accctgggca tggacgagct gtacaagggc
     8101 agcggcagcg gcagcgatat cgaattcggc agcggcagcg gctcaggtga gcttgcgggg
     8161 ttgcgagcaa cactccagca acgaacagtg cccaagtcag gaatctgcag tcagcctggg
     8221 ctttcggcgg ctttttcttg ggcaaacagc ttgcactcat gccagcgcgg cttgtccagc
     8281 ctcacttgag ctttccagct gctaccagcc gggctatacg acagcgacag agccatagcg
     8341 tggaatcact tatttgggtt gccgaagtag cggtcggagc gtgagttctt ggtcaagccg
     8401 ccccttatcc ggttcctgtc cgtgtctttg tccctcgttc acccttcgcg gcacccttca
     8461 tccccttgct tgcaggttgg agccacccgc agttcgagaa gtaaccgctc cgtgtaaatg
     8521 gaggcgctcg ttgatctgag ccttgccccc tgacgaacgg cggtggatgg aagatactgc
     8581 tctcaagtgc tgaagcggta gcttagctcc ccgtttcgtg ctgatcagtc tttttcaaca
     8641 cgtaaaaagc ggaggagttt tgcaattttg ttggttgtaa cgatcctccg ttgattttgg
     8701 cctctttctc catgggcggg ctgggcgtat ttgaagcgac tagtacgcgt gctgaggctt
     8761 gacatgattg gtgcgtatgt ttgtatgaag ctacaggact gatttggcgg gctatgaggg
     8821 cgggggaagc tctggaaggg ccgcgatggg gcgcgcggcg tccagaaggc gccatacggc
     8881 ccgctggcgg cacccatccg gtataaaagc ccgcgacccc gaacggtgac ctccactttc
     8941 agcgacaaac gagcacttat acatacgcga ctattctgcc gctatacata accactcagc
     9001 tagcttaaga tcccatccct agggcatgcc gggcgcgcca gaaggagcgc agccaaacca
     9061 ggatgatgtt tgatggggta tttgagcact tgcaaccctt atccggaagc cccctggccc
     9121 acaaaggcta ggcgccaatg caagcagttc gcatgcagcc cctggagcgg tgccctcctg
     9181 ataaaccggc cagggggcct atgttcttta cttttttaca agagaagtca ctcaacatct
     9241 taaaatggcc aggtgagtcg acgagcaagc ccggcggatc aggcagcgtg cttgcagatt
     9301 tgacttgcaa cgcccgcatt gtgtcgacga aggcttttgg ctcctctgtc gctgtctcaa
     9361 gcagcatcta accctgcgtc gccgtttcca tttgcaggaa gcttactccg ccctccccgg
     9421 tgctgaagaa tttcgaagca tggacgatgc gttgcgtgca ctgcggggtc ggtatcccgg
     9481 ttgtgagtgg gttgttgtgg aggatggggc ctcgggggct ggtgtttatc ggcttcgggg
     9541 tggtgggcgg gagttgtttg tcaaggtggc agctctgggg gccggggtgg gcttgttggg
     9601 tgaggctgag cggctggtgt ggttggcgga ggtggggatt cccgtacctc gtgttgtgga
     9661 gggtggtggg gacgagaggg tcgcctggtt ggtcaccgaa gcggttccgg ggcgtccggc
     9721 cagtgcgcgg tggccgcggg agcagcggct ggacgtggcg gtggcgctcg cggggctcgc
     9781 tcgttcgctg cacgcgctgg actgggagcg gtgtccgttc gatcgcagtc tcgcggtgac
     9841 ggtgccgcag gcggcccgtg ctgtcgctga agggagcgtc gacttggagg atctggacga
     9901 ggagcggaag gggtggtcgg gggagcggct tctcgccgag ctggagcgga ctcggcctgc
     9961 ggacgaggat ctggcggttt gccacggtga cctgtgcccg gacaacgtgc tgctcgaccc
    10021 tcgtacctgc gaggtgaccg ggctgatcga cgtggggcgg gtcggccgtg cggaccggca
    10081 ctccgatctc gcgctggtgc tgcgcgagct ggcccacgag gaggacccgt ggttcgggcc
    10141 ggagtgttcc gcggcgttcc tgcgggagta cgggcgcggg tgggatgggg cggtatcgga
    10201 ggaaaagctg gcgttttacc ggctgttgga cgagttcttc tgactcgagt gaccgctccg
    10261 tgtaaatgga ggcgctcgtt gatctgagcc ttgccccctg acgaacggcg gtggatggaa
    10321 gatactgctc tcaagtgctg aagcggtagc ttagctcccc gtttcgtgct gatcagtctt
    10381 tttcaacacg taaaaagcgg aggagttttg caattttgtt ggttgtaacg atcctccgtt
    10441 gattttggcc tctttctcca tgggcgggct gggcgtattt gaagcggacc cggtacccag
    10501 cttttgttcc ctttagtgag ggttaattgc gcgcttggcg taatcatggt catagctgtt
    10561 tcctgtgtga aattgttatc cgctcacaat tccacacaac atacgagccg gaagcataaa
    10621 gtgtaaagcc tggggtgcct aatgagtgag ctaactcaca ttaattgcgt tgcgctcact
    10681 gcccgctttc cagtcgggaa acctgtcgtg ccagctgcat taatgaatcg gccaacgcgc
    10741 ggggagaggc ggtttgcgta ttgggcgctc ttccgcttcc tcgctcactg actcgctgcg
    10801 ctcggtcgtt cggctgcggc gagcggtatc agctcactca aaggcggtaa tacggttatc
    10861 cacagaatca ggggataacg caggaaagaa catgtgagca aaaggccagc aaaaggccag
    10921 gaaccgtaaa aaggccgcgt tgctggcgtt tttccatagg ctccgccccc ctgacgagca
    10981 tcacaaaaat cgacgctcaa gtcagaggtg gcgaaacccg acaggactat aaagatacca
    11041 ggcgtttccc cctggaagct ccctcgtgcg ctctcctgtt ccgaccctgc cgcttaccgg
    11101 atacctgtcc gcctttctcc cttcgggaag cgtggcgctt tctcatagct cacgctgtag
    11161 gtatctcagt tcggtgtagg tcgttcgctc caagctgggc tgtgtgcacg aaccccccgt
    11221 tcagcccgac cgctgcgcct tatccggtaa ctatcgtctt gagtccaacc cggtaagaca
    11281 cgacttatcg ccactggcag cagccactgg taacaggatt agcagagcga ggtatgtagg
    11341 cggtgctaca gagttcttga agtggtggcc taactacggc tacactagaa ggacagtatt
    11401 tggtatctgc gctctgctga agccagttac cttcggaaaa agagttggta gctcttgatc
    11461 cggcaaacaa accaccgctg gtagcggtgg tttttttgtt tgcaagcagc agattacgcg
    11521 cagaaaaaaa ggatctcaag aagatccttt gatcttttct acggggtctg acgctcagtg
    11581 gaacgaaaac tcacgttaag ggattttggt catgagatta tcaaaaagga tcttcaccta
    11641 gatcctttta aattaaaaat gaagttttaa atcaatctaa agtatatatg agtaaacttg
    11701 gtctgacagt taccaatgct taatcagtga ggcacctatc tcagcgatct gtctatttcg
    11761 ttcatccata gttgcctgac tccccgtcgt gtagataact acgatacggg agggcttacc
    11821 atctggcccc agtgctgcaa tgataccgcg agacccacgc tcaccggctc cagatttatc
    11881 agcaataaac cagccagccg gaagggccga gcgcagaagt ggtcctgcaa ctttatccgc
    11941 ctccatccag tctattaatt gttgccggga agctagagta agtagttcgc cagttaatag
    12001 tttgcgcaac gttgttgcca ttgctacagg catcgtggtg tcacgctcgt cgtttggtat
    12061 ggcttcattc agctccggtt cccaacgatc aaggcgagtt acatgatccc ccatgttgtg
    12121 caaaaaagcg gttagctcct tcggtcctcc gatcgttgtc agaagtaagt tggccgcagt
    12181 gttatcactc atggttatgg cagcactgca taattctctt actgtcatgc catccgtaag
    12241 atgcttttct gtgactggtg agtactcaac caagtcattc tgagaatagt gtatgcggcg
    12301 accgagttgc tcttgcccgg cgtcaatacg ggataatacc gcgccacata gcagaacttt
    12361 aaaagtgctc atcattggaa aacgttcttc ggggcgaaaa ctctcaagga tcttaccgct
    12421 gttgagatcc agttcgatgt aacccactcg tgcacccaac tgatcttcag catcttttac
    12481 tttcaccagc gtttctgggt gagcaaaaac aggaaggcaa aatgccgcaa aaaagggaat
    12541 aagggcgaca cggaaatgtt gaatactcat actcttcctt tttcaatatt attgaagcat
    12601 ttatcagggt tattgtctca tgagcggata catatttgaa tgtatttaga aaaataaaca
    12661 aataggggtt ccgcgcacat ttccccgaaa agtgccacac taaattgtaa gcgttaatat
    12721 tttgttaaaa ttcgcgttaa atttttgtta aatcagctca ttttttaacc aataggccga
    12781 aatcggcaaa atcccttata aatcaaaaga atagaccgag atagggttga gtgttgttcc
    12841 agtttggaac aagagtccac tattaaagaa cgtggactcc aacgtcaaag ggcgaaaaac
    12901 cgtctatcag ggcgatggcc cactacgtga accatcaccc taatcaagtt ttttggggtc
    12961 gaggtgccgt aaagcactaa atcggaaccc taaagggagc ccccgattta gagcttgacg
    13021 gggaaagccg gcgaacgtgg cgagaaagga agggaagaaa gcgaaaggag cgggcgctag
    13081 ggcgctggca agtgtagcgg tcacgctgcg cgtaaccacc acacccgccg cgcttaatgc
    13141 gccgctacag ggcgcgtccc attcgccatt caggctgcgc aactgttggg aagggcgatc
    13201 ggtgcgggcc tcttcgctat tacgccagct ggcgaaaggg ggatgtgctg caaggcgatt
    13261 aagttgggta acgccagggt tttcccagtc acgacgttgt aaaacgacgg ccagtgagcg
    13321 cgcgtaatac gactcactat agggcgaatt ggagctccac cgcggtggcg gccg
//
```

**4XPcPS plasmid**

```
LOCUS       Exported               15477 bp DNA     circular SYN 04-SEP-2021
DEFINITION  synthetic circular DNA
ACCESSION   .
VERSION     .
KEYWORDS    .
SOURCE      synthetic DNA construct
  ORGANISM  recombinant plasmid
REFERENCE   1  (bases 1 to 15477)
  AUTHORS   Gordon Wellman
  TITLE     Direct Submission
  JOURNAL   Exported Feb 13, 2022 from SnapGene 5.3.3
            https://www.snapgene.com
FEATURES             Location/Qualifiers
     source          1..15477
                     /organism="recombinant plasmid"
                     /mol_type="other DNA"
     promoter        8..274
                     /label=HSP70Ap promoter
                     /label=HSP70Ap
     promoter        275..466
                     /label=P-bTUB2
     misc_feature    361..366
                     /label=inserted ATANTT motif
     5'UTR           join(473..548,694..748)
                     /label=5UTR bTUB2
     misc_feature    495..500
                     /label=rebuild ATANTT from ATATT
     misc_feature    538
                     /label=XhoI site deleted T->A
     intron          549..693
                     /label=rbcS2 intron 1
     CDS             752..754
                     /codon_start=1
                     /label=Start
                     /translation="M"
     CDS             join(761..1198,1344..1844,1990..2463,2609..2857)
                     /codon_start=1
                     /label=Patchoulol synthase
                     /translation="ELYAQSVGVGAASRPLANFHPCVWGDKFIVYNPQSCQAGEREEAE
                     ELKVELKRELKEASDNYMRQLKMVDAIQRLGIDYLFVEDVDEALKNLFEMFDAFCKNNH
                     DMHATALSFRLLRQHGYRVSCEVFEKFKDGKDGFKVPNEDGAVAVLEFFEATHLRVHGE
                     DVLDNAFDFTRNYLESVYATLNDPTAKQVHNALNEFSFRRGLPRVEARKYISIYEQYAS
                     HHKGLLKLAKLDFNLVQALHRRELSEDSRWWKTLQVPTKLSFVRDRLVESYFWASGSYF
                     EPNYSVARMILAKGLAVLSLMDDVYDAYGTFEELQMFTDAIERWDASCLDKLPDYMKIV
                     YKALLDVFEEVDEELIKLGAPYRAYYGKEAMKYAARAYMEEAQWREQKHKPTTKEYMKL
                     ATKTCGYITLIILSCLGVEEGIVTKEAFDWVFSRPPFIEATLIIARLVNDITGHEFEKK
                     REHVRTAVECYMEEHKVGKQEVVSEFYNQMESAWKDINEGFLRPVEFPIPLLYLILNSV
                     RTLEVIYKEGDSYTHVGPAMQNIIKQLYLHPVPYGSG"
     intron          1199..1343
                     /label=rbcS2 intron 1
     intron          1845..1989
                     /label=rbcS2 intron 1
     intron          2464..2608
                     /label=rbcS2 intron 1
     CDS             join(2864..3301,3447..3947,4093..4566,4712..4960)
                     /codon_start=1
                     /label=Patchoulol synthase
                     /translation="ELYAQSVGVGAASRPLANFHPCVWGDKFIVYNPQSCQAGEREEAE
                     ELKVELKRELKEASDNYMRQLKMVDAIQRLGIDYLFVEDVDEALKNLFEMFDAFCKNNH
                     DMHATALSFRLLRQHGYRVSCEVFEKFKDGKDGFKVPNEDGAVAVLEFFEATHLRVHGE
                     DVLDNAFDFTRNYLESVYATLNDPTAKQVHNALNEFSFRRGLPRVEARKYISIYEQYAS
                     HHKGLLKLAKLDFNLVQALHRRELSEDSRWWKTLQVPTKLSFVRDRLVESYFWASGSYF
                     EPNYSVARMILAKGLAVLSLMDDVYDAYGTFEELQMFTDAIERWDASCLDKLPDYMKIV
                     YKALLDVFEEVDEELIKLGAPYRAYYGKEAMKYAARAYMEEAQWREQKHKPTTKEYMKL
                     ATKTCGYITLIILSCLGVEEGIVTKEAFDWVFSRPPFIEATLIIARLVNDITGHEFEKK
                     REHVRTAVECYMEEHKVGKQEVVSEFYNQMESAWKDINEGFLRPVEFPIPLLYLILNSV
                     RTLEVIYKEGDSYTHVGPAMQNIIKQLYLHPVPYGSG"
     intron          3302..3446
                     /label=rbcS2 intron 1
     intron          3948..4092
                     /label=rbcS2 intron 1
     intron          4567..4711
                     /label=rbcS2 intron 1
     CDS             join(4967..5404,5550..6050,6196..6669,6815..7063)
                     /codon_start=1
                     /label=Patchoulol synthase
                     /translation="ELYAQSVGVGAASRPLANFHPCVWGDKFIVYNPQSCQAGEREEAE
                     ELKVELKRELKEASDNYMRQLKMVDAIQRLGIDYLFVEDVDEALKNLFEMFDAFCKNNH
                     DMHATALSFRLLRQHGYRVSCEVFEKFKDGKDGFKVPNEDGAVAVLEFFEATHLRVHGE
                     DVLDNAFDFTRNYLESVYATLNDPTAKQVHNALNEFSFRRGLPRVEARKYISIYEQYAS
                     HHKGLLKLAKLDFNLVQALHRRELSEDSRWWKTLQVPTKLSFVRDRLVESYFWASGSYF
                     EPNYSVARMILAKGLAVLSLMDDVYDAYGTFEELQMFTDAIERWDASCLDKLPDYMKIV
                     YKALLDVFEEVDEELIKLGAPYRAYYGKEAMKYAARAYMEEAQWREQKHKPTTKEYMKL
                     ATKTCGYITLIILSCLGVEEGIVTKEAFDWVFSRPPFIEATLIIARLVNDITGHEFEKK
                     REHVRTAVECYMEEHKVGKQEVVSEFYNQMESAWKDINEGFLRPVEFPIPLLYLILNSV
                     RTLEVIYKEGDSYTHVGPAMQNIIKQLYLHPVPYGSG"
     intron          5405..5549
                     /label=rbcS2 intron 1
     intron          6051..6195
                     /label=rbcS2 intron 1
     intron          6670..6814
                     /label=rbcS2 intron 1
     CDS             join(7070..7507,7653..8153,8299..8772,8918..9166)
                     /codon_start=1
                     /label=Patchoulol synthase
                     /translation="ELYAQSVGVGAASRPLANFHPCVWGDKFIVYNPQSCQAGEREEAE
                     ELKVELKRELKEASDNYMRQLKMVDAIQRLGIDYLFVEDVDEALKNLFEMFDAFCKNNH
                     DMHATALSFRLLRQHGYRVSCEVFEKFKDGKDGFKVPNEDGAVAVLEFFEATHLRVHGE
                     DVLDNAFDFTRNYLESVYATLNDPTAKQVHNALNEFSFRRGLPRVEARKYISIYEQYAS
                     HHKGLLKLAKLDFNLVQALHRRELSEDSRWWKTLQVPTKLSFVRDRLVESYFWASGSYF
                     EPNYSVARMILAKGLAVLSLMDDVYDAYGTFEELQMFTDAIERWDASCLDKLPDYMKIV
                     YKALLDVFEEVDEELIKLGAPYRAYYGKEAMKYAARAYMEEAQWREQKHKPTTKEYMKL
                     ATKTCGYITLIILSCLGVEEGIVTKEAFDWVFSRPPFIEATLIIARLVNDITGHEFEKK
                     REHVRTAVECYMEEHKVGKQEVVSEFYNQMESAWKDINEGFLRPVEFPIPLLYLILNSV
                     RTLEVIYKEGDSYTHVGPAMQNIIKQLYLHPVPYGSG"
     intron          7508..7652
                     /label=rbcS2 intron 1
     intron          8154..8298
                     /label=rbcS2 intron 1
     intron          8773..8917
                     /label=rbcS2 intron 1
     CDS             9179..9196
                     /codon_start=1
                     /label=GSGS-Linker
                     /translation="GSGSGS"
     CDS             join(9197..9395,9541..9857,10003..10200)
                     /codon_start=1
                     /label=mVenus
                     /translation="VSKGEELFTGVVPILVELDGDVNGHKFSVSGEGEGDATYGKLTLK
                     LICTTGKLPVPWPTLVTTLGYGLQCFARYPDHMKQHDFFKSAMPEGYVQERTIFFKDDG
                     NYKTRAEVKFEGDTLVNRIELKGIDFKEDGNILGHKLEYNYNSHNVYITADKQKNGIKA
                     NFKIRHNIEDGGVQLADHYQQNTPIGDGPVLLPDNHYLSYQSKLSKDPNEKRDHMVLLE
                     FVTAAGITLGMDELYK"
     intron          9396..9540
                     /label=rbcS2 intron 1
     intron          9858..10002
                     /label=rbcS2 intron 1
     CDS             10201..10218
                     /codon_start=1
                     /label=GSGS-Linker
                     /translation="GSGSGS"
     CDS             join(10231..10249,10579..10580)
                     /codon_start=1
                     /label=GSGS-Linker
                     /translation="GSGSGSG"
     intron          10250..10578
                     /label=rbcS2 intron 2
     CDS             10581..10604
                     /codon_start=1
                     /product="peptide that binds Strep-Tactin(R), an engineered
                     form of streptavidin"
                     /label=Strep-Tag II
                     /translation="WSHPQFEK"
     misc_feature    10608..10841
                     /label=3'UTR
     misc_feature    10842..10853
                     /label=MCS
     promoter        10854..11120
                     /label=HSP70Ap
                     /label=HSP70Ap(1)
     promoter        join(11127..11355,11501..11502)
                     /label=RBCS2p promoter
                     /label=RBCS2p
     intron          11356..11500
                     /label=rbcS2 intron 1
     CDS             11543..12346
                     /codon_start=1
                     /label=APHVIII
                     /translation="MDDALRALRGRYPGCEWVVVEDGASGAGVYRLRGGGRELFVKVAA
                     LGAGVGLLGEAERLVWLAEVGIPVPRVVEGGGDERVAWLVTEAVPGRPASARWPREQRL
                     DVAVALAGLARSLHALDWERCPFDRSLAVTVPQAARAVAEGSVDLEDLDEERKGWSGER
                     LLAELERTRPADEDLAVCHGDLCPDNVLLDPRTCEVTGLIDVGRVGRADRHSDLALVLR
                     ELAHEEDPWFGPECSAAFLREYGRGWDGAVSEEKLAFYRLLDEFF"
     misc_feature    12356..12594
                     /label=3'UTR
                     /label=3'UTR(1)
     rep_origin      complement(13054..13642)
                     /direction=LEFT
                     /label=ori
                     /note="high-copy-number ColE1/pMB1/pBR322/pUC origin of
                     replication"
     CDS             complement(13813..14673)
                     /codon_start=1
                     /gene="bla"
                     /product="beta-lactamase"
                     /label=AmpR
                     /note="confers resistance to ampicillin, carbenicillin, and
                     related antibiotics"
                     /translation="MSIQHFRVALIPFFAAFCLPVFAHPETLVKVKDAEDQLGARVGYI
                     ELDLNSGKILESFRPEERFPMMSTFKVLLCGAVLSRIDAGQEQLGRRIHYSQNDLVEYS
                     PVTEKHLTDGMTVRELCSAAITMSDNTAANLLLTTIGGPKELTAFLHNMGDHVTRLDRW
                     EPELNEAIPNDERDTTMPVAMATTLRKLLTGELLTLASRQQLIDWMEADKVAGPLLRSA
                     LPAGWFIADKSGAGERGSRGIIAALGPDGKPSRIVVIYTTGSQATMDERNRQIAEIGAS
                     LIKHW"
     promoter        complement(14674..14778)
                     /gene="bla"
                     /label=AmpR promoter
     rep_origin      complement(14805..15260)
                     /direction=LEFT
                     /label=f1 ori
                     /note="f1 bacteriophage origin of replication; arrow
                     indicates direction of (+) strand synthesis"
ORIGIN
        1 ctctagagct gaggcttgac atgattggtg cgtatgtttg tatgaagcta caggactgat
       61 ttggcgggct atgagggcgg gggaagctct ggaagggccg cgatggggcg cgcggcgtcc
      121 agaaggcgcc atacggcccg ctggcggcac ccatccggta taaaagcccg cgaccccgaa
      181 cggtgacctc cactttcagc gacaaacgag cacttataca tacgcgacta ttctgccgct
      241 atacataacc actcagctag cttaagatcc catcctggca ctttcttgcg ctatgacact
      301 tccagcaaaa ggtagggcgg gctgcgagac ggcttcccgg cgctgcatgc aacaccgatg
      361 atacttatgc ttcgaccccc cgaagctcct tcggggctgc atgggcgctc cgatgccgct
      421 ccagggcgag cgctgtttaa atagccaggc ccccgactgc aaagacaccg gtattatagc
      481 gagctaccaa agccatactt caaacaccta gatcactacc acttctacac aggccacacg
      541 agcttgtggt gagtcgacga gcaagcccgg cggatcaggc agcgtgcttg cagatttgac
      601 ttgcaacgcc cgcattgtgt cgacgaaggc ttttggctcc tctgtcgctg tctcaagcag
      661 catctaaccc tgcgtcgccg tttccatttg cagatcgcac tccgctaagg gggcgcctct
      721 tcctcttcgt ttcagtcaca acccgcaaca tatgggatcc gagctgtacg cccagagcgt
      781 gggcgtgggc gccgccagcc gccccctggc caacttccac ccctgcgtgt ggggcgacaa
      841 gttcatcgtg tacaaccccc agagctgcca ggccggcgag cgcgaggagg ccgaggagct
      901 gaaggtggag ctgaagcgcg agctgaagga ggccagcgac aactacatgc gccagctgaa
      961 gatggtggac gccatccagc gcctgggcat cgactacctg ttcgtggagg acgtggacga
     1021 ggccctgaag aacctgttcg agatgttcga cgccttctgc aagaacaacc acgacatgca
     1081 cgccaccgcc ctgagcttcc gcctgctgcg ccagcacggc taccgcgtgt cctgcgaggt
     1141 gttcgagaag ttcaaggacg gcaaggacgg cttcaaggtg cccaacgagg acggcgcggt
     1201 gagtcgacga gcaagcccgg cggatcaggc agcgtgcttg cagatttgac ttgcaacgcc
     1261 cgcattgtgt cgacgaaggc ttttggctcc tctgtcgctg tctcaagcag catctaaccc
     1321 tgcgtcgccg tttccatttg caggtggcgg tgctggagtt cttcgaggcc acccacctgc
     1381 gcgtgcacgg cgaggacgtg ctggacaacg ccttcgactt cacccgcaac tacctggaga
     1441 gcgtgtacgc caccctgaac gaccccaccg ccaagcaggt gcacaacgcg ctgaacgagt
     1501 tctccttccg ccgcggcctg ccccgcgtgg aggcccgcaa gtacatcagc atctacgagc
     1561 agtacgccag ccaccacaag ggcctgctga agctggccaa gctggacttc aacctggtgc
     1621 aggcgctgca ccgccgcgag ctgtccgagg acagccgctg gtggaagacc ctgcaggtgc
     1681 ccaccaagct gagcttcgtg cgcgaccgcc tggtggagag ctacttctgg gccagcggca
     1741 gctacttcga gcccaactac agcgtggccc gcatgatcct ggcgaagggc ctggccgtgc
     1801 tgagcctgat ggacgacgtg tacgacgcct acggcacctt cgaggtgagt cgacgagcaa
     1861 gcccggcgga tcaggcagcg tgcttgcaga tttgacttgc aacgcccgca ttgtgtcgac
     1921 gaaggctttt ggctcctctg tcgctgtctc aagcagcatc taaccctgcg tcgccgtttc
     1981 catttgcagg agctgcagat gttcaccgac gcgatcgagc gctgggacgc cagctgcctg
     2041 gacaagctgc ccgactacat gaagatcgtg tacaaggccc tgctggacgt gttcgaggag
     2101 gtggacgagg agctgatcaa gctgggcgcc ccctaccgcg cctactacgg caaggaggcc
     2161 atgaagtacg ccgcccgcgc ctacatggag gaggcccagt ggcgcgagca gaagcacaag
     2221 cccaccacca aggagtacat gaagctggcg accaagacct gcggctacat caccctgatc
     2281 atcctgtcct gcctgggcgt ggaggagggc atcgtgacca aggaggcgtt cgactgggtg
     2341 ttcagccgcc cgccgttcat cgaggcgacc ctgatcatcg cccgcctggt caacgacatc
     2401 accggccacg agttcgagaa gaagcgcgag cacgtgcgca ccgccgtgga gtgctacatg
     2461 gaggtgagtc gacgagcaag cccggcggat caggcagcgt gcttgcagat ttgacttgca
     2521 acgcccgcat tgtgtcgacg aaggcttttg gctcctctgt cgctgtctca agcagcatct
     2581 aaccctgcgt cgccgtttcc atttgcagga gcacaaggtg ggcaagcagg aggtggtgtc
     2641 cgagttctac aaccagatgg agagcgcctg gaaggacatc aacgagggct tcctgcgccc
     2701 cgtggagttc cccatccccc tgctgtacct gatcctgaac agcgtgcgca ccctggaggt
     2761 gatctacaag gagggcgaca gctacaccca cgtgggcccg gccatgcaga acatcatcaa
     2821 gcagctgtac ctgcaccccg tgccctacgg cagcggcaga tccgagctgt acgcccagag
     2881 cgtgggcgtg ggcgccgcca gccgccccct ggccaacttc cacccctgcg tgtggggcga
     2941 caagttcatc gtgtacaacc cccagagctg ccaggccggc gagcgcgagg aggccgagga
     3001 gctgaaggtg gagctgaagc gcgagctgaa ggaggccagc gacaactaca tgcgccagct
     3061 gaagatggtg gacgccatcc agcgcctggg catcgactac ctgttcgtgg aggacgtgga
     3121 cgaggccctg aagaacctgt tcgagatgtt cgacgccttc tgcaagaaca accacgacat
     3181 gcacgccacc gccctgagct tccgcctgct gcgccagcac ggctaccgcg tgtcctgcga
     3241 ggtgttcgag aagttcaagg acggcaagga cggcttcaag gtgcccaacg aggacggcgc
     3301 ggtgagtcga cgagcaagcc cggcggatca ggcagcgtgc ttgcagattt gacttgcaac
     3361 gcccgcattg tgtcgacgaa ggcttttggc tcctctgtcg ctgtctcaag cagcatctaa
     3421 ccctgcgtcg ccgtttccat ttgcaggtgg cggtgctgga gttcttcgag gccacccacc
     3481 tgcgcgtgca cggcgaggac gtgctggaca acgccttcga cttcacccgc aactacctgg
     3541 agagcgtgta cgccaccctg aacgacccca ccgccaagca ggtgcacaac gcgctgaacg
     3601 agttctcctt ccgccgcggc ctgccccgcg tggaggcccg caagtacatc agcatctacg
     3661 agcagtacgc cagccaccac aagggcctgc tgaagctggc caagctggac ttcaacctgg
     3721 tgcaggcgct gcaccgccgc gagctgtccg aggacagccg ctggtggaag accctgcagg
     3781 tgcccaccaa gctgagcttc gtgcgcgacc gcctggtgga gagctacttc tgggccagcg
     3841 gcagctactt cgagcccaac tacagcgtgg cccgcatgat cctggcgaag ggcctggccg
     3901 tgctgagcct gatggacgac gtgtacgacg cctacggcac cttcgaggtg agtcgacgag
     3961 caagcccggc ggatcaggca gcgtgcttgc agatttgact tgcaacgccc gcattgtgtc
     4021 gacgaaggct tttggctcct ctgtcgctgt ctcaagcagc atctaaccct gcgtcgccgt
     4081 ttccatttgc aggagctgca gatgttcacc gacgcgatcg agcgctggga cgccagctgc
     4141 ctggacaagc tgcccgacta catgaagatc gtgtacaagg ccctgctgga cgtgttcgag
     4201 gaggtggacg aggagctgat caagctgggc gccccctacc gcgcctacta cggcaaggag
     4261 gccatgaagt acgccgcccg cgcctacatg gaggaggccc agtggcgcga gcagaagcac
     4321 aagcccacca ccaaggagta catgaagctg gcgaccaaga cctgcggcta catcaccctg
     4381 atcatcctgt cctgcctggg cgtggaggag ggcatcgtga ccaaggaggc gttcgactgg
     4441 gtgttcagcc gcccgccgtt catcgaggcg accctgatca tcgcccgcct ggtcaacgac
     4501 atcaccggcc acgagttcga gaagaagcgc gagcacgtgc gcaccgccgt ggagtgctac
     4561 atggaggtga gtcgacgagc aagcccggcg gatcaggcag cgtgcttgca gatttgactt
     4621 gcaacgcccg cattgtgtcg acgaaggctt ttggctcctc tgtcgctgtc tcaagcagca
     4681 tctaaccctg cgtcgccgtt tccatttgca ggagcacaag gtgggcaagc aggaggtggt
     4741 gtccgagttc tacaaccaga tggagagcgc ctggaaggac atcaacgagg gcttcctgcg
     4801 ccccgtggag ttccccatcc ccctgctgta cctgatcctg aacagcgtgc gcaccctgga
     4861 ggtgatctac aaggagggcg acagctacac ccacgtgggc ccggccatgc agaacatcat
     4921 caagcagctg tacctgcacc ccgtgcccta cggcagcggc agatccgagc tgtacgccca
     4981 gagcgtgggc gtgggcgccg ccagccgccc cctggccaac ttccacccct gcgtgtgggg
     5041 cgacaagttc atcgtgtaca acccccagag ctgccaggcc ggcgagcgcg aggaggccga
     5101 ggagctgaag gtggagctga agcgcgagct gaaggaggcc agcgacaact acatgcgcca
     5161 gctgaagatg gtggacgcca tccagcgcct gggcatcgac tacctgttcg tggaggacgt
     5221 ggacgaggcc ctgaagaacc tgttcgagat gttcgacgcc ttctgcaaga acaaccacga
     5281 catgcacgcc accgccctga gcttccgcct gctgcgccag cacggctacc gcgtgtcctg
     5341 cgaggtgttc gagaagttca aggacggcaa ggacggcttc aaggtgccca acgaggacgg
     5401 cgcggtgagt cgacgagcaa gcccggcgga tcaggcagcg tgcttgcaga tttgacttgc
     5461 aacgcccgca ttgtgtcgac gaaggctttt ggctcctctg tcgctgtctc aagcagcatc
     5521 taaccctgcg tcgccgtttc catttgcagg tggcggtgct ggagttcttc gaggccaccc
     5581 acctgcgcgt gcacggcgag gacgtgctgg acaacgcctt cgacttcacc cgcaactacc
     5641 tggagagcgt gtacgccacc ctgaacgacc ccaccgccaa gcaggtgcac aacgcgctga
     5701 acgagttctc cttccgccgc ggcctgcccc gcgtggaggc ccgcaagtac atcagcatct
     5761 acgagcagta cgccagccac cacaagggcc tgctgaagct ggccaagctg gacttcaacc
     5821 tggtgcaggc gctgcaccgc cgcgagctgt ccgaggacag ccgctggtgg aagaccctgc
     5881 aggtgcccac caagctgagc ttcgtgcgcg accgcctggt ggagagctac ttctgggcca
     5941 gcggcagcta cttcgagccc aactacagcg tggcccgcat gatcctggcg aagggcctgg
     6001 ccgtgctgag cctgatggac gacgtgtacg acgcctacgg caccttcgag gtgagtcgac
     6061 gagcaagccc ggcggatcag gcagcgtgct tgcagatttg acttgcaacg cccgcattgt
     6121 gtcgacgaag gcttttggct cctctgtcgc tgtctcaagc agcatctaac cctgcgtcgc
     6181 cgtttccatt tgcaggagct gcagatgttc accgacgcga tcgagcgctg ggacgccagc
     6241 tgcctggaca agctgcccga ctacatgaag atcgtgtaca aggccctgct ggacgtgttc
     6301 gaggaggtgg acgaggagct gatcaagctg ggcgccccct accgcgccta ctacggcaag
     6361 gaggccatga agtacgccgc ccgcgcctac atggaggagg cccagtggcg cgagcagaag
     6421 cacaagccca ccaccaagga gtacatgaag ctggcgacca agacctgcgg ctacatcacc
     6481 ctgatcatcc tgtcctgcct gggcgtggag gagggcatcg tgaccaagga ggcgttcgac
     6541 tgggtgttca gccgcccgcc gttcatcgag gcgaccctga tcatcgcccg cctggtcaac
     6601 gacatcaccg gccacgagtt cgagaagaag cgcgagcacg tgcgcaccgc cgtggagtgc
     6661 tacatggagg tgagtcgacg agcaagcccg gcggatcagg cagcgtgctt gcagatttga
     6721 cttgcaacgc ccgcattgtg tcgacgaagg cttttggctc ctctgtcgct gtctcaagca
     6781 gcatctaacc ctgcgtcgcc gtttccattt gcaggagcac aaggtgggca agcaggaggt
     6841 ggtgtccgag ttctacaacc agatggagag cgcctggaag gacatcaacg agggcttcct
     6901 gcgccccgtg gagttcccca tccccctgct gtacctgatc ctgaacagcg tgcgcaccct
     6961 ggaggtgatc tacaaggagg gcgacagcta cacccacgtg ggcccggcca tgcagaacat
     7021 catcaagcag ctgtacctgc accccgtgcc ctacggcagc ggcagatccg agctgtacgc
     7081 ccagagcgtg ggcgtgggcg ccgccagccg ccccctggcc aacttccacc cctgcgtgtg
     7141 gggcgacaag ttcatcgtgt acaaccccca gagctgccag gccggcgagc gcgaggaggc
     7201 cgaggagctg aaggtggagc tgaagcgcga gctgaaggag gccagcgaca actacatgcg
     7261 ccagctgaag atggtggacg ccatccagcg cctgggcatc gactacctgt tcgtggagga
     7321 cgtggacgag gccctgaaga acctgttcga gatgttcgac gccttctgca agaacaacca
     7381 cgacatgcac gccaccgccc tgagcttccg cctgctgcgc cagcacggct accgcgtgtc
     7441 ctgcgaggtg ttcgagaagt tcaaggacgg caaggacggc ttcaaggtgc ccaacgagga
     7501 cggcgcggtg agtcgacgag caagcccggc ggatcaggca gcgtgcttgc agatttgact
     7561 tgcaacgccc gcattgtgtc gacgaaggct tttggctcct ctgtcgctgt ctcaagcagc
     7621 atctaaccct gcgtcgccgt ttccatttgc aggtggcggt gctggagttc ttcgaggcca
     7681 cccacctgcg cgtgcacggc gaggacgtgc tggacaacgc cttcgacttc acccgcaact
     7741 acctggagag cgtgtacgcc accctgaacg accccaccgc caagcaggtg cacaacgcgc
     7801 tgaacgagtt ctccttccgc cgcggcctgc cccgcgtgga ggcccgcaag tacatcagca
     7861 tctacgagca gtacgccagc caccacaagg gcctgctgaa gctggccaag ctggacttca
     7921 acctggtgca ggcgctgcac cgccgcgagc tgtccgagga cagccgctgg tggaagaccc
     7981 tgcaggtgcc caccaagctg agcttcgtgc gcgaccgcct ggtggagagc tacttctggg
     8041 ccagcggcag ctacttcgag cccaactaca gcgtggcccg catgatcctg gcgaagggcc
     8101 tggccgtgct gagcctgatg gacgacgtgt acgacgccta cggcaccttc gaggtgagtc
     8161 gacgagcaag cccggcggat caggcagcgt gcttgcagat ttgacttgca acgcccgcat
     8221 tgtgtcgacg aaggcttttg gctcctctgt cgctgtctca agcagcatct aaccctgcgt
     8281 cgccgtttcc atttgcagga gctgcagatg ttcaccgacg cgatcgagcg ctgggacgcc
     8341 agctgcctgg acaagctgcc cgactacatg aagatcgtgt acaaggccct gctggacgtg
     8401 ttcgaggagg tggacgagga gctgatcaag ctgggcgccc cctaccgcgc ctactacggc
     8461 aaggaggcca tgaagtacgc cgcccgcgcc tacatggagg aggcccagtg gcgcgagcag
     8521 aagcacaagc ccaccaccaa ggagtacatg aagctggcga ccaagacctg cggctacatc
     8581 accctgatca tcctgtcctg cctgggcgtg gaggagggca tcgtgaccaa ggaggcgttc
     8641 gactgggtgt tcagccgccc gccgttcatc gaggcgaccc tgatcatcgc ccgcctggtc
     8701 aacgacatca ccggccacga gttcgagaag aagcgcgagc acgtgcgcac cgccgtggag
     8761 tgctacatgg aggtgagtcg acgagcaagc ccggcggatc aggcagcgtg cttgcagatt
     8821 tgacttgcaa cgcccgcatt gtgtcgacga aggcttttgg ctcctctgtc gctgtctcaa
     8881 gcagcatcta accctgcgtc gccgtttcca tttgcaggag cacaaggtgg gcaagcagga
     8941 ggtggtgtcc gagttctaca accagatgga gagcgcctgg aaggacatca acgagggctt
     9001 cctgcgcccc gtggagttcc ccatccccct gctgtacctg atcctgaaca gcgtgcgcac
     9061 cctggaggtg atctacaagg agggcgacag ctacacccac gtgggcccgg ccatgcagaa
     9121 catcatcaag cagctgtacc tgcaccccgt gccctacggc agcggcagat ctgacgtcgg
     9181 cagcggcagc ggcagcgtga gcaagggcga ggagctgttc accggcgtgg tgcccatcct
     9241 ggtggagctg gacggcgacg tgaacggcca caagttcagc gtgagcggcg agggcgaggg
     9301 cgacgccacc tacggcaagc tgaccctgaa gctgatctgc accaccggca agctgcccgt
     9361 gccctggccc accctggtga ccaccctggg ctacggtgag tcgacgagca agcccggcgg
     9421 atcaggcagc gtgcttgcag atttgacttg caacgcccgc attgtgtcga cgaaggcttt
     9481 tggctcctct gtcgctgtct caagcagcat ctaaccctgc gtcgccgttt ccatttgcag
     9541 gcctgcagtg cttcgcccgc taccccgacc acatgaagca gcacgacttc ttcaagagcg
     9601 ccatgcccga gggctacgtg caggagcgca ccatcttctt caaggacgac ggtaactaca
     9661 agacccgcgc cgaggtgaag ttcgagggcg acaccctggt gaaccgcatc gagctgaagg
     9721 gcatcgactt caaggaggac ggcaacatcc tgggccacaa gctggagtac aactacaaca
     9781 gccacaacgt gtacatcacc gccgacaagc agaagaacgg catcaaggcc aacttcaaga
     9841 tccgccacaa catcgaggtg agtcgacgag caagcccggc ggatcaggca gcgtgcttgc
     9901 agatttgact tgcaacgccc gcattgtgtc gacgaaggct tttggctcct ctgtcgctgt
     9961 ctcaagcagc atctaaccct gcgtcgccgt ttccatttgc aggacggcgg cgtgcagctg
    10021 gccgaccact accagcagaa cacccccatc ggcgacggcc ccgtgctgct gcccgacaac
    10081 cactacctga gctaccagag caagctgagc aaggacccca acgagaagcg cgaccacatg
    10141 gtgctgctgg agttcgtgac cgccgccggc atcaccctgg gcatggacga gctgtacaag
    10201 ggcagcggca gcggcagcga tatcgaattc ggcagcggca gcggctcagg tgagcttgcg
    10261 gggttgcgag caacactcca gcaacgaaca gtgcccaagt caggaatctg cagtcagcct
    10321 gggctttcgg cggctttttc ttgggcaaac agcttgcact catgccagcg cggcttgtcc
    10381 agcctcactt gagctttcca gctgctacca gccgggctat acgacagcga cagagccata
    10441 gcgtggaatc acttatttgg gttgccgaag tagcggtcgg agcgtgagtt cttggtcaag
    10501 ccgcccctta tccggttcct gtccgtgtct ttgtccctcg ttcacccttc gcggcaccct
    10561 tcatcccctt gcttgcaggt tggagccacc cgcagttcga gaagtaaccg ctccgtgtaa
    10621 atggaggcgc tcgttgatct gagccttgcc ccctgacgaa cggcggtgga tggaagatac
    10681 tgctctcaag tgctgaagcg gtagcttagc tccccgtttc gtgctgatca gtctttttca
    10741 acacgtaaaa agcggaggag ttttgcaatt ttgttggttg taacgatcct ccgttgattt
    10801 tggcctcttt ctccatgggc gggctgggcg tatttgaagc gactagtacg cgtgctgagg
    10861 cttgacatga ttggtgcgta tgtttgtatg aagctacagg actgatttgg cgggctatga
    10921 gggcggggga agctctggaa gggccgcgat ggggcgcgcg gcgtccagaa ggcgccatac
    10981 ggcccgctgg cggcacccat ccggtataaa agcccgcgac cccgaacggt gacctccact
    11041 ttcagcgaca aacgagcact tatacatacg cgactattct gccgctatac ataaccactc
    11101 agctagctta agatcccatc cctagggcat gccgggcgcg ccagaaggag cgcagccaaa
    11161 ccaggatgat gtttgatggg gtatttgagc acttgcaacc cttatccgga agccccctgg
    11221 cccacaaagg ctaggcgcca atgcaagcag ttcgcatgca gcccctggag cggtgccctc
    11281 ctgataaacc ggccaggggg cctatgttct ttactttttt acaagagaag tcactcaaca
    11341 tcttaaaatg gccaggtgag tcgacgagca agcccggcgg atcaggcagc gtgcttgcag
    11401 atttgacttg caacgcccgc attgtgtcga cgaaggcttt tggctcctct gtcgctgtct
    11461 caagcagcat ctaaccctgc gtcgccgttt ccatttgcag gaagcttact ccgccctccc
    11521 cggtgctgaa gaatttcgaa gcatggacga tgcgttgcgt gcactgcggg gtcggtatcc
    11581 cggttgtgag tgggttgttg tggaggatgg ggcctcgggg gctggtgttt atcggcttcg
    11641 gggtggtggg cgggagttgt ttgtcaaggt ggcagctctg ggggccgggg tgggcttgtt
    11701 gggtgaggct gagcggctgg tgtggttggc ggaggtgggg attcccgtac ctcgtgttgt
    11761 ggagggtggt ggggacgaga gggtcgcctg gttggtcacc gaagcggttc cggggcgtcc
    11821 ggccagtgcg cggtggccgc gggagcagcg gctggacgtg gcggtggcgc tcgcggggct
    11881 cgctcgttcg ctgcacgcgc tggactggga gcggtgtccg ttcgatcgca gtctcgcggt
    11941 gacggtgccg caggcggccc gtgctgtcgc tgaagggagc gtcgacttgg aggatctgga
    12001 cgaggagcgg aaggggtggt cgggggagcg gcttctcgcc gagctggagc ggactcggcc
    12061 tgcggacgag gatctggcgg tttgccacgg tgacctgtgc ccggacaacg tgctgctcga
    12121 ccctcgtacc tgcgaggtga ccgggctgat cgacgtgggg cgggtcggcc gtgcggaccg
    12181 gcactccgat ctcgcgctgg tgctgcgcga gctggcccac gaggaggacc cgtggttcgg
    12241 gccggagtgt tccgcggcgt tcctgcggga gtacgggcgc gggtgggatg gggcggtatc
    12301 ggaggaaaag ctggcgtttt accggctgtt ggacgagttc ttctgactcg agtgaccgct
    12361 ccgtgtaaat ggaggcgctc gttgatctga gccttgcccc ctgacgaacg gcggtggatg
    12421 gaagatactg ctctcaagtg ctgaagcggt agcttagctc cccgtttcgt gctgatcagt
    12481 ctttttcaac acgtaaaaag cggaggagtt ttgcaatttt gttggttgta acgatcctcc
    12541 gttgattttg gcctctttct ccatgggcgg gctgggcgta tttgaagcgg acccggtacc
    12601 cagcttttgt tccctttagt gagggttaat tgcgcgcttg gcgtaatcat ggtcatagct
    12661 gtttcctgtg tgaaattgtt atccgctcac aattccacac aacatacgag ccggaagcat
    12721 aaagtgtaaa gcctggggtg cctaatgagt gagctaactc acattaattg cgttgcgctc
    12781 actgcccgct ttccagtcgg gaaacctgtc gtgccagctg cattaatgaa tcggccaacg
    12841 cgcggggaga ggcggtttgc gtattgggcg ctcttccgct tcctcgctca ctgactcgct
    12901 gcgctcggtc gttcggctgc ggcgagcggt atcagctcac tcaaaggcgg taatacggtt
    12961 atccacagaa tcaggggata acgcaggaaa gaacatgtga gcaaaaggcc agcaaaaggc
    13021 caggaaccgt aaaaaggccg cgttgctggc gtttttccat aggctccgcc cccctgacga
    13081 gcatcacaaa aatcgacgct caagtcagag gtggcgaaac ccgacaggac tataaagata
    13141 ccaggcgttt ccccctggaa gctccctcgt gcgctctcct gttccgaccc tgccgcttac
    13201 cggatacctg tccgcctttc tcccttcggg aagcgtggcg ctttctcata gctcacgctg
    13261 taggtatctc agttcggtgt aggtcgttcg ctccaagctg ggctgtgtgc acgaaccccc
    13321 cgttcagccc gaccgctgcg ccttatccgg taactatcgt cttgagtcca acccggtaag
    13381 acacgactta tcgccactgg cagcagccac tggtaacagg attagcagag cgaggtatgt
    13441 aggcggtgct acagagttct tgaagtggtg gcctaactac ggctacacta gaaggacagt
    13501 atttggtatc tgcgctctgc tgaagccagt taccttcgga aaaagagttg gtagctcttg
    13561 atccggcaaa caaaccaccg ctggtagcgg tggttttttt gtttgcaagc agcagattac
    13621 gcgcagaaaa aaaggatctc aagaagatcc tttgatcttt tctacggggt ctgacgctca
    13681 gtggaacgaa aactcacgtt aagggatttt ggtcatgaga ttatcaaaaa ggatcttcac
    13741 ctagatcctt ttaaattaaa aatgaagttt taaatcaatc taaagtatat atgagtaaac
    13801 ttggtctgac agttaccaat gcttaatcag tgaggcacct atctcagcga tctgtctatt
    13861 tcgttcatcc atagttgcct gactccccgt cgtgtagata actacgatac gggagggctt
    13921 accatctggc cccagtgctg caatgatacc gcgagaccca cgctcaccgg ctccagattt
    13981 atcagcaata aaccagccag ccggaagggc cgagcgcaga agtggtcctg caactttatc
    14041 cgcctccatc cagtctatta attgttgccg ggaagctaga gtaagtagtt cgccagttaa
    14101 tagtttgcgc aacgttgttg ccattgctac aggcatcgtg gtgtcacgct cgtcgtttgg
    14161 tatggcttca ttcagctccg gttcccaacg atcaaggcga gttacatgat cccccatgtt
    14221 gtgcaaaaaa gcggttagct ccttcggtcc tccgatcgtt gtcagaagta agttggccgc
    14281 agtgttatca ctcatggtta tggcagcact gcataattct cttactgtca tgccatccgt
    14341 aagatgcttt tctgtgactg gtgagtactc aaccaagtca ttctgagaat agtgtatgcg
    14401 gcgaccgagt tgctcttgcc cggcgtcaat acgggataat accgcgccac atagcagaac
    14461 tttaaaagtg ctcatcattg gaaaacgttc ttcggggcga aaactctcaa ggatcttacc
    14521 gctgttgaga tccagttcga tgtaacccac tcgtgcaccc aactgatctt cagcatcttt
    14581 tactttcacc agcgtttctg ggtgagcaaa aacaggaagg caaaatgccg caaaaaaggg
    14641 aataagggcg acacggaaat gttgaatact catactcttc ctttttcaat attattgaag
    14701 catttatcag ggttattgtc tcatgagcgg atacatattt gaatgtattt agaaaaataa
    14761 acaaataggg gttccgcgca catttccccg aaaagtgcca cactaaattg taagcgttaa
    14821 tattttgtta aaattcgcgt taaatttttg ttaaatcagc tcatttttta accaataggc
    14881 cgaaatcggc aaaatccctt ataaatcaaa agaatagacc gagatagggt tgagtgttgt
    14941 tccagtttgg aacaagagtc cactattaaa gaacgtggac tccaacgtca aagggcgaaa
    15001 aaccgtctat cagggcgatg gcccactacg tgaaccatca ccctaatcaa gttttttggg
    15061 gtcgaggtgc cgtaaagcac taaatcggaa ccctaaaggg agcccccgat ttagagcttg
    15121 acggggaaag ccggcgaacg tggcgagaaa ggaagggaag aaagcgaaag gagcgggcgc
    15181 tagggcgctg gcaagtgtag cggtcacgct gcgcgtaacc accacacccg ccgcgcttaa
    15241 tgcgccgcta cagggcgcgt cccattcgcc attcaggctg cgcaactgtt gggaagggcg
    15301 atcggtgcgg gcctcttcgc tattacgcca gctggcgaaa gggggatgtg ctgcaaggcg
    15361 attaagttgg gtaacgccag ggttttccca gtcacgacgt tgtaaaacga cggccagtga
    15421 gcgcgcgtaa tacgactcac tatagggcga attggagctc caccgcggtg gcggccg
//
```
