## Supplemental Table 1 - Plasmids used in this work for "Combinatorial engineering for photoautotrophic production of recombinant products from the green microalga *Chlamydomonas reinhardtii*"

**Table 1: Genetic constructs used in this study**

| **Construct name** | **Selection in *C. reinhardtii*** | **Plasmid size (bp)** | | **CDS (bp)** | | **Introns** | | **Protein length (aa)** | | **Predicted molecular weight (kDa)** | | **Reference** |
| --- | --- | --- | --- | --- | --- | --- | --- | --- | --- | --- | --- | --- |
| pPO3 - chloroplast | phosphite | 9350 | | 1014 | | n/a | | 338 | | 36.6 | | Changko et al. (2020) |
| pMN24 (Nit1) | nitrate | 17622 | | 2649 | | 15 native introns | | 882 | | 97.8 | | Fernández et al. (1989) |
| pMN68 (Nit2) | nitrate | 15499 | | 2265 | | 6 native introns | | 754 | | 76.5 | | Schnell & Lefebvre (1993) |
|  |  | **Gene length with introns from start codon to stop (bp)** | |  | | **RBCS2 i1** | |  | | ***Pc*PS fusions (kDa)** | |  |
| pOpt2_mVenus_Paro | paromomycin | 1287 | 813 | | 1 | | 270 | | 30.6 | | Wichmann et al. (2018) | |
| pOptN_1X*Pc*PS_YFP | paromomycin | 3547 | 2493 | | 6 | | 830 | | 94.6 | | This work | |
| pOptN_2X*Pc*PS_YFP | paromomycin | 5650 | 4161 | | 9 | | 1386 | | 159.1 | | This work | |
| pOptN_3X*Pc*PS_YFP | paromomycin | 7753 | 5829 | | 12 | | 1942 | | 223.6 | | This work | |
| pOptN_4X*Pc*PS_YFP | paromomycin | 9856 | 7497 | | 15 | | 2498 | | 288.1 | | This work | |
| *pOpt2_cCA_g*Luc_SQS_k.d._Spec | spectinomycin | 1104 | 630 | | 1 | | 209 | | - | | Wichmann et al. (2018) | |
|  | **UniProt ID** |  |  | |  | |  | |  | |  | |
| *Pc*Ps alone | Q49SP3 | 2097 | 1662 | | 3 | | 554 | | 64.3 | | Lauersen et al. (2016) | |
